# Arabidopsis Proteome and the Mass Spectral Assay Library

**DOI:** 10.1101/665547

**Authors:** Huoming Zhang, Pei Liu, Tiannan Guo, Huayan Zhao, Dalila Bensaddek, Ruedi Aebersold, Liming Xiong

## Abstract

*Arabidopsis* is an important model organism and the first plant with its genome sequenced. Knowledge from studying this species has either direct or indirect applications to agriculture and human health. Quantitative proteomics by data-independent acquisition (SWATH/DIA-MS) was recently developed and considered as a high-throughput targeted-like approach for accurate proteome quantitation. In this approach, a high-quality and comprehensive library is a prerequisite. Here, we generated a protein expression atlas of 10 organs of *Arabidopsis* and created a library consisting of 15,514 protein groups, 187,265 unique peptide sequences, and 278,278 precursors. The identified protein groups correspond to ~56.5% of the predicted proteome. Further proteogenomics analysis identified 28 novel proteins. We subsequently applied DIA-mass spectrometry using this library to quantify the effect of abscisic acid on *Arabidopsis.* We were able to recover 8,793 protein groups with 1,787 of them being differentially expressed which includes 65 proteins known to respond to abscisic acid stress. Mass spectrometry data are available via ProteomeXchange with identifier PXD012710 for data-dependent acquisition and PXD014032 for DIA analyses.

|  |  |
| --- | --- |
| Design Type(s) | reference design • replicate design • protein expression profiling • quality control testing design |
| Measurement Type(s) | protein expression profiling |
| Technology Type(s) | mass spectrometry assay |
| Factor Type(s) | purification • proteolysis |
| Sample Characteristic(s) | *Arabidopsis thaliana* • leaf • flower • stem • seed • root • root cell culture |

## Background & Summary

The *Arabidopsis thaliana* is a flowering plant with a short but complex life cycle. It has a relatively small genome size with low repetitive content (10%). These features make it an ideal organism for laboratory research. Knowledge from studying this species has either direct or indirect applications to agriculture and human health^1^. *Arabidopsis* therefore became the first plant to have its genome completely sequenced and annotated under “The *Arabidopsis* Genome Initiative 2000”^2^, which greatly promoted whole genome sequencing and global transcriptome analysis using next generation sequencing technology^3^. Proteins bridge genetic information and phenotypes. However, the protein abundances generally exhibit poor correlation with genetic variations^4^, necessitating direct study of proteins under different biological conditions.

Proteomics, defined as the study of all proteins in any given sample, has advanced at a fast rate in the last decade, especially in quantitative proteomics that has been widely used for both discovery and targeted analyses. Three commonly used discovery quantitative proteomics strategies are chemical labeling such as isobaric tags for relative and absolute quantitation (iTRAQ^5^) and tandem mass tags (TMT^6^), metabolic labeling (SILAC^7^, ^15^N labeling^8^ etc.) and label-free^9,10^ approaches. These methods are generally high-throughput and provide in-depth coverage suitable for system-wide analyses. However, in order to maximize proteome coverage, it is necessary to incorporate a sample prefractionation step prior to liquid chromatography-mass spectrometry (LC-MS) analysis. In addition to substantially increasing both data acquisition and analysis times, this leads to a reduction in the reproducibility of measurements and the quantitative accuracy especially in extended label-free experiments. Multiplexed analyses such as TMT/iTRAQ reduce the analysis time, but suffer from ratio compressions which in turn impacts protein quantification^11,12^. On the other hand, the targeted proteomics (S/MRM^13^: selected/multiple reaction monitoring; PRM^14^: parallel reaction monitoring) provides higher sensitivity and reproducibility but it has limited use for proteome-wide survey.

Recently, a relatively new technique termed SWATH/DIA-MS (SWATH^15^: sequential window acquisition of all theoretical mass spectra; DIA: data-independent acquisition) was developed to complement discovery and targeted proteomics. In this approach, the precursors across the mass range of interest (e.g. 400-1200 Da) are sequentially and cyclically isolated using a wide mass window (typically 25 Da) and subjected to fragmentation. Thus, the spectra of all ions including low abundance ions from a sample are acquired in an unbiased fashion. Subsequently targeted extraction of fragmented spectra can be performed by comparing the acquired spectral data with a pre-constructed ion libraries consisting of pairs of ion spectra and their accurate retention time to identify and quantify proteins. The advantages of this approach include its ability for proteome-wide quantitation with high consistency and accuracy, acquiring data via a hypothesis-free approach^16^, and its particular suitability for studying a large number of sample in a reproducible way^17^.

The sensitivity in SWATH/DIA-MS relies heavily on a high-quality and comprehensive assay library. In this study, we used two common mass spectrometry platforms (Orbitrap and TripleTOF) to analyze 10 different organs of *Arabidopsis* after extensive offline sample fractionation. We acquired a total of 836 mass spectrometry raw files including 463 from Orbitrap Fusion and 373 from TripleTOF5600 plus. As a result, we constructed a spectral library containing more than 19,000 proteins (>15,000 protein groups), accounting for approximately 56.5% of the predicted *Arabidopsis* proteome. The usefulness of this library has been clearly illustrated in the subsequent DIA-MS analysis of *Arabidopsis* leaves treated with the plant hormone abscisic acid (ABA).

## Methods

### Plant Materials and Growth Conditions

*Arabidopsis thaliana* ecotype Col-0 was used for this study. Seeds were obtained from the *Arabidopsis* Biological Resource Center (ABRC) and the European *Arabidopsis* Stock Centre. They were surface sterilized with 75% ethanol and 0.1% Triton X-100 for 10 min, washed with 95% ethanol twice (2 min each), and then planted on growth media containing half strength Murashige and Skoog (½ x MS) salt and 1% (w/v) sucrose, and solidified with 0.8% agar. The plates were kept at 4 °C in the dark for 2 d and then moved to a growth chamber (CU36-L5, Percival Scientific) at 21 °C under a photoperiod of 16 h light and 8 h darkness for germination and growth. Cotyledons and roots were collected following growth for 6 days and 20 days, respectively. For preparation of leaves (rosette leaves and cauline leaves), stems, flowers (buds and open flowers) and siliques (green siliques), seedlings grown for one week on the plate were transferred to soil in a greenhouse room under the same growth conditions as in the growth chamber for an additional 5 week period, except rosette leaves which were grown for 3 weeks. Seeds were harvested when siliques turned yellow or brown. For the study of abscisic acid effect on *Arabidopsis,* rosette leaves were sprayed with or without 100 μM of abscisic acid and harvested at 2 h, 24 h and 72 h post treatment. All materials were collected and immediately frozen in liquid nitrogen and stored at −80 °C until use.

### *Arabidopsis thaliana* root cell suspension culture

Cells isolated from roots of *Arabidopsis thaliana* were grown in Gamborg’s B5 basal salt mixture (Sigma-Aldrich) with 2,4-dichlorophenoxyacetic acid (2,4-D; 1 mg mL^−1^) and kinetin (0.05 μg mL^−1^) in sterile flask as described^18^. Briefly, cells were grown in a growth chamber (Innova^®^ 43, New Brunswick Scientific Co., NJ) with shaking at 120 rpm, and subcultured every 10 days. Photosynthetic light of the growth chamber was set for 16 h light/8 h dark cycles at 21°C. The cells were harvested by draining off the media using Stericup^®^ filter unit (Millipore, Billerica, MA), immediately flash frozen in liquid nitrogen and stored at −80°C until use.

### Protein extraction and digestion

All plant materials were ground in liquid nitrogen with a prechilled mortar and a pestle. The fine powder was resuspended with the extraction buffer (50 mm Tris, pH 8, 8 M urea, and 0.5% SDS) supplemented with protease inhibitor (Roche Diagnostics GmbH, Mannheim, Germany), and homogenized with a Dounce homogenizer. In order to extract more proteins, the crude homogenate was further subjected to 30 cyclic high/low pressurization (50 sec of 35, 000 PSI and 10 sec of ambient pressure) using a pressure cycling technology (Barocycle, PressureBioSciences, MA). The extracts were then centrifuged at 10,000 *g* for 5 min at 4°C. The proteins in the supernatant were purified using methanol/chloroform precipitation and dried under vacuum. The dried pellets were resuspended into the extraction buffer (50 mm Tris, pH 8, 8 M urea, and 0.5% SDS) with assistance of sonication and sonicated. The protein content was determined using a microBCA kit (Thermo Scientific). For library generation, approximately 200 μg of proteins were reduced, alkylated and digested with trypsin as described^19^. The digests were desalted with microcolumns packed with C18 and Poros oligo R3 materials prior to a shallow-gradient Strong Cation Exchange (SCX) fractionation. For protein quantitation of ABA-treated sample, approximately 10 μg of proteins were digested using FASP method^20^ prior to DIA-MS analysis.

### SCX Peptide fractionation

The peptides were reconstituted in 90 μL SCX buffer A (10 mM KH_2_PO_4_, 25% acetonitrile (ACN), pH 3.0) and loaded into the polySULFOETHYL A column (200 × 4.6 mm, 5 μm, 200 Å) (PolyLC, Columbia, MD) for SCX fractionation on Accela HPLC (Thermo Scientific). An increasing gradient of buffer B (10 mM KH_2_PO_4_, 500 mM KCl and 25% ACN, pH 3.0) was applied in a shallow-gradient elution protocol of total 100 min. The gradient consists of 100% buffer A for the initial 5 min, 0%-15% buffer B for 80 min, 30%-100% buffer B for 5 min, 100% buffer B for 5 min and 100% A for 5 min at a flow rate of 1 mL/min. The chromatography was monitored at 214 nm using diode array detector. After the pooling of some fractions based on the absorption intensity, a total of 30 fractions were obtained, desalted as described above and dried in the SpeedVac (Thermo Scientific).

### MS analysis using TripleTOF 5600+

The dried peptide mixture was redissolved into 0.1% formic acid (FA) and 3% ACN in water supplemented with indexed retention time (iRT) peptide standards according to the manual (Biognosys, Switzerland). They were then analyzed using a TripleTOF 5600 Plus MS (Sciex, USA) coupled with an UltiMate™ 3000 UHPLC (Thermo Scientific). Briefly, approximately 1.5 μg of peptide mixture was injected into a precolumn (Acclaim PepMap, 300 μm×5 mm, 5 μm particle size) and desalted for 15 min with 3% ACN and 0.1% FA in water at a flow rate of 5 μl/min. The peptides were eluted into an Acclaim PepMap100 C18 capillary column (75 μm I.D. X 25 cm, 3 μm particle sizes, 100 Å pore sizes) and separated with a 135-min gradient at constant 300 nL/min, at 40°C. The gradient was established using mobile phase A (0.1% FA in H_2_O) and mobile phase B (0.1% FA, 95% ACN in H_2_O): 2.1%-5.3% B for 5 min, 5.3%-10.5% for 15 min, 10.5%-21.1% for 70 min, 21.1%-31.6% B for 18 min, ramping from 31.6% to 94.7% B in 2 min, maintaining at 94.7% for 5 min, and 4.7% B for 15-min column conditioning. The sample was introduced into the TripleTOF MS through a Nanospray III source (Sciex, USA) with an electrospray potential of 2.2 kV. The ion source was set with an interface heater temperature of 150 °C, a curtain gas of 25 PSI, and a nebulizer gas of 6 PSI. The mass spectrometry (MS) was performed with information dependent acquisition (IDA). The mass range of survey scans was set to 350-1250 Da. The top 30 ions of high intensity higher than 1,000 counts per second and a charge-state of 2+ to 4+ were selected for collision-induced dissociation. A rolling collision energy option was applied. The maximum cycle time was fixed to 2 s and a maximum accumulation time for individual ions was set for 250 ms. Dynamic exclusion was set to 15 s with a 50 mDa mass tolerance.

### MS analysis using Orbitrap Fusion Lumos

In both data-dependent acquisition (DDA) and DIA analysis, an Orbitrap Fusion Lumos mass spectrometer (Thermo Scientific) was coupled with an UltiMate™ 3000 UHPLC (Thermo Scientific). The peptide injection and elution gradient were essentially the same as described above. The peptides were separated and introduced into the Orbitrap MS through an integrated Easy-Spray LC column ( 50 cm x 75 μm ID, PepMap C18, 2 μm particles, 100 Å pore size, Thermo Scientific) with an electrospray potential of 1.9 kV. The ion transfer tube temperature was set at 270°C. The MS parameters included application mode as standard for peptide, default charge state of 3 and the use of EASY-IC as internal mass calibration in both precursor ions (MS1) and fragment ions (MS2).

In DDA mode, a full MS scan (375-1400 m/z range) was acquired in the Orbitrap at a resolution of 120,000 (at 200 m/z) in a profile mode. The cycle time was 3 s between master scans, whereas the RF lens was set to 30%. A maximum ion accumulation time was 100 milliseconds and a target value was 4e5. MIPS (monoisotopic peak determination of peptide) was activated. The isolation window for ions was 1.6 m/z. The ions above an intensity threshold of 5 e4 and carrying charges from 2^+^ to 5^+^ were selected for fragmentation using higher energy collision dissociation (HCD) at 30% energy. They were dynamically excluded after 1 event for 10 s with a mass tolerance of 10 ppm. The injections for all available parallelizable time were activated. The spectral with a first mass fixed at 100 (m/z) was acquired in Orbitrap at a resolution of 30,000 in a centroid mode. A maximum injection time of 100 ms and a target value of 5 e4 were used.

For the DIA-MS analysis, quadrupole isolation window of 6 m/z was selected for the HCD fragmentation. The sample was gas-fractionated into precursor mass ranges among 400-550; 550-700 and 700-850 m/z respectively in each injection. The mass defect was 0.9995. The HCD collision energy was set at 30%. MS2 has a resolution of 30,000, scan range to 350-1500 m/z, a maximum ion accumulation time of 100 milliseconds, a target value of 1e6, and data type to centroid.

### Protein Identification

Mass spectrometry data (.raw file from Orbitrap Fusion and .wiff file from TripleTOF 5600+) were processed using the Maxquant software (version 1.5.3.30)^21^. The TAIR10 proteome database (35,386) and 262 common contaminant sequences were combined and used for the database search. Carbamidomethylation at cysteine residues was set as a fixed modification. The variable modifications included oxidation at methionine residues and N - terminal protein acetylation. The enzyme limits were set at full trypsin cleavage with a maximum of two missed cleavages was allowed. A positive peptide was required to contain a minimum of seven amino acids and a maximum of five modifications. The mass tolerances of the precursor ion of Orbitrap fusion data were set to 20 and 4.5 ppm for the first and main searches, respectively, whereas they were 0.007 and 0.0006 Da for the TripleTOF 5600 data. The mass tolerances of the fragment ion were set 20 ppm and 40 ppm for Orbitrap Fusion and TripleTOF 5600 data, respectively. The mass tolerances of the fragments were 20 ppm for HCD. The false discovery rates (FDRs) of peptide-spectral match (PSM), protein identification and site decoy fraction were all set to 0.01.

To identify novel proteins, the Maxquant results were loaded into Scaffold software (version 4.4, Proteome software Inc., Portland, OR). The unmatched spectra from Maxquant searches were exported. They were then searched against a proteome database constructed using six-frame translation of the TAIR9 genome. The novel identifications were manually verified to further minimize FDR. At least two good spectra were required for confirming the identification. Each protein was matched with the Araprot11 transcript (DNA) using blast program (tblastn) to extract detailed annotation information in TAIR (https://www.arabidopsis.org) such as genomics locus and gene model type.

### MS Spectral Library Generation and DIA data analysis

All the DDA MS data files were loaded into Spectronaut Pulsar X (version 12, Biognosys, Switzerland) for the library generation. The protein database was the combination of TAIR10 proteome sequence and the aforementioned novel identifications. The default settings for database match include: full trypsin cleavage, peptide length of between 7 and 52 amino acids and maximum missed cleavage of 2. Besides, lysine and arginine (KR) were used as special amino acids for decoy generation, and N-terminal methionine was removed during pre-processing of the protein database. Fixed modification was carbamidomethylation at cysteine and variable modifications were acetylation at protein N-terminal and oxidation at methionine. All FDRs were set as 0.01 for the peptide-spectrum match (PSM), peptide and protein. The used Biognosys default spectral library filters include amino acid length of ion more than 2, ion mass-to-charge between 300 and 1800 Da, and minimum relative intensity of 5%. The best 3-6 fragments per peptide were included in the library. The iRT calibration was required with minimum R-Square of 0.8.

DIA data were analyzed using Spectronaut software against the spectral libraries to identify and quantify peptides and proteins. The Biognosys default settings were applied for identification: excluding duplicate assay; generation decoy based on mutated method at 10% of library size; and estimation of FDRs using Q value as 0.01 for both precursors and proteins. The p-value was calculated by kernel-density estimator. Interference correction was activated and a minimum of 3 fragment ions and 2 precursor ions were kept for the quantitation. The area of extracted ion chromatogram (XIC) at MS2 level were used for quantitation. Peptide (stripped sequence) quantity was measured by the mean of 1-3 best precursors, and protein quantity was calculated accordingly by the mean of 1-3 best peptides. Local normalization strategy and q-value sparse selection were used for cross run normalization. Differential expression was determined by performing paired Student’s t-test. Proteins with a fold-change of higher than 1.5 and a q-value of less than 0.01 were considered as differentially expressed proteins.

### Bioinformatic analyses

The candidate proteins were submitted to the web-based platform of the Database for Annotation, Visualization and Integrated Discovery (DAVID; http://david.abcc.ncifcrf.gov) for Gene Ontology (GO) enrichment and pathway analysis^22^. The abscisic acid responding protein expression data was input into MultiExperimentView^23^ software (version 4.9) using two color array. Following with Figure of Merit (FOM) analysis, these proteins were partitioned into 4 groups using k-means clustering. Protein–protein interactions were predicted using the STRING database (http://string-db.org, version 11).

## Data Records

The mass spectrometry DDA proteomics data acquired using TripleTof 5600 plus and Orbitrap Fusion Lumos have been deposited to the ProteomeXchange Consortium via the PRIDE^24^ partner repository with the dataset identifier PXD012710 (Data Citation 1), whereas the mass spectrometry DIA proteomics data acquired using Orbitrap Fusion Lumos have been deposited at the same server with the dataset identifier PXD014032 (Data Citation 2).

## Technical Validation

### Experimental design

In SWATH/DIA-MS, the high quality and coverage of an assay library is the key for accurate quantitation of high number of proteins. The *Arabidopsis* is a complex organism in that they have specialized organs at different developmental stages, and its proteome undergoes dynamic changes in different tissues during development.

To build a comprehensive assay library, we collected 10 samples from four different organs of *Arabidopsis* (Figure 1a) including leaf, stem, flower and root. We improved the protein and peptide preparation protocols (Figure 1b): use of cryogenic grinding of tissues, Dounce homogenization and pressure cyclic treatment (PCT technology) sequentially to get better yields in protein amount and species; purifying proteins sample by methanol/chloroform which helped to remove majority of non-proteins (such as lipids and pigments), and desalting peptides using a cartridge containing both C18 and R3 material to minimize loss of hydrophilic peptides. Besides we fractionated sample extensively using an optimal 100-min shallow-gradient SCX and combined into final 30 fractions for LC-MS analysis by two MS platforms (Figure 1c).

**Figure 1.**
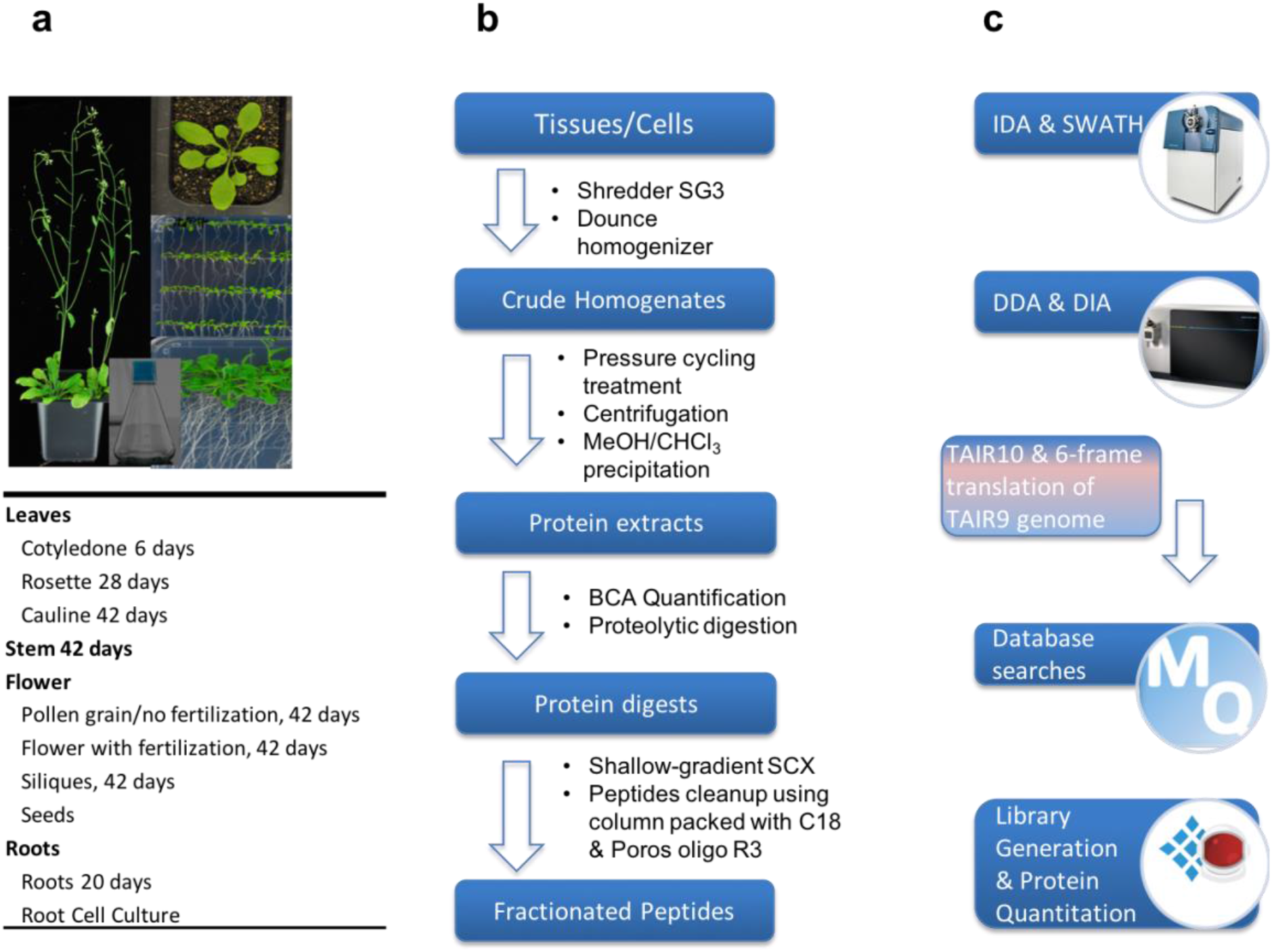
Schematic diagram of experimental workflow. a. Detailed sample information of the 10 *Arabidopsis* organs used for generation of spectral library; b. Optimal protein extraction and peptide purification procedures for in-depth coverage of *Arabidopsis* spectral library; c. The samples were analyzed using Orbitrap Fusion and TripleTOF mass spectrometer platforms for data dependent acquisition to construct of the spectral library. The DDA data were analyzed following by protein identification by Maxquant and generation of comprehensive library using Spectronaut Pulsar.

### Protein identification

Using the optimal workflow, we identified a high number of proteins from each organ (**Table 1**). The newer mass spectrometry platform Orbitrap Fusion generally gave ~30% more identification compared to the TripleTOF 5600 plus. In total, we identified more than 180,000 unique peptides from ~15,400 distinct protein groups, which were approximately 30% higher compared with two earlier genome-wide proteome analyses of Arabidoposis^25,26^, and accounted for ~55% of the total predicted proteome of *Arabidopsis*.

**Table 1.** Number of proteins identified from each organ using either TripleTOF 5600+ or Orbitrap Fusion Lumos.

| Organs | No. of Proteins identified |  |
| --- | --- | --- |
|  | TripleTOF 5600 | Orbitrap Fusion |
| Cotyledone | 7660 | 9308 |
| Rosette | 6034 | 7889 |
| Cauline | 6592 | 8851 |
| Stem | 6596 | 8946 |
| Pollen grain | 8591 | 11321 |
| Flower | 8737 | 11594 |
| Silique | 7950 | 10252 |
| Seeds | 4608 | 6739 |
| Root | 8665 | 10325 |
| Root cells | 8835 | 10640 |

Next, we constructed a new proteome database using six-frame translation of TAIR9 genome. The unmatched spectra from earlier searches against TAIR10 proteome database were searched again with the new database to identify potential novel genes or gene models as described^26^. As a result, we were able to identify a total of 42 novel proteins. Most novel findings were previously annotated as “transposable element gene”, “novel transcribed region” and “long noncoding RNA” as well as novel alternative translation and splicing. Of these, 28 proteins were not documented even in the latest TAIR11 proteome database (**Table 2**). The sequences of novel identifications and their matched peptides as well as the matched spectra if the spectral count is smaller than 4 are presented in Supplementary File 1. Interestingly, half of these novel proteins (n=14) were annotated as transposable element genes in TAIR webpage. There were 2 novel proteins annotated as novel transcribed region, 2 as long noncoding RNA, and 1 antisense long RNA. Four proteins (Table 2, No. 20-24) were probably alternatively transcribed and translated proteins since there were other proteins from the same genomic locus. There were also 5 proteins (Table 2, No. 24-28) with slightly different amino acids from the proteins in the predicted TAIR proteome.

**Table 2.**
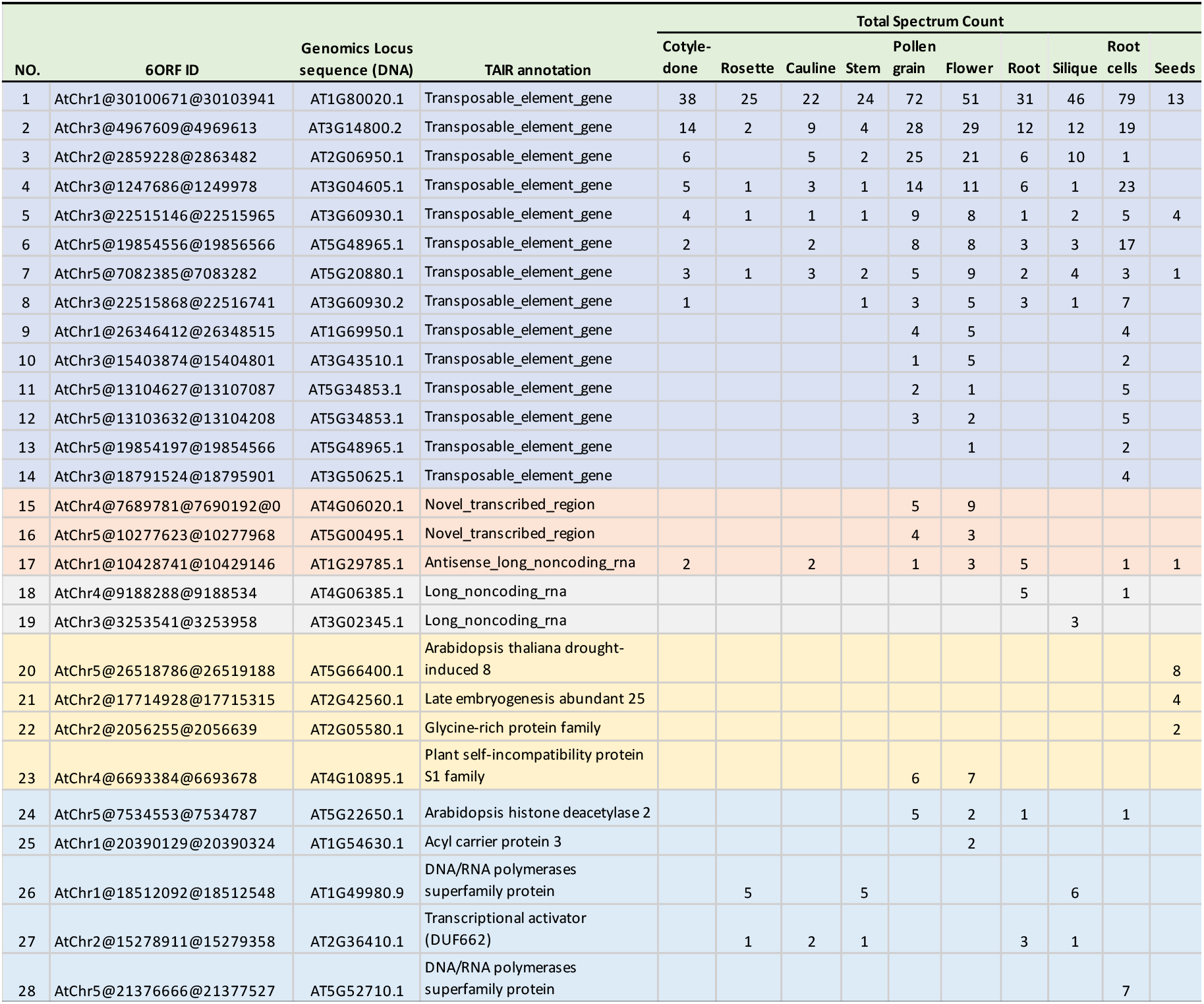
The list of novel proteins identified using proteogenomics approach. These proteins were not present in TAIR proteome database.

| NO. | 6ORF ID | Genomics Locus sequence (DNA) | TAIR annotation | Total Spectrum Count |  |  |  |  |  |  |  |  |  |
| --- | --- | --- | --- | --- | --- | --- | --- | --- | --- | --- | --- | --- | --- |
|  |  |  |  | Cotyle-<br>done | Rosette | Cauline | Stem | Pollen<br>grain | Flower | Root | Silique | Root<br>cells | Seeds |
| 1 | AtChr1@30100671@30103941 | AT1G80020.1 | Transposable_element_gene | 38 | 25 | 22 | 24 | 72 | 51 | 31 | 46 | 79 | 13 |
| 2 | AtChr3@4967609@4969613 | AT3G14800.2 | Transposable_element_gene | 14 | 2 | 9 | 4 | 28 | 29 | 12 | 12 | 19 |  |
| 3 | AtChr2@2859228@2863482 | AT2G06950.1 | Transposable_element_gene | 6 |  | 5 | 2 | 25 | 21 | 6 | 10 | 1 |  |
| 4 | AtChr3@1247686@1249978 | AT3G04605.1 | Transposable_element_gene | 5 | 1 | 3 | 1 | 14 | 11 | 6 | 1 | 23 |  |
| 5 | AtChr3@22515146@22515965 | AT3G60930.1 | Transposable_element_gene | 4 | 1 | 1 | 1 | 9 | 8 | 1 | 2 | 5 | 4 |
| 6 | AtChr5@19854556@19856566 | AT5G48965.1 | Transposable_element_gene | 2 |  | 2 |  | 8 | 8 | 3 | 3 | 17 |  |
| 7 | AtChr5@7082385@7083282 | AT5G20880.1 | Transposable_element_gene | 3 | 1 | 3 | 2 | 5 | 9 | 2 | 4 | 3 | 1 |
| 8 | AtChr3@22515868@22516741 | AT3G60930.2 | Transposable_element_gene | 1 |  |  | 1 | 3 | 5 | 3 | 1 | 7 |  |
| 9 | AtChr1@26346412@26348515 | AT1G69950.1 | Transposable_element_gene |  |  |  |  | 4 | 5 |  |  | 4 |  |
| 10 | AtChr3@15403874@15404801 | AT3G43510.1 | Transposable_element_gene |  |  |  |  | 1 | 5 |  |  | 2 |  |
| 11 | AtChr5@13104627@13107087 | AT5G34853.1 | Transposable_element_gene |  |  |  |  | 2 | 1 |  |  | 5 |  |
| 12 | AtChr5@13103632@13104208 | AT5G34853.1 | Transposable_element_gene |  |  |  |  | 3 | 2 |  |  | 5 |  |
| 13 | AtChr5@19854197@19854566 | AT5G48965.1 | Transposable_element_gene |  |  |  |  |  | 1 |  |  | 2 |  |
| 14 | AtChr3@18791524@18795901 | AT3G50625.1 | Transposable_element_gene |  |  |  |  |  |  |  |  | 4 |  |
| 15 | AtChr4@7689781@7690192@0 | AT4G06020.1 | Novel_transcribed_region |  |  |  |  | 5 | 9 |  |  |  |  |
| 16 | AtChr5@10277623@10277968 | AT5G00495.1 | Novel_transcribed_region |  |  |  |  | 4 | 3 |  |  |  |  |
| 17 | AtChr1@10428741@10429146 | AT1G29785.1 | Antisense_long_noncoding_rna | 2 |  | 2 |  | 1 | 3 | 5 |  | 1 | 1 |
| 18 | AtChr4@9188288@9188534 | AT4G06385.1 | Long_noncoding_rna |  |  |  |  |  |  | 5 |  | 1 |  |
| 19 | AtChr3@3253541@3253958 | AT3G02345.1 | Long_noncoding_rna |  |  |  |  |  |  |  | 3 |  |  |
| 20 | AtChr5@26518786@26519188 | AT5G66400.1 | Arabidopsis thaliana drought-induced 8 |  |  |  |  |  |  |  |  |  | 8 |
| 21 | AtChr2@17714928@17715315 | AT2G42560.1 | Late embryogenesis abundant 25 |  |  |  |  |  |  |  |  |  | 4 |
| 22 | AtChr2@2056255@2056639 | AT2G05580.1 | Glycine-rich protein family |  |  |  |  |  |  |  |  |  | 2 |
| 23 | AtChr4@6693384@6693678 | AT4G10895.1 | Plant self-incompatibility protein S1 family |  |  |  |  | 6 | 7 |  |  |  |  |
| 24 | AtChr5@7534553@7534787 | AT5G22650.1 | Arabidopsis histone deacetylase 2 |  |  |  |  | 5 | 2 | 1 |  | 1 |  |
| 25 | AtChr1@20390129@20390324 | AT1G54630.1 | Acyl carrier protein 3 |  |  |  |  |  | 2 |  |  |  |  |
| 26 | AtChr1@18512092@18512548 | AT1G49980.9 | DNA/RNA polymerases superfamily protein |  | 5 |  | 5 |  |  |  | 6 |  |  |
| 27 | AtChr2@15278911@15279358 | AT2G36410.1 | Transcriptional activator (DUF662) |  | 1 | 2 | 1 |  |  | 3 | 1 |  |  |
| 28 | AtChr5@21376666@21377527 | AT5G52710.1 | DNA/RNA polymerases superfamily protein |  |  |  |  |  |  |  |  | 7 |  |

Fourteen 14 out of the 42 novel identifications were now found in the latest TAIR11 proteome database, confirming the validity of our proteogenomics approach (Table 3). The sequences of these novel identifications and their matched peptides as well as the matched spectra if the spectral count is smaller than 4 are presented in Supplementary File 2. These include 2 new isoforms/entries (Table 3, No.1-2) that were added in TAIR11 in that they share most part of sequences with other isoforms/entries in TAIR10 but at least 1 unique peptide sequence were identified in this study. Six proteins (Table 3, No. 3-8) were only identified in this study and in TAIR11 but not in TAIR10 proteome database; and that included two earlier annotated as “transposable element gene”. One protein (AT5G13590.1) was identified with difference in two amino acids (in TAIR10: T*R*GAFLNSNR, and *D*EEPTELNLSLSK; this study, they were: T*S*GAFLNSNR and *N*EEPTELNLSLSK). There were also 5 proteins (Table 3, No.10-14) containing additional sequences that are not present in TAIR10 protein database. Taken it together, these data provide direct experimental evidence confirming the new revision of protein sequences.

**Table 3.**
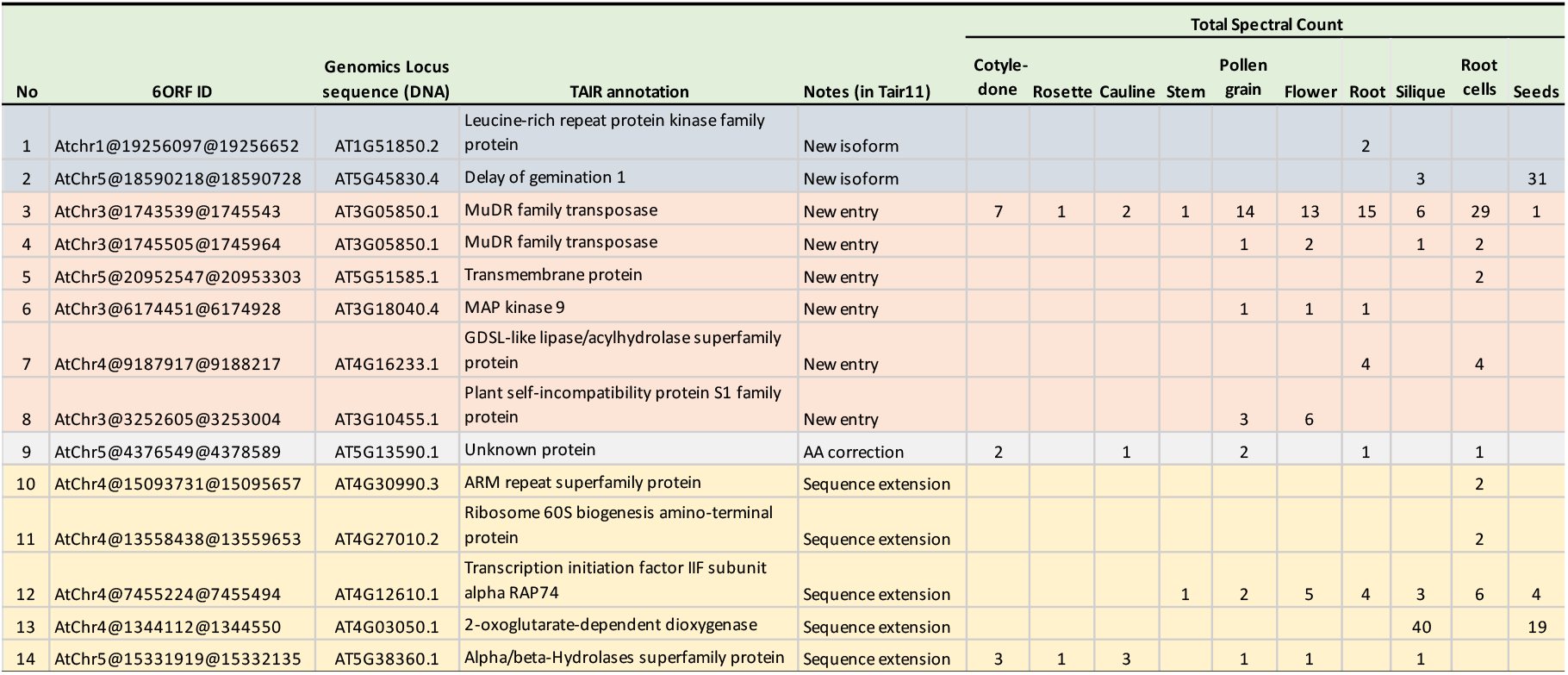
The list of novel proteins identified using proteogenomics approach. These proteins/sequences were either not present in TAIR10 or incomplete, but present in the latest TAIR11 proteome database.

In the subsequent DIA-MS analysis of ABA-treated leave sample, seven novel proteins (Table 2: No.1-5, No.7 and Table 3: No.3) with their relatively high spectral counts were also identified and quantified. Of these, the protein AtChr1@30100671@30103941 was observed to be down-regulated (fold change, 0.57, q<0.0027) at 2 h post ABA-treatment, suggesting that this “transposable element gene” protein is not only expressed at relatively high abundance but may be also functionally active.

### *Arabidopsis* spectral libraries

In order to quantify proteome dynamics in Arabidopsis by DIA-MS, we constructed a combined spectral library as well as platform-specific libraries from individual LC-MS platform (Table 4) using Spectronaut Pulsar. The library from Orbitrap Fusion analysis was comprised of 15,514 protein groups, 187,265 unique peptide sequences, and 278,278 precursors, while the library from the TripleTOF analysis contains 10,915 protein groups, 80,492 peptides, and 118,475 precursors. The combined library was comprised of a similar number of proteins (15,485) and slightly higher number of 284,418 precursors, suggesting that Orbitrap Fusion apparently recovered nearly all proteins and peptides that from TripleTOF 5600 mass spectrometry in this study.

**Table 4.**
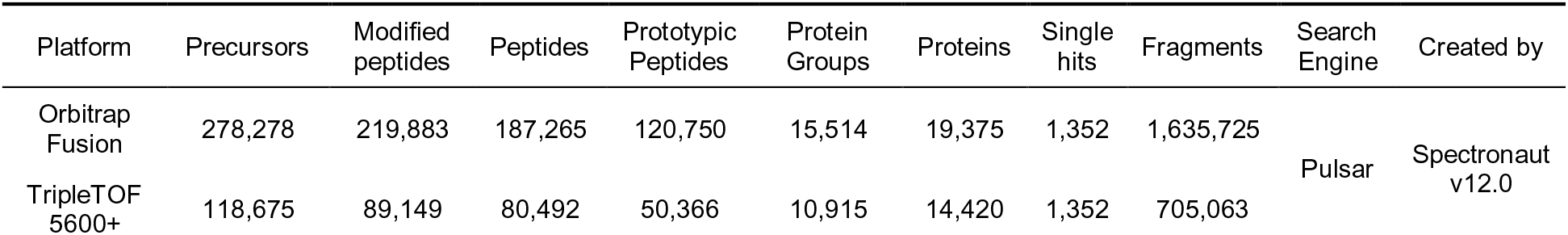
Arabidopsis spectral libraries constructed from Orbitrap Fusion and TripleTOF 5600.

| Platform | Precursors | Modified peptides | Peptides | Prototypic Peptides | Protein Groups | Proteins | Single hits | Fragments | Search Engine | Created by |
| --- | --- | --- | --- | --- | --- | --- | --- | --- | --- | --- |
| Orbitrap Fusion | 278,278 | 219,883 | 187,265 | 120,750 | 15,514 | 19,375 | 1,352 | 1,635,725 | Pulsar | Spectronaut v12.0 |
| TripleTOF 5600+ | 118,675 | 89,149 | 80,492 | 50,366 | 10,915 | 14,420 | 1,352 | 705,063 |  |  |

The peptide distribution of the combined library was shown in Figure 2a. Most peptides ranged from 8 to 15 amino acids in length, consistent with the properties of full tryptic peptides. Approximately 84 % of precursors falls in a mass range between 400-900 m/z (Figure 2b). In SWATH/DIA-MS analysis, most studies acquired over a mass window from 400 to 1200 Da (m/z) to obtain full coverage of peptides. The large scan window compromises the resolution and increases the complexity of mass spectra, leading to fewer peptide identifications. Our data could provide a valuable reference for designing DIA-MS methods with optimal mass windows. Approximately 50% of precursors have a charge of 2 (Figure 2c), and 15% are cysteine modified peptides (Figure 2d). The large number of cysteine-containing peptides may serve as a useful resource for redox proteomics quantitation.

**Figure 2.**
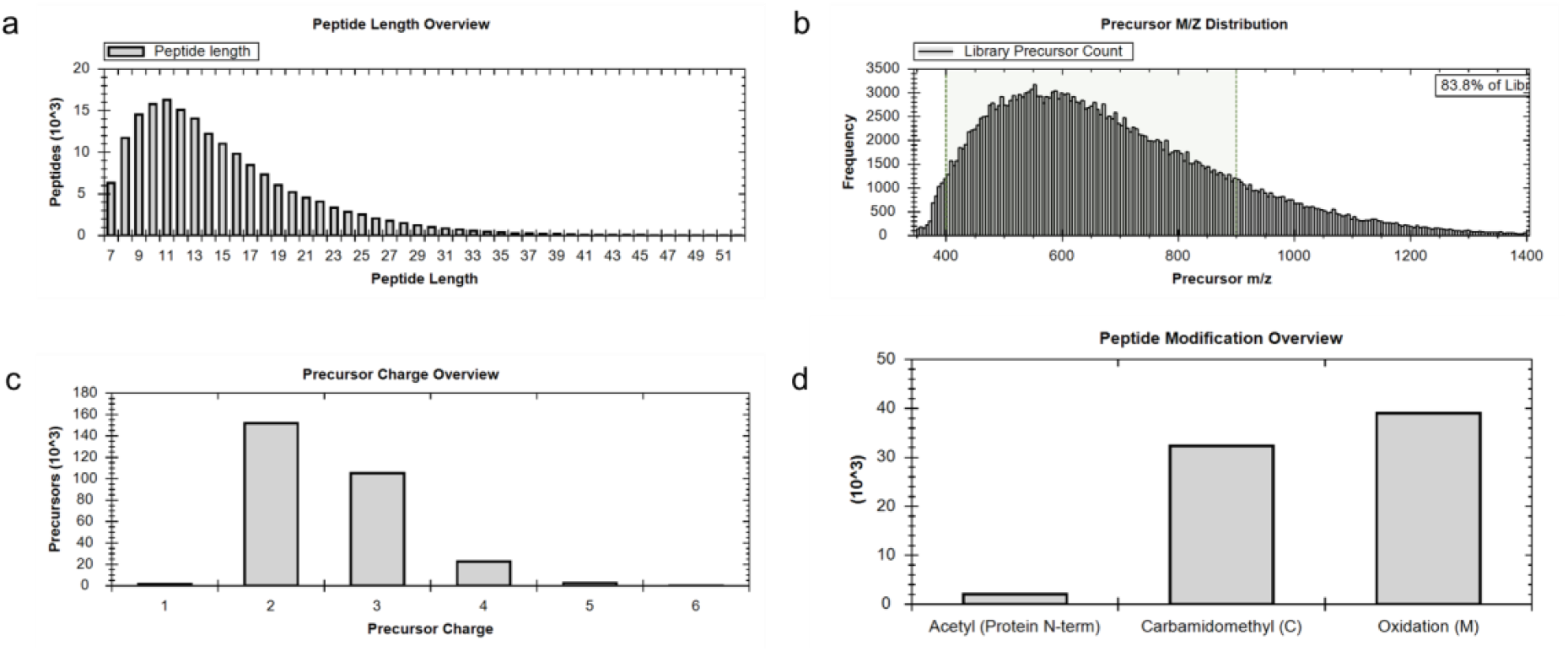
Peptide properties in the *Arabidopsis* spectral library.

### DIA-MS analysis

To demonstrate the usefulness of our assay library, we performed DIA analysis of ABA-treated *Arabidopsis* leaf sample. A total of 8,793 protein groups were quantified (PXD014032) with low number of missing values from replicates (Figure 3). The number of the identification represents 56.7% recovery of the *Arabidopsis* library. The median coefficient of variation (CV) for the experiment was below 10%, indicating high reproducibility and high quantitation accuracy (Figure S1). Together, these clearly showed the advantages of DIA-analysis for the high-throughput quantitation analysis for the *Arabidopsis.*

**Figure 3.**
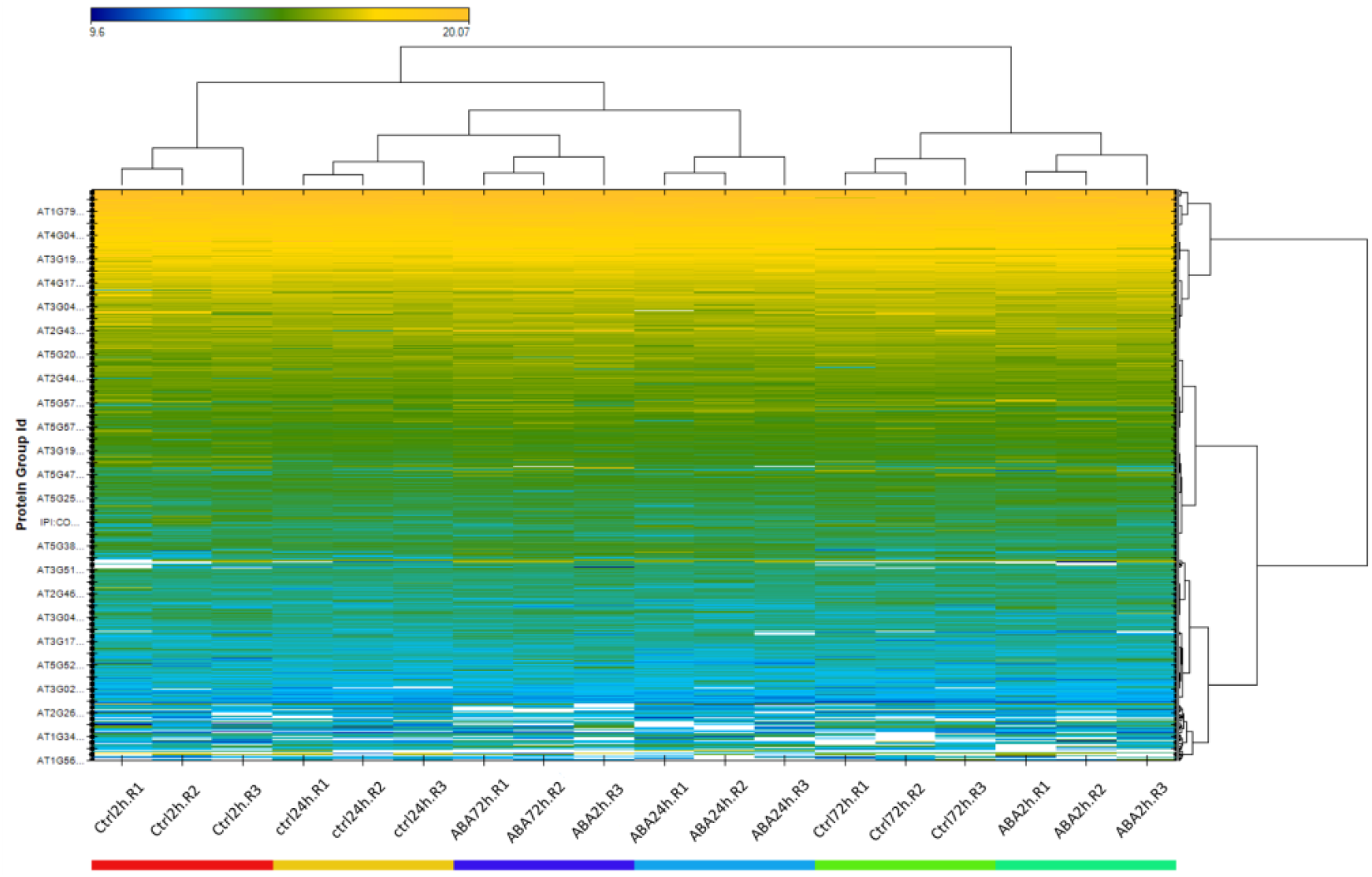
Heatmap shows the clustering of 3 technical replicates under 6 experimental conditions. Runs within the same condition cluster nicely as illustrated by the condition-based color code in the bottom of the heatmap and the x-axis dendrogram.

**Figure S1.**
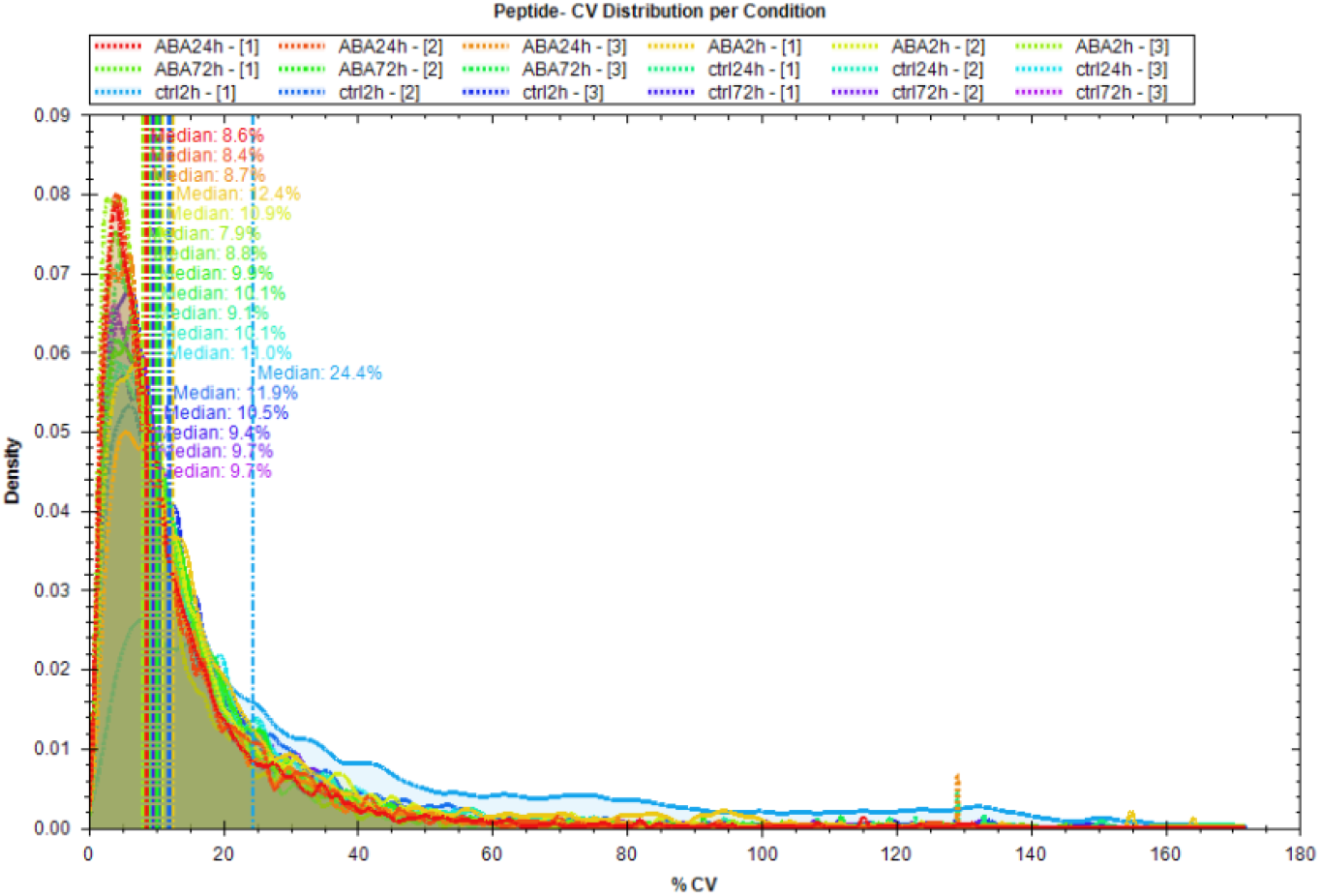
The Coefficients of Variation (%CV) distribution of all 18 sample. The low CV indicates high quantitation accuracy.

### Functional analysis of the abscisic acid (ABA)-regulated proteins

The abundance of proteins in ABA-treated sample was directly compared to their controls at the same developmental stage and with the same treatment time (e.g. ABA 2h versus control 2h; ABA 24h versus control 24h) to eliminate growth effect. Of the 8,793 protein groups, 1,787 were found to be regulated by ABA treatment at least at one of the three measured time points. To gain insights into the ABA-regulated proteome, we used the DAVID functional annotation tools to perform GO enrichment analysis (biological processing) of all ABA-responding proteins. The enriched GO terms were plotted versus their enrichment p-values (logarithm transformation of Benjamin corrected p-value) of the GO terms biological process (Figure 4). The enriched biological process (BP) term of down-regulated protein groups were shown in Figure 4a,b,c, whereas those upregulated were showed in Figure 4d,e,f. There were slightly fewer GO terms enriched at 24 h treatment compared with 2 h and 72 h treatment.

Not surprisingly, the “response to abscisic acid” was enriched in all conditions. The other highly enriched BP terms were “oxidative-reduction process”, “hydrogen peroxide catabolic process” and “response to oxidative stress”. These oxidative-stress related processes were enriched among the up- and down-regulated proteins at all measured time points, indicating ABA treatment induced active reactive oxygen species (ROS) production. Indeed, it has been well documented that ABA can cause oxidative stress in *Arabidopsis*^27–29^. Several metabolic processes including carbohydrate metabolic process, sucrose biosynthetic/metabolic process, chitin catabolic process and macromolecule catabolic process were downregulated at the 2 h post-treatment (Figure 4a), whereas they were enriched from the upregulation group of proteins (Figure 4f) at 72 h post-treatment, indicating the ABA treatment initially reduces metabolism followed by gradually increasing the metabolism to the highest level at 72 h post treatment. The other highly significant enrichments at 2 h included RNA secondary unwinding, translation, rRNA processing and ribosome biogenesis (Figure 4d), all of which are related to gene transcription and translation, suggesting that protein synthesis was immediately activated in response to ABA stimuli.

**Figure 4.**
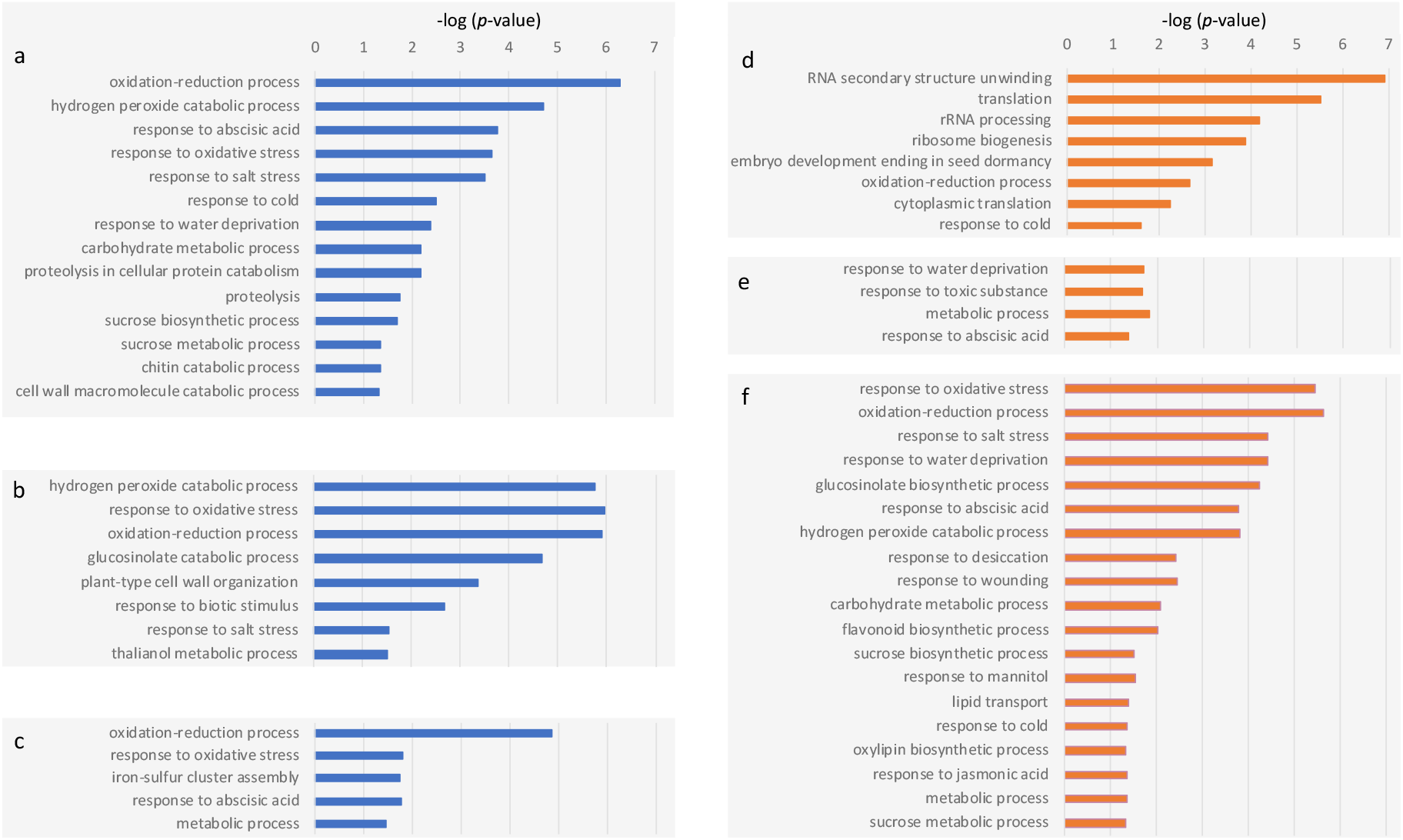
Gene ontology enrichment analysis (biological process: BP) of differential expressed proteins in response to ABA treatment. Blue bar indicates down-regulation of biological processes at 2 h (a), 24 h (b), and 72 h post-treatment (d) respectively, whereas the orange bar indicates upregulation of biological process at 2 h (d), 24 h (e), and 72 h post-treatment (f) respectively. Only BP terms with a Benjamini corrected p-value of less than 0.05 were included.

### Pathway alteration in response to ABA treatment

Based on KEGG database, pathway ath03010: Ribosome was the most significant enrichment (with Benjamini adjusted p-value of 8.41E-14) from the upregulated proteins at 2 h post treatment. This significance attenuates at both 24 h (a p-value of 0.022) and at 72 h post-treatment (p-value >0.05). Other significant enrichments were related to biosynthesis and metabolic pathways. Most of these pathways were down-regulated at 2 h (ath00940: Phenylpropanoid biosynthesis; ath01100: Metabolic pathways; ath01110: Biosynthesis of secondary metabolites) and 24 h (ath00940: Phenylpropanoid biosynthesis; Biosynthesis of secondary metabolites). In contrast, they were enriched from the upregulated proteins at 72 h post-treatment (ath00940:Phenylpropanoid biosynthesis; ath01110: Biosynthesis of secondary metabolites; ath01100: Metabolic pathways). These results, consistent with the GO BP enrichment analysis, suggesting that both transcription and translation were more active upon ABA treatment and they gradually returned to normal after days, whereas the metabolic pathways are suppressed at earlier stages and subsequently became more active.

### Temporal profiling of known abscisic acid responsive targets

Among the differentially expressed proteins included 64 previously known as “response to abscisic acid”. In this study, we further revealed their temproal profiling after treatment. Of these, 37, 18 and 42 proteins were differentially expressed at 2 h, 24 h, and 72 h post treatment respectively. Based on Figure of Merit analysis, these proteins were clustered into 4 groups using k-means clustering (Figure S2). Cluster 1 contains 25 proteins that were down-regulated at 2 h but their expression level increased thereafter (Figure 5a). Cluster 2 has 5 proteins with their lower expression at 24 h post treatment (Figure 5b). Cluster 3 contains 11 proteins with their expression of down regulated at 2 h but upregulated at 24 h and 72 h (Figure 5b). Cluster 4 has 22 proteins with the highest expression level at 2 h, and gradually dropped to lower level at 72 h post treamtent (Figure 5d). Proteins from the same cluster likely share similar functions. Indeed, based on String network analysis, proteins interacted with each other and shown in blue (Figure S3) were mostly from cluster 4, whereas proteins shown in light-green were mostly from cluster 1.

**Figure S2.**
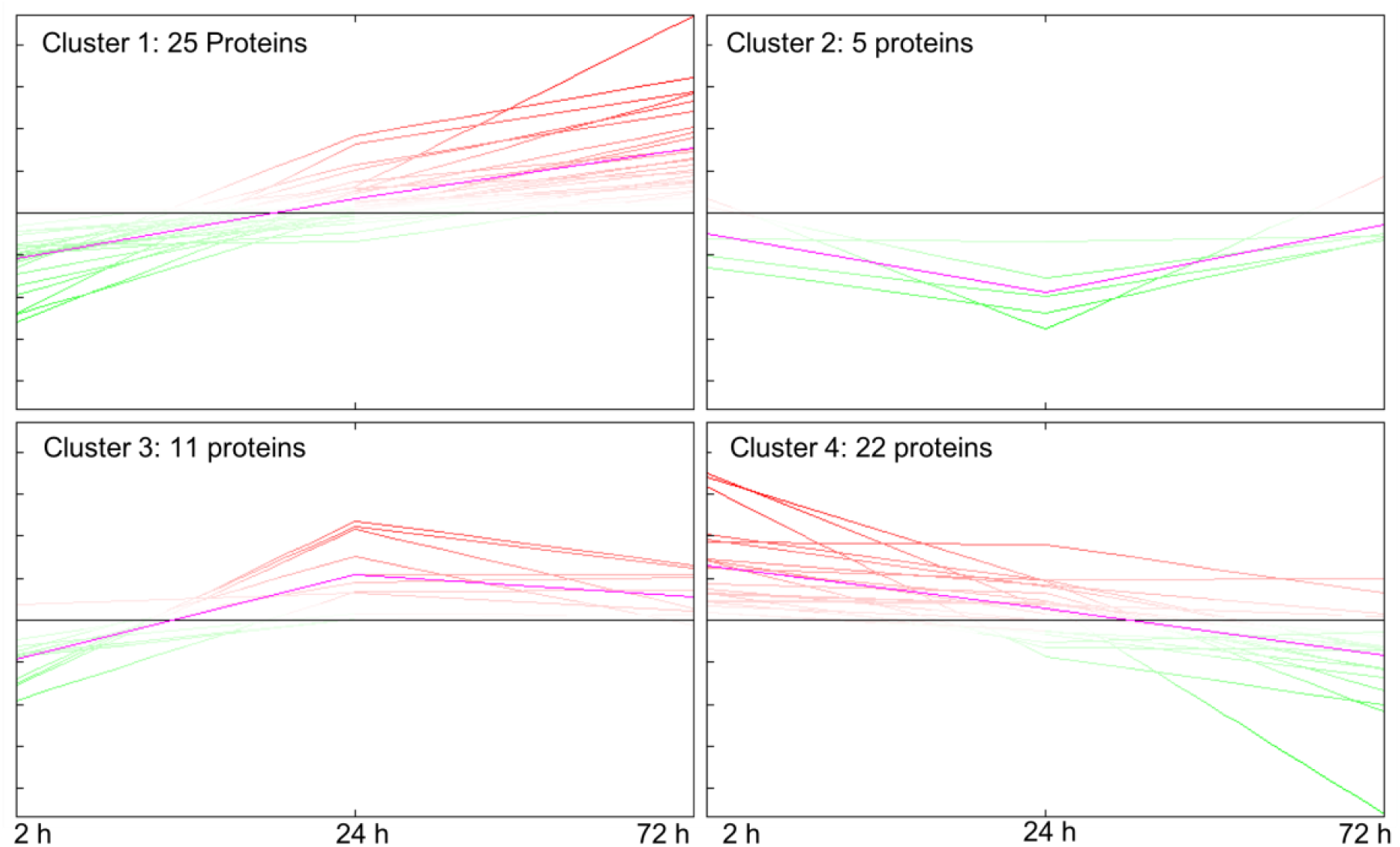
K-mean analysis of protein expression clustered into 4 clusters.

**Figure 5.**
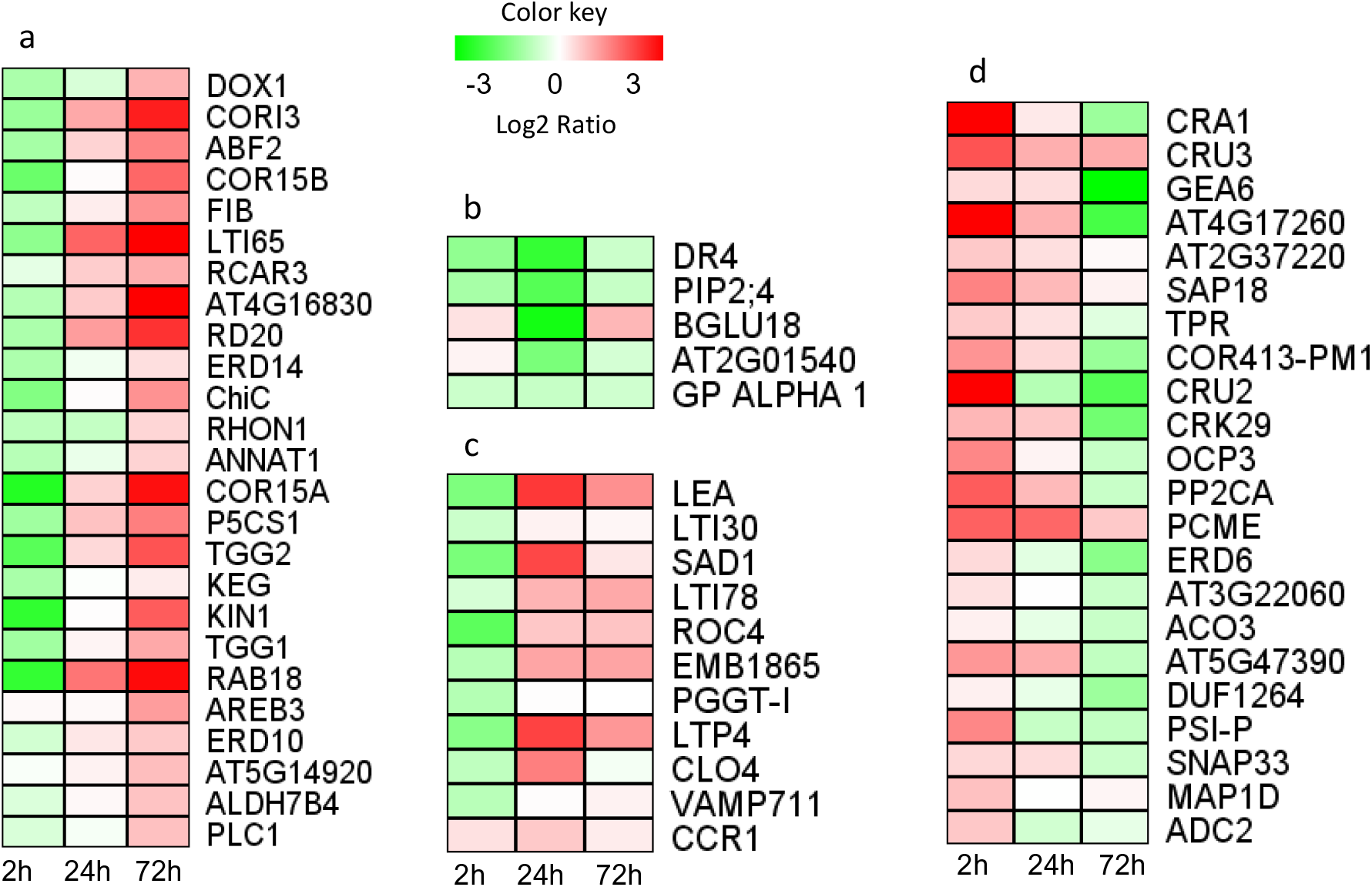
Heatmap shows the temporal profiling of previously known ABA-responding proteins. The proteins were clustered into 4 groups based on k-mean analysis, with most of proteins tending to either constantly increase (a) or decrease (d) their expression within the measured time frames. Red indicates upregulation and green as downregulation.

**Figure S3.**
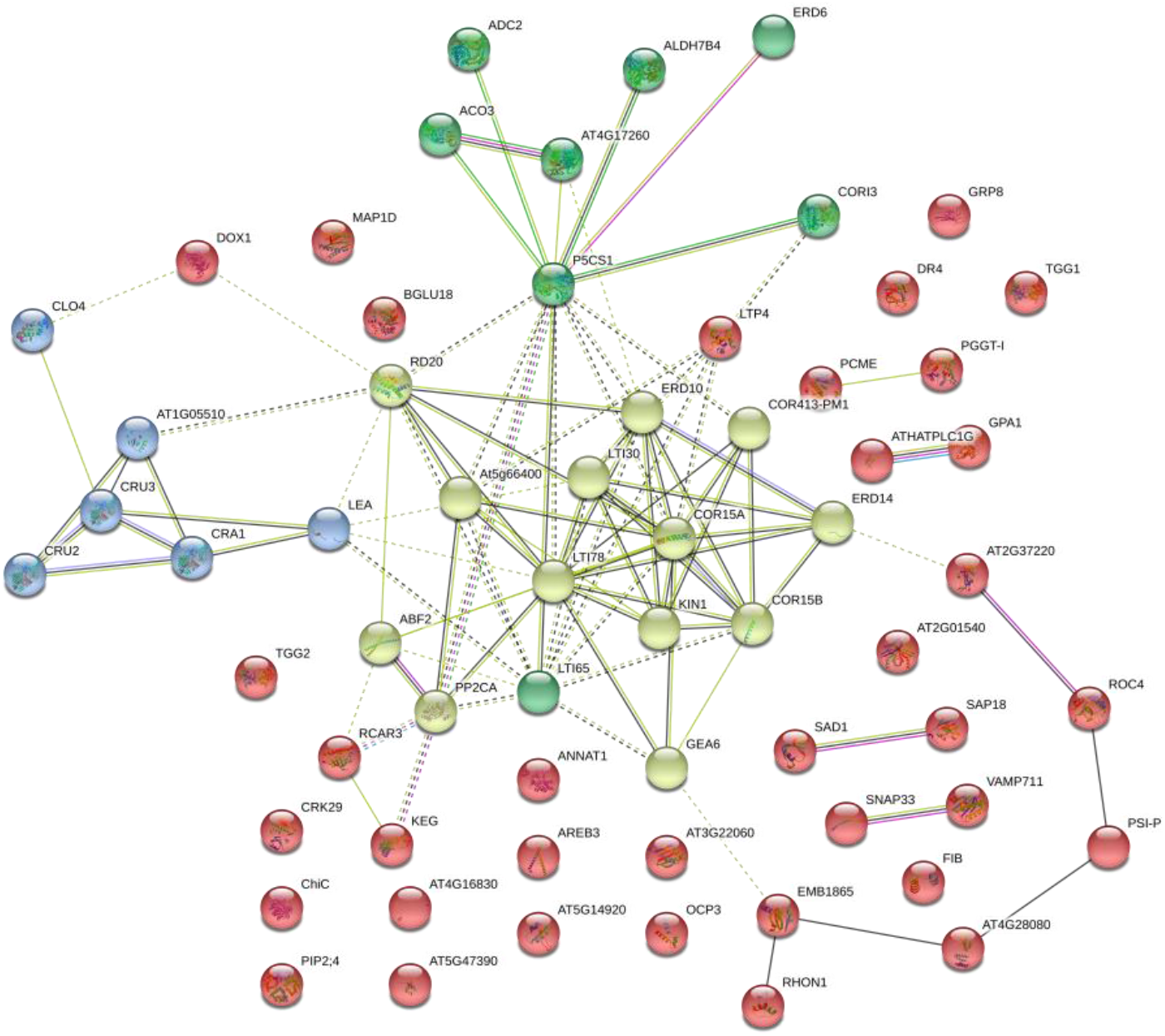
String analysis revealed protein-protein interaction network complex. The PPI enrichment have a *p*-value less than 1.0e-16, a total of 63 nodes and 101 edges, and average node degree of 3.21. The network partitioned into 4 clusters based on k-mean clustering algorithm.

## Usage Notes

In this study, we generated the largest set of tissue-specific DDA data using two high resolution of mass spectrometry platforms. This in-depth proteomics spectral information enabled us to identify and validate novel proteins using proteogenomics analysis by a 6-frame translation approach. These data are made available as a resource to the research community. Researchers can use these data to further annotate *Arabidopsis* genome data and correlate with RNAseq data.

This study provided the first comprehensive *Arabidopsis* spectral libraries, either instrument-specific or a combined library. We used this library in DIA-MS and quantified a higher number of proteins (6,000-9,000) in study of Arabidopsis proteome dynamics upon abscisic acid treatment, suggesting that rich information can be obtained in a highthroughput approach. Since this library contained approximately 30,000 cysteine-modified peptides, it can be also used for redox study of reversible cysteine modification. In addition, targeted proteomics using SRM and PRM can be designed based on this reference spectral library.

## Acknowledgements

We thank the facilities director of bioscience and analytical core labs, Stine Buechmann-Moeller, for her endorsement and support in this project.

## Author contributions

Study design: Z.H., G.T., R.A. and X.L. Experiment: Z.H., L.P. and Z.HY. Data analysis: Z.H., L.P. and G.T. Manuscript preparation: Z.H., L.P., D.B. and X.L.

## Additional Information

Competing interests: The authors declare no competing interests.

