## Supplementary File 1 for "Arabidopsis Proteome and the Mass Spectral Assay Library"

**Table 2. The list of novel proteins identified using proteogenomics approach. These proteins were not present in TAIR proteome database**. The detailed information about the novel identification, its annotation from TAIR webpage and their matched peptides (highlighted in yellow) were presented following with this table.

Furthermore:

- If the spectral counts are smaller than 4, the matched spectra were also presented.
- If the novel identification shares with TAIR10 protein entry a group of peptides, then the matched TAIR10 protein entry were included, and the unique peptide(s) associated with the novel identification were specified.
- If different set of peptides (from different tissues) were associated with the novel identification, all sequence matches were listed.

| NO. | 6ORF ID | Genomics Locus sequence (DNA) | TAIR annotation | Total Spectrum Count | | | | | | | | | |
|---|---|---|---|---|---|---|---|---|---|---|---|---|---|
| | | | | Cotyle-done | Rosette | Cauline | Stem | Pollen grain | Flower | Root | Silique | Root cells | Seeds |
| 1 | AtChr1@30100671@30103941 | AT1G80020.1 | Transposable_element_gene | 38 | 25 | 22 | 24 | 72 | 51 | 31 | 46 | 79 | 13 |
| 2 | AtChr3@4967609@4969613 | AT3G14800.2 | Transposable_element_gene | 14 | 2 | 9 | 4 | 28 | 29 | 12 | 12 | 19 | |
| 3 | AtChr2@2859228@2863482 | AT2G06950.1 | Transposable_element_gene | 6 | | 5 | 2 | 25 | 21 | 6 | 10 | 1 | |
| 4 | AtChr3@1247686@1249978 | AT3G04605.1 | Transposable_element_gene | 5 | 1 | 3 | 1 | 14 | 11 | 6 | 1 | 23 | |
| 5 | AtChr3@22515146@22515965 | AT3G60930.1 | Transposable_element_gene | 4 | 1 | 1 | 1 | 9 | 8 | 1 | 2 | 5 | 4 |
| 6 | AtChr5@19854556@19856566 | AT5G48965.1 | Transposable_element_gene | 2 | | 2 | | 8 | 8 | 3 | 3 | 17 | |
| 7 | AtChr5@7082385@7083282 | AT5G20880.1 | Transposable_element_gene | 3 | 1 | 3 | 2 | 5 | 9 | 2 | 4 | 3 | 1 |
| 8 | AtChr3@22515868@22516741 | AT3G60930.2 | Transposable_element_gene | 1 | | | 1 | 3 | 5 | 3 | 1 | 7 | |
| 9 | AtChr1@26346412@26348515 | AT1G69950.1 | Transposable_element_gene | | | | | 4 | 5 | | | 4 | |
| 10 | AtChr3@15403874@15404801 | AT3G43510.1 | Transposable_element_gene | | | | | 1 | 5 | | | 2 | |
| 11 | AtChr5@13104627@13107087 | AT5G34853.1 | Transposable_element_gene | | | | | 2 | 1 | | | 5 | |
| 12 | AtChr5@13103632@13104208 | AT5G34853.1 | Transposable_element_gene | | | | | 3 | 2 | | | 5 | |
| 13 | AtChr5@19854197@19854566 | AT5G48965.1 | Transposable_element_gene | | | | | | 1 | | | 2 | |
| 14 | AtChr3@18791524@18795901 | AT3G50625.1 | Transposable_element_gene | | | | | | | | | 4 | |
| 15 | AtChr4@7689781@7690192@0 | AT4G06020.1 | Novel_transcribed_region | | | | | 5 | 9 | | | | |
| 16 | AtChr5@10277623@10277968 | AT5G00495.1 | Novel_transcribed_region | | | | | 4 | 3 | | | | |
| 17 | AtChr1@10428741@10429146 | AT1G29785.1 | Antisense_long_noncoding_rna | 2 | | 2 | | 1 | 3 | 5 | | 1 | 1 |
| 18 | AtChr4@9188288@9188534 | AT4G06385.1 | Long_noncoding_rna | | | | | | | 5 | | 1 | |
| 19 | AtChr3@3253541@3253958 | AT3G02345.1 | Long_noncoding_rna | | | | | | | | 3 | | |
| 20 | AtChr5@26518786@26519188 | AT5G66400.1 | Arabidopsis thaliana drought-induced 8 | | | | | | | | | | 8 |
| 21 | AtChr2@17714928@17715315 | AT2G42560.1 | Late embryogenesis abundant 25 | | | | | | | | | | 4 |
| 22 | AtChr2@2056255@2056639 | AT2G05580.1 | Glycine-rich protein family | | | | | | | | | | 2 |
| 23 | AtChr4@6693384@6693678 | AT4G10895.1 | Plant self-incompatibility protein S1 family | | | | | 6 | 7 | | | | |
| 24 | AtChr5@7534553@7534787 | AT5G22650.1 | Arabidopsis histone deacetylase 2 | | | | | 5 | 2 | 1 | | 1 | |
| 25 | AtChr1@20390129@20390324 | AT1G54630.1 | Acyl carrier protein 3 | | | | | | 2 | | | | |
| 26 | AtChr1@18512092@18512548 | AT1G49980.9 | DNA/RNA polymerases superfamily protein | | 5 | | 5 | | | | 6 | | |
| 27 | AtChr2@15278911@15279358 | AT2G36410.1 | Transcriptional activator (DUF662) | | 1 | 2 | 1 | | | 3 | 1 | | |
| 28 | AtChr5@21376666@21377527 | AT5G52710.1 | DNA/RNA polymerases superfamily protein | | | | | | | | | 7 | |

#### No.1. AtChr1@30100671@30103941: Transposable_element_gene

**AtChr1@30100671@30103941 (100%), 122,861.1 Da**
**| potential novel**
**13 exclusive unique peptides, 13 exclusive unique spectra, 38 total spectra, 165/1090 amino acids (15% coverage)**

| | | | | |
|---|---|---|---|---|
| FYFASPGLFF | PAKDMDTNSL | MLIDNNGSFE | IDDQSHVHVD | ASHTLEPVTL |
| PTKRRRKKSD | VWNHFTVETV | SPGSAKACCN | HCRKSFAYIN | GQKLAGTSHL |
| KRHIQLGICP | MNR**DDSSTLA** | **QIVTPTTTTD** | **PPK**KRHR**SSA** | **SYTPLDQDR**C |
| YNQMAKMIIM | HDYPLHMVEH | SGFTGFVQAL | CPQFTMASFN | TIHADCVNMY |
| LSEKHK**LSNF** | **ITDIPGR**VNL | TVDLWTSNQS | LGYAFVTGHF | IDRDWKLTHR |
| LLNVAVVPSP | DSDFALNQPI | AACLSDWNLE | RR**LSSITVGQ** | **SVVNK**TSIEN |
| LRCCLSAR**NQ** | **NVLNGQLLLG** | **NCYARLLSSM** | **AQDLLGAEDF** | **KTPIKK**VRDS |
| VKYVKTKDSC | QERFDELKKQ | LQTPSTK**DLL** | **IDNQTK**WDTS | YDMLLAACEH |
| KEVFSCLGNC | DLDYKILTSP | EEWRKIEILC | SSLK**ILFDAA** | **NVLTGSTRLT** |
| **ANDLFHEMTK** | LQLDLGNSAT | SEDQDVSNLA | NTLKDKFDEY | WRECFLLMAV |
| AVVMDPRFKM | K**LIEFSFSK**A | YGEDADKWIR | SVDDAVHELY | HDYAEQSHSL |
| LDAYVVHGND | GFSETDMSQV | HFHHEYHNSN | GLSHDQIFEQ | PEDGNPLNEK |
| PLEGLSGEGQ | PTGTESVGGN | TTQGVEQEGH | LGDAGTQENN | IVEGQLEESK |
| PMDGLTQKPL | LTEENLQSNG | EPTTAQESRL | TDETMEETHA | NQSAEEMAHE |
| TQLVEEFPCE | SQPSENVPLQ | SQSGEGNVPE | TLPMEEIVED | TQPVEEVMQE |
| ETSQVSRPVE | EIPLENQQVD | DKDITHDVQP | VEEMLEDTQP | VEGVDHEAQE |
| LEVHDDIQSV | EEVGHEIQPE | GELLEDTQPV | EEGAQEAQLV | EENHDDVQLV |
| EEAGHETHAV | EGILEDTQPV | EEVAQEARPV | EQIPENSQNP | QEYSREGESV |
| QEEQQSEQHP | QSHAMPHEEE | AQHDSQSHAM | PQEEATFTIS | QEGHHVDVLL |
| QEGHHLEASS | QEFPLITIGD | GFSDFELYIS | EVGSHQQMK**S** | **ELDQYLEESL** |
| **IPRSQDFEVL** | **GWWSLNR**TKY | PTLSKMAADV | LSVPFCTVSP | DSVFDTEVKK |
| MDNYRSSLRH | VTLEALFCAK | DWFKHSSSNS | TSENNLKMES | |

#### No.2. AtChr3@4967609@4969613: Transposable_element_gene

**AtChr3@4967609@4969613 (100%), 76,551.7 Da**
**| potential novel**
**5 exclusive unique peptides, 5 exclusive unique spectra, 19 total spectra, 73/668 amino acids (11% coverage)**

| | | | | |
|---|---|---|---|---|
| QGYKFCMMDE | SNEIILQKSK | RLTSVVWNYF | ERVRKADVCY | AVCIQCNKKL |
| SGSSNSGTTH | LRNHLMRCLK | RTNHDMSQLL | TPKRRKKENP | VTVATINFDD |
| GQAKEEYLRP | KFDQDQRRDE | VVLSRGSGGR | FSQERSQVDL | AR**MIILHNYP** |
| **LAMVDHVGFK** | VFARNLQPLF | EAVPNSTIED | SCMEIYIREK | QRVQHTLNHL |
| YGKVNLSVEM | WSSRDNSNYV | CLASNYIDEE | WRLHRNVLNF | ITLDPSHTED |
| MLSEVIIRCL | IEWSLENKLF | AVTFDSVSVN | EEIVLRIKDH | MSQSSQILIN |
| GQLFELKSAA | HLLNSLVEDC | LEAMRDVIQK | IRGSVRYVKS | SQSTQVRFNE |
| IAQLAGINSQ | KILVLDSIVN | SNSTFVMLET | VLEYKGAFCH | LRDHDHSFDS |
| SLTDEEWEWT | RYVTGYLKLV | FDIASDFSAN | KCPTANVYFA | EMCDIHIQLV |
| EWCKNQDNFL | SSLAANMKAK | FDEYWNKCSL | VLAIAAILDP | RFKMKLVEYY |
| YSK**IYGSTAL** | **DR**IKEVSNGV | KELLDAYSMC | SAIVGEDSFS | GSGLGRASMD |
| TRDRLKGFDK | **FLHETSQNQN** | **TTTDLDKYLS** | **EPIFPRSGEF** | **NILNYWK**VHT |
| PR**YPILSLLA** | **R**DILGTPMSI | CAPDSTFNSG | TPVISDSQSS | LNPDIRQALF |
| CAHDWLSTET | EGTISNSI | | | |

#### No.3. AtChr2@2859228@2863482: Transposable_element_gene

**AtChr2@2859228@2863482 (100%), 158,765.0 Da**
**| potential novel**
**5 exclusive unique peptides, 5 exclusive unique spectra, 25 total spectra, 65/1418 amino acids (5% coverage)**

SIAFAFMVSE LMEHRSMELY SVPSLNISNC VTVTLTAKNY ILWKSQFESF
LDGQGLLGFV TGSIPAPSQT SVVSDIDGST SASPNPEYYT WFKTDRVVKS
WLLGSFLEDI LSVVVNCNTS HEVWISVANH FNRVSSSRLF ELQRRLQNVS
KRDKSMDEYL KDLK**TICDQL** **ASVGSPVTEK** MK**IFAALNGL** **GREYEPIKTT**
**IENSMDALPG** **PSLEDVIPK**L TGYDDRLQGY LEETAVSPHV AFNITTSDDS
NASGYFNAYN RGKGKSNRGR NSFSTRGRGF HQQISSTNSS SGSQSGGTSV
VCQICGKMGH PALKCWHR**FN** **NSYQYEELPR** ALAAMRITDI TDQHGNEWLP
DSAATAHVTN SPRSLQQSQP YHGSDAVMVA DGNFLPITHT GSTNLASSSG
NVPLTDVLVC PSITKSLLSV SKLTQDYPCT VEFDSDGVRI NDKATKKLLI
MGSTCDGLYC LKDDSQFKAF FSTRQQSASD EVWHRRLGHP HPQVLQQLVK
TNSISINKTS KSLCEACQLG KSTRLPFVSS SFTSNRPLER VHCDLWGPSP
ITSVQGFRYY AVFIDHYSRF SWIYPLKLKS DFYNIFVAFH KLVENQLNHK
ISVFQCDGGG EFVNHKFLQH LQNHGIQQHI SYPHTPQQNG LAERKHRHLV
ELGLSMLFQS KVPLKFWVEA FFTANFLINL LPTSAVEDAI SPYEKLHQTT
PDYTALRSFG CACFPTMRDY AMNKFDPRSL KCVFLGYNDK YKGYRCLYPP
TGRVYISRHV IFDETAYPFS HHYKHLHSQP TTPLLAAWFK GFESSVSQAP
PKVSPAQPPQ RKATLPTPPL FTAADFPPLP RRSPQLSQNS AAALVSQPST
TTINSTHPPA VVNESSERTI NFDSASIGDS SHSSQLLVDD TVEDLMAAPV
PTQQAPPPTN THPMITRAKV GITKPNPRYV FLSHKVTYPE PKTVTAALKH
PGWTGAMTEE MGNCSETNTW SLVPYTPNMH VLGSKWVFRT KLHADGTLNK
LKARIVAKCF LQEEGIGYLE TYSPVVRTPT VQLVLHLATA LNWELKQMDV
KNAFLHGDLN ETVYMTQPAG FVDKSKPTHV CLLHKSIYGL KQSPRAWFDK
FSTFLLEFGF FCSKSDPSLF IYAHNNNLIL LLLYVDDMVI TGNSSQTLSS
LLAALNKEFR MTDMGQLHYF LGIQVQRNQH GLFMSQQKYA EDLLVASAME
NCTPLPTPLP VQLDRVPHQE EPFTDPTYFR SIAGKLQYLT LTRPDIHFAV
NFVCQKMHQP TMSDFHLLKR ILRYIKGTIT MGISYNQNSP TLLQAYSDSD
WGNCKLTRRS VGGLCTFMAT NLVSWSSKKH PTVSRSSTEA EYRTLSDAAS
EILWLSTLLR ELGIPLPDTP ELFCDNLSAV YHTANPAFHA MIDKATKSTN
FRVLYPNGDS ILTWHENA

#### No.4. AtChr3@1247686@1249978: Transposable_element_gene

**AtChr3@1247686@1249978 (100%), 86,896.1 Da**
**| potential novel**
**5 exclusive unique peptides, 5 exclusive unique spectra, 23 total spectra, 61/764 amino acids (8% coverage)**

IKKIVSAMAN RDLMLGQNRN IGIGQGQSLV LGHSHSLGLG QNHVDVELGQ
GHNMDLGLAH HDHDHHEIDL GNPHDDHELD LGNSEDQDGE DHHHHHHHDY
MNENEISVDQ KPGHDDVDPD LVLSSQNHEF SLSDNNHQLV VGENHELDDN
LELAVDSSHE LEIDQHGEMV MVSTPVQARA LTADMTYQLT VGQEFPDVKS
CRRALRDMAI ALHFEMQTIK SDKTRFTAK**C** **SSDGCPWR**VH AAK**LPGVPTF**
**TIR**TIHESHS CGGINHLGHQ QASVQWVASS VEQRLRENPN CKPKEILEEI
HRVHGITLSY KQAWRGKERI MATMRGSFEE GYRLLPQYCE QVKRTNPGSI
ASVYGSPADN CFQRLFISFQ ASIYGFLNAC RPLLGLDRTY LKSKYLGTLL
LATGFDGDGA LFPLAFGIVD EENDENWMWF LCELHNLLET NTENMPRLTI
LSDRQK**GIVE** **GVEQNFPTAF** **HGFCMR**HLSE SFRKEFNNTL LVNYLWEAAQ
ALTVIEFEAK ILEIEEISQD AAYWIRRIPP RLWATAYFEG QRFGHLTANI
VESLNSWIAE ASGLPIIQMM ECIRRQLMTW FNERRETSMQ WTSILVPTAE
RRVAEALELA RTYQVLRANE AEFEVISHEG NNIVDIRNRC CLCRGWQLYG
LPCAHAVAAL LSCRQNVHRF TESCFTVATY RKTYSQTIHP IPDKSHWR**EL**
**SEGDPNVNKA** **ALDAIINPPK** SLRPPGRPRK RRVRAEDRGR VKRVVHCSRC
NQTGHFRTTC AAPI

**AtChr3@1247686@1249978 (100%), 86,896.1 Da**
**| potential novel**
**6 exclusive unique peptides, 6 exclusive unique spectra, 11 total spectra, 71/764 amino acids (9% coverage)**

IKKIVSAMAN RDLMLGQNRN IGIGQGQSLV LGHSHSLGLG QNHVDVELGQ
GHNMDLGLAH HDHDHHEIDL GNPHDDHELD LGNSEDQDGE DHHHHHHHDY
MNENEISVDQ KPGHDDVDPD LVLSSQNHEF SLSDNNHQLV VGENHELDDN
LELAVDSSHE LEIDQHGEMV MVSTPVQARA LTADMTYQLT VGQEFPDVKS
CRRALRDMAI ALHFEMQTIK SDKTRFTAK**C** **SSDGCPWR**VH AAKLPGVPTF
TIRTIHESHS CGGINHLGHQ QASVQWVASS VEQRLRENPN CKPKEILEEI
HRVHGITLSY KQAWRGKERI MATMRGSFEE GYRLLPQYCE QVKRTNPGSI
ASVYGSPADN CFQRLFISFQ ASIYGFLNAC RPLLGLDRTY LKSKYLGTLL
LATGFDGDGA LFPLAFGIVD EENDENWMWF LCELHNLLET NTENMPRLTI
LSDRQKGIVE GVEQNFPTAF HGFCMRHLSE SFRKEFNNTL LVNYLWEAAQ
ALTVIEFEAK ILEIEEISQD AAYWIRRIPP R**LWATAYFEG** **QR**FGHLTANI
VESLNSWIAE ASGLPIIQMM ECIRRQLMTW FNERRETSMQ WTSILVPTAE
RR**VAEALELA** **R**TYQVLR**ANE** **AEFEVISHEG** **NNIVDIR**NRC CLCRGWQLYG
LPCAHAVAAL LSCRQNVHRF TESCFTVATY RKTYSQTIHP IPDKSHWR**EL**
**SEGDPNVNKA** **ALDAIINPPK** SLRPPGRPRK RRVRAEDRGR VKRVVHCSRC
NQTGHFRTTC AAPI

#### No.5. AtChr3@22515146@22515965: Transposable_element_gene

**AtChr3@22515146@22515965 (100%), 30,005.7 Da**
**| potential novel**
**4 exclusive unique peptides, 5 exclusive unique spectra, 9 total spectra, 92/273 amino acids (34% coverage)**

RDGEFWVVVV RNLLVPEVML ISFSNCNFPF AERTLK**FLEP** **IPDDFLSAFH**
**ALSAR**KCDWL KHFSRERVEC ALRLLHGVSC PTSSESSDHR TQFFVDMQST
KLTLREVYAK KKEDKERRLA EERRLVNTGL ISPR**ADPEAT** **QDGNVIPDAT**
**APQTAPEASR** **DGVNPYTAGP** **VDAAPAEAQG** **AEPSAAAPEA** **VLALPAIDK**A
AGKRIRVDDV SSKKKKKKKK ASGSEVEK**IL** **PIFEDR**TASA NLLGGCVGPL
LPPPDTLLES RKYAETASHF LRV

#### No.6. AtChr5@19854556@19856566: Transposable_element_gene

**AtChr5@19854556@19856566 (100%), 76,200.0 Da**
**| potential novel**
**7 exclusive unique peptides, 7 exclusive unique spectra, 17 total spectra, 86/670 amino acids (13% coverage)**

VCVLVLRPNG LSTLFMIVMT DVWISVFDRS SRTTASEAVV PVVSAGDIVV
GDDMIANDFQ IEMGIMPEES SPLPCNFVLT DEKQHIK**AAQ** **QWENAITGVD**
**QRFNSFTEFR** DALHKYSIAH GFTYKYKKND SHRVSVKCKA QGCPWRITAS
R**LSTTQLICI** **K**KMNTRHTCE RAVVKAGYRA SRGWVGSIIK EKLKAFPDYK
PKDIAEDIKR EYGIQLNYSQ AWRAKEIARE QLQGSYKK**AY** **SQLPSFCK**KI
RETNPGSIAI FMTKEDSSFH RLFISFYASI SGFRQGCRPL LFLDTADLNS
KYQGVMLVAT APDAEDGIFP VAFAVVDAET EDNWVWFLEH LKLALADPRT
ITFVADFQNG LK**TALPLVFE** **K**QHHHAYCLR HLAEKLNMDL QAQFSHEARR
FILNDFYAAA YATQPDAYYR SLENIKSISP DAYTWVIESE PLHWANALFE
GERYNHMNSI FGLDFYSWVS EAHELPITQM IDELRAKLMQ SIYTHQVQSR
EWIVSTLTPT NEEKLQKEIE LARSLQVSAP HNSLFEVHGE TINLVDINQC
DCDCKVWR**LT** **GLPCSHAVAV** **VECIGKSPYE** **YCSRYFTSES** **YR**LTYAESIN
PVPNTTMTMM ILEEPPVEGV VSVTPPPTRL TPPGRPKSKQ VEPLDMFKRQ
LQCSNCKGLG HNKKTCKAVS

#### No. 7 AtChr5@7082385@7083282: Transposable_element_gene

**AtChr5@7082385@7083282 (100%), 33,959.3 Da**
**| potential novel**
**5 exclusive unique peptides, 5 exclusive unique spectra, 9 total spectra, 63/299 amino acids (21% coverage)**

RNKTCIEKEG FFVEEKFVRS LLSNSSSMAA IQILDQVDDS ADSLIESMRG
VQNTMALLHS TVGELLNSMS ETLNSMAEMK **SALLQQPTAT** **ATSESTKVTV**
**DHDLPVAPPP** **PTK**QSTAAED NSIDTHPHRM SRYIQLMKPK IEMPVFEGPN
VNSWLTRAER YFEFGSFTDA EKIQLVYMSV EGR**ALCWFNL** **ENRNPFVDWN**
**DFK**ARVLQRF GDPRSAMERL LTLKQVDSVI SYLGEFEDLS TQCLDDLDTV
LEVVFVKGLK EEIQEMLRVF QPK**GLSDIIM** **MAR**RLENSPF CRLIKNSQV

#### No. 8. AtChr3@22515868@22516741: Transposable_element_gene

**AtChr3@22515868@22516741 (100%), 32,149.4 Da**
**| potential novel**
**3 exclusive unique peptides, 3 exclusive unique spectra, 7 total spectra, 50/291 amino acids (17% coverage)**

```
SARGMCWSLA  SSSRYSSGIS  EVCRDGISLS  AGMNSVAYNL  SWCYHYKANF
SLFQAVASMN  RMVHSYDSAM  RSNMEVAGKL  AEAESRIQAI  EREKNEALSE
AAAAKLEKEE  VERTAHVNKE  NAIKMAEQNL  KANSEIVRLK  RMLSEARGLR
DSEVARAVQT  TRREVSETFI  AKMKNAEHKV  SLLDEVNDRF  MYLSQARANA
QLIEALEGGG  VLESEKEQVD  EWLKDFADAE  VNLNRFMSEL  KDDLKAPAPE
PAPLSPGGHR  SVESLADEAG  ITDQAGSLLP  AKDNRPSEDL  D
```

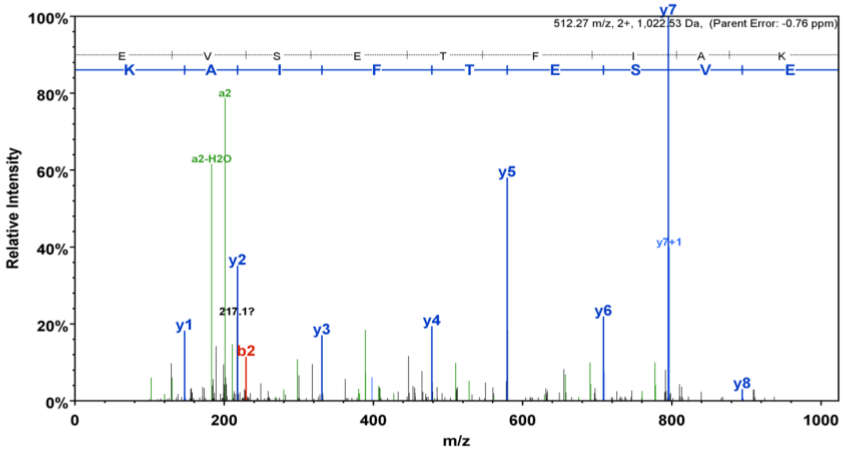


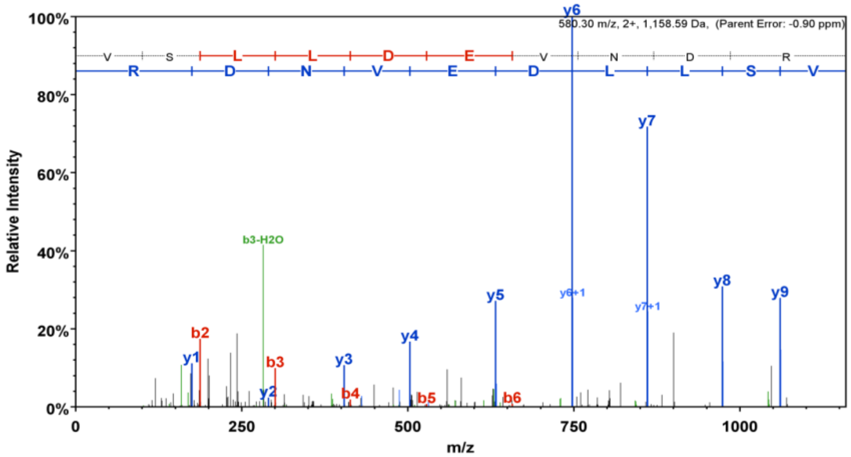


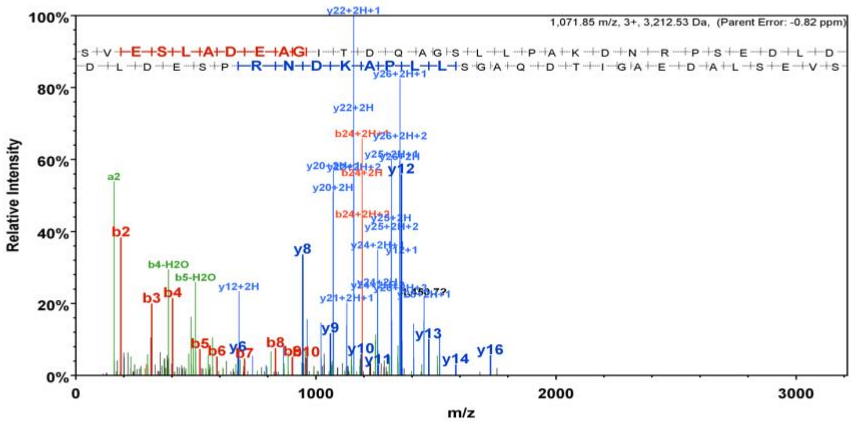

#### No. 9. AtChr1@26346412@26348515: Transposable_element_gene

**AtChr1@26346412@26348515 (100%), 81,381.4 Da**
**| potential novel**
**2 exclusive unique peptides, 2 exclusive unique spectra, 4 total spectra, 23/701 amino acids (3% coverage)**

| | | | | |
|---|---|---|---|---|
| DFRFCVTWFR | CLNSNYLVKI | LLFVYMCRRQ | KKLMINNEMD | LSDAVIVKSG |
| RLK**SVVWNDF** | **DR**VRKGETYI | AICRHCKKRL | SGSSASGTSH | LRNHLIRCRR |
| RTNGNNNGVA | QYFVKGKKKE | LANERIKDEE | VLSVVNVRYE | HEKEEHEDVN |
| VVSMGLDQRR | CRFDLARMII | LHGYPLSMVE | DVGFRMFIGN | LQPLFELVAF |
| ERVESDCMEI | YAKEKHKIFE | ALDKLPGKIS | ISVDVWSGSG | DSDEFLCLAA |
| HYIDEGWELK | KRVLNFFMVD | PSHSGEMLAE | VIMTCLMEWD | IDRKLFSMAS |
| SHAPPFSENV | ASKIRDRLSQ | NKFLYCYGQL | FDVSCGVNVI | NEMVQDSLEA |
| CCDTINIIRE | SIRYVKSSES | IQDRFNQWIV | ETGAVSERNL | CIDDPMRWDS |
| TCTMLENALE | QKSAFSLMNE | HDPDSVLCPS | DLEWERLGTI | VEFLKVFVEV |
| INAFTKSSCL | PANMYFPEVC | DIHLRLIEWS | KNPDDFISSL | VVNMRKKFDD |
| FWDKNYLVLA | IATILDPRFK | MKLVEYYYPL | FYGTSASELI | EDISECIKLL |
| YDEHSVGSLL | ASSNQALDWQ | NHHHRSNGVA | HGKEPDDRLT | EFDR**YINETT** |
| **TTPGQDSK**SD | LEKYLEEPLF | PRNSDFDILN | WWKVHTPKYP | ILSMMARNVL |
| AVPMLNVSSE | EDAFETCQRR | RVSETWRSLR | PSTVQALMCA | QDWIQSELES |
| S | | | | |

**AtChr1@26346412@26348515 (100%), 81,381.4 Da**
**| potential novel**
**2 exclusive unique peptides, 2 exclusive unique spectra, 5 total spectra, 20/701 amino acids (3% coverage)**

| | | | | |
|---|---|---|---|---|
| DFRFCVTWFR | CLNSNYLVKI | LLFVYMCRRQ | KKLMINNEMD | LSDAVIVKSG |
| RLK**SVVWNDF** | **DR**VRKGETYI | AICRHCKKRL | SGSSASGTSH | LRNHLIRCRR |
| RTNGNNNGVA | QYFVKGKKKE | LANERIKDEE | VLSVVNVRYE | HEKEEHEDVN |
| VVSMGLDQRR | CRFDLARMII | LHGYPLSMVE | DVGFRMFIGN | LQPLFELVAF |
| ERVESDCMEI | YAKEKHKIFE | ALDKLPGKIS | ISVDVWSGSG | DSDEFLCLAA |
| HYIDEGWELK | KRVLNFFMVD | PSHSGEMLAE | VIMTCLMEWD | IDRKLFSMAS |
| SHAPPFSENV | ASKIRDRLSQ | NKFLYCYGQL | FDVSCGVNVI | NEMVQDSLEA |
| CCDTINIIRE | SIRYVKSSES | IQDRFNQWIV | ETGAVSERNL | CIDDPMRWDS |
| TCTMLENALE | QKSAFSLMNE | HDPDSVLCPS | DLEWERLGTI | VEFLKVFVEV |
| INAFTKSSCL | PANMYFPEVC | DIHLRLIEWS | KNPDDFISSL | VVNMRKKFDD |
| FWDKNYLVLA | IATILDPRFK | MKLVEYYYPL | FYGTSASELI | EDISECIKLL |
| YDEHSVGSLL | ASSNQALDWQ | NHHHRSNGVA | HGKEPDDRLT | EFDRYINETT |
| TTPGQDSKSD | LEKYLEEPLF | PR**NSDFDILN** | **WWK**VHTPKYP | ILSMMARNVL |
| AVPMLNVSSE | EDAFETCQRR | RVSETWRSLR | PSTVQALMCA | QDWIQSELES |
| S | | | | |

**AtChr1@26346412@26348515 (100%), 81,381.4 Da**
**| potential novel**
**3 exclusive unique peptides, 3 exclusive unique spectra, 4 total spectra, 34/701 amino acids (5% coverage)**

| | | | | |
|---|---|---|---|---|
| DFRFCVTWFR | CLNSNYLVKI | LLFVYMCRRQ | KKLMINNEMD | LSDAVIVKSG |
| RLKSVVWNDF | DRVRKGETYI | AICRHCKKRL | SGSSASGTSH | LRNHLIRCRR |
| RTNGNNNGVA | QYFVKGKKKE | LANERIKDEE | VLSVVNVRYE | HEKEEHEDVN |
| VVSMGLDQRR | CRFDLARMII | LHGYPLSMVE | DVGFRMFIGN | LQPLFELVAF |
| ERVESDCMEI | YAKEKHKIFE | ALDKLPGKIS | ISVDVWSGSG | DSDEFLCLAA |
| HYIDEGWELK | KRVLNFFMVD | PSHSGEMLAE | VIMTCLMEWD | IDRKLFSMAS |
| SHAPPFSENV | ASKIRDRLSQ | NKFLYCYGQL | FDVSCGVNVI | NEMVQDSLEA |
| CCDTINIIRE | SIRYVKSSES | IQDRFNQWIV | ETGAVSERNL | CIDDPMRWDS |
| TCTMLENALE | QKSAFSLMNE | HDPDSVLCPS | DLEWERLGTI | VEFLKVFVEV |
| INAFTKSSCL | PANMYFPEVC | DIHLRLIEWS | KNPDDFISSL | VVNMRKKFDD |
| FWDKNYLVLA | IATILDPRFK | MKLVEYYYPL | FYGTSASELI | EDISECIKLL |
| YDEHSVGSLL | ASSNQALDWQ | NHHHRSNGVA | HGKEPDDRLT | EFDR**YINETT** |
| **TTPGQDSK**SD | LEK**YLEEPLF** | **PRNSDFDILN** | **WWK**VHTPKYP | ILSMMARNVL |
| AVPMLNVSSE | EDAFETCQRR | RVSETWRSLR | PSTVQALMCA | QDWIQSELES |
| S | | | | |

#### No.10. AtChr3@15403874@15404801: Transposable_element_gene

**AtChr3@15403874@15404801 (100%), 35,653.2 Da**
**| potential novel**
**2 exclusive unique peptides, 2 exclusive unique spectra, 2 total spectra, 28/309 amino acids (9% coverage)**

```
AFPVKEYVRM LPYSCSPFGV LYSLDRISCC RTPNGNACEA LDTRSSSTHG
FSLIETWRFD GRSYLSWASE MELFLKQMKL AYVLSEPYPG IGSGSSQDWL
RDDYLCHNHL MKSLSDDLYR QYLKRFKHAK ELWEELKWVY RREESNSKMV
QVRKYIDFKM VEERPVLEQV QEFIKIADSI VGAGMVLDET FYVSTIISKF
PSSWSGFVTR LMEEEFLTVC MLVERLKAEE EFLRSGKKRV TVSAATGSSQ
IEMRPSLGTM HTGSQSVSSK RKEPERDIIP EEKAPKKPNQ MTSSVAEFVD
SETQAKNQH
```

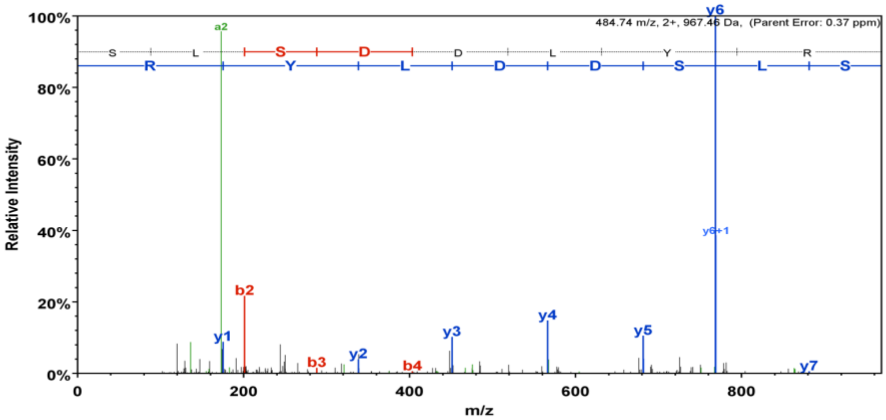


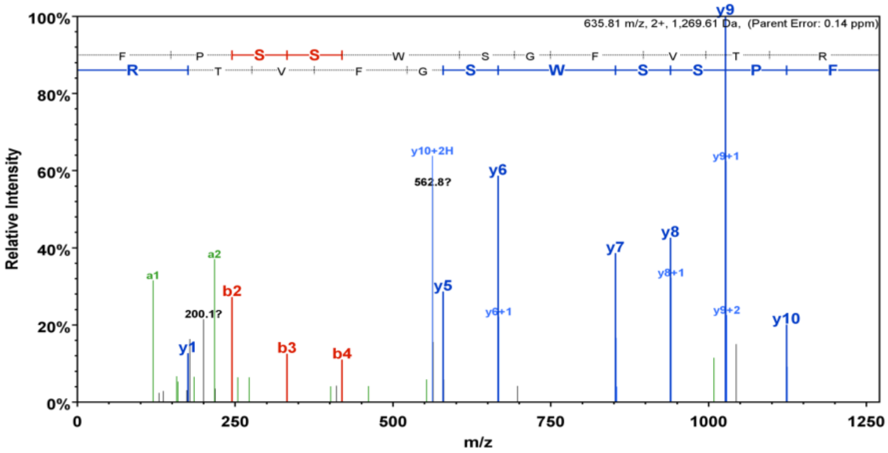


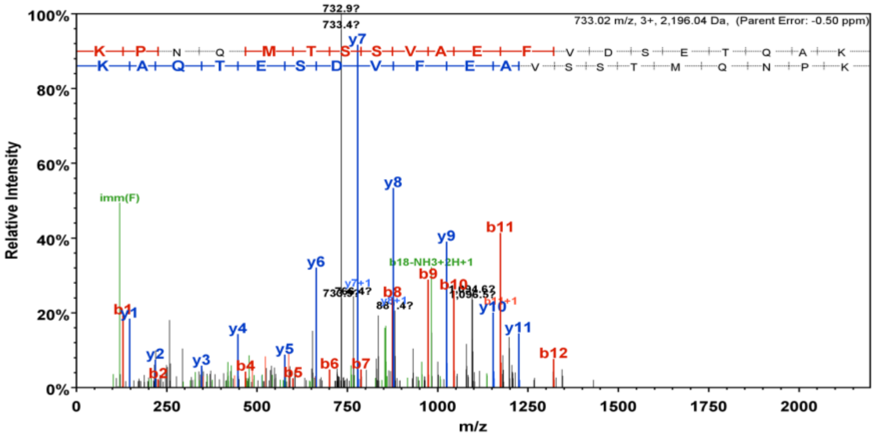

#### No. 11. AtChr5@13104627@13107087: Transposable_element_gene

**AtChr5@13104627@13107087 (100%), 91,881.4 Da**
**| potential novel**
**4 exclusive unique peptides, 4 exclusive unique spectra, 5 total spectra, 66/820 amino acids (8% coverage)**

SISVFNCRSY NIAIGTETTP VAYGNVANVP IQVITGTPEE NNLADINLSS
RKVAPR**VTSE** **SSGLIDIPVT** **ISTIPVGPAK** STSKKFKTKG KNSLVSNITN
LTPKSLKQTF IGSNPGSKSS PPTSLSVVCG VTTSSPRSVS KRRRVMEEPI
ILLQDENVTD TRRRSLRNRG EIRKPVIETD DDDVDADEVD SDEDDSDDVD
ADNVEDDDDK DYVQDIETYY PETEDLDYER EINYSISEAN DGSVESLVAS
WKRCITGVGQ GFESVVEFRD ALQKYAVACR FGYRLRKNES NRACGVCLVG
GCPWK**IYASW** **VPSESVFR**IK KFNRRHTCGG ESWKSAHPK**K** **NWVVGIIK**ER
LQENPNQKTK NIADSFFQDF GIELSYCTIR RGIDEAKGGL HTSFKEAYKY
LPHFVNKVVE ANPGSMVNLV VGEDRRFQRL FLSFQSCIHG FQTGCRPLLF
LDAIPFKSRY HEILLTASAL DGDDCVLPVA LALVDVETDE TWRWFLEQLK
VALSSLRPLT FVSDREKGLV TSVLEIFENA HHGYSIHYLM EDFMRSLR**GP**
**FLGDGKPSLT** **YYLLAAAR**AD RLDGFKVYTE QIRRVSPRAY DWVMQIESKH
WACALFEGEP YSHITSDVAE IYSKWIEEIQ ETSIVQKLVA FVNKIVELVN
SSQEKSKPWF SQLVPSKEES LVEECKKAGS LKVFFCSDTL FEVHDGSVQL
VDISNQTCSC FGWKPTGLPC QHAIAVLNTK GRNLYDYCSS FFTVDSYRLT
YSVALGAVAI DLALVENEGS GKEEDEQVLP PLFSRVQGVE KRIKDRKRGR
SVCCTKCGGV GHNKATCKDD

**AtChr5@13104627@13107087 (100%), 91,881.4 Da**
**| potential novel**
**2 exclusive unique peptides, 2 exclusive unique spectra, 2 total spectra, 37/820 amino acids (5% coverage)**

SISVFNCRSY NIAIGTETTP VAYGNVANVP IQVITGTPEE NNLADINLSS
RKVAPR**VTSE** **SSGLIDIPVT** **ISTIPVGPAK** STSKKFKTKG K**NSLVSNITN**
**LTPK**SLKQTF IGSNPGSKSS PPTSLSVVCG VTTSSPRSVS KRRRVMEEPI
ILLQDENVTD TRRRSLRNRG EIRKPVIETD DDDVDADEVD SDEDDSDDVD
ADNVEDDDDK DYVQDIETYY PETEDLDYER EINYSISEAN DGSVESLVAS
WKRCITGVGQ GFESVVEFRD ALQKYAVACR FGYRLRKNES NRACGVCLVG
GCPWKIYASW VPSESVFRIK KFNRRHTCGG ESWKSAHPKK NWVVGIIKER
LQENPNQKTK NIADSFFQDF GIELSYCTIR RGIDEAKGGL HTSFKEAYKY
LPHFVNKVVE ANPGSMVNLV VGEDRRFQRL FLSFQSCIHG FQTGCRPLLF
LDAIPFKSRY HEILLTASAL DGDDCVLPVA LALVDVETDE TWRWFLEQLK
VALSSLRPLT FVSDREKGLV TSVLEIFENA HHGYSIHYLM EDFMRSLRGP
FLGDGKPSLT YYLLAAARAD RLDGFKVYTE QIRRVSPRAY DWVMQIESKH
WACALFEGEP YSHITSDVAE IYSKWIEEIQ ETSIVQKLVA FVNKIVELVN
SSQEKSKPWF SQLVPSKEES LVEECKKAGS LKVFFCSDTL FEVHDGSVQL
VDISNQTCSC FGWKPTGLPC QHAIAVLNTK GRNLYDYCSS FFTVDSYRLT
YSVALGAVAI DLALVENEGS GKEEDEQVLP PLFSRVQGVE KRIKDRKRGR
SVCCTKCGGV GHNKATCKDD

#### No. 12. AtChr5@13103632@13104208: Transposable_element_gene

**AtChr5@13103632@13104208 (100%), 21,974.1 Da**
**| potential novel**
**2 exclusive unique peptides, 2 exclusive unique spectra, 5 total spectra, 22/192 amino acids (11% coverage)**

```
CNFFLPFRVP QVSVGMGKGK LILICQSGGK FVTDDDGTMT YTGGEAEAID
INHETTFDDF KLKLAKLLNL AYSSLSLKYF LPGNRRTLIT MKQEKDMKRM
YDFHLSSVTA EVFITGQYGF QSEAVLSPGT RYSLLFNHVS CRNFFSLYRF
RSFLFYFNLS LPENVVDVSV LHHLRNNNVM ITDNLLSHLF ES
```

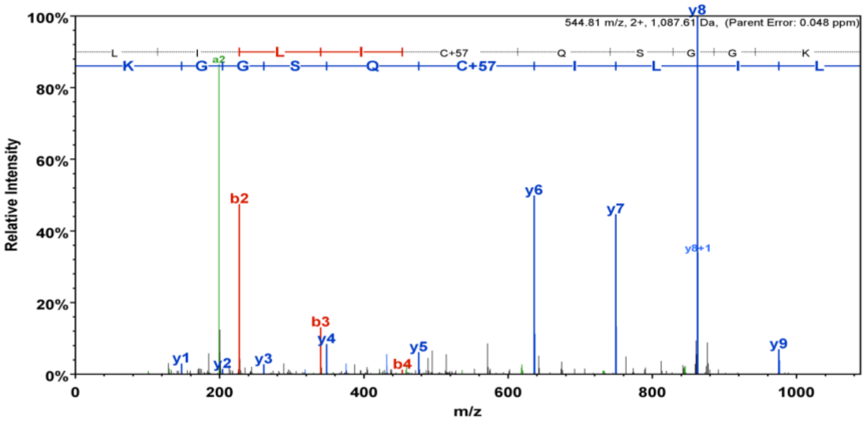


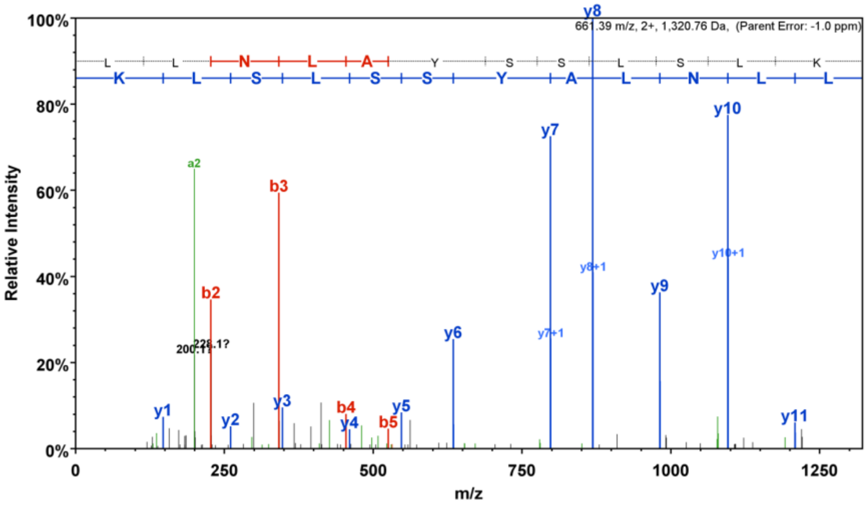

#### No. 13. AtChr3@18791524@18795901: Transposable_element_gene

**AtChr5@19854197@19854566 (100%), 13,673.0 Da**
**| potential novel**
**1 exclusive unique peptides, 1 exclusive unique spectra, 2 total spectra, 15/123 amino acids (12% coverage)**

RFLVMATKK**V** **IAICQSGGEF** **VTNK**DGSLAY SGGDAYAIDI DQDTCMSDFK
SELAENFGFS WENMTLKYFL PGNKKTLITI SKDKDFQRMV SFSADAPNVE
IFVLPEEAEA RNVSNMPASR CVF

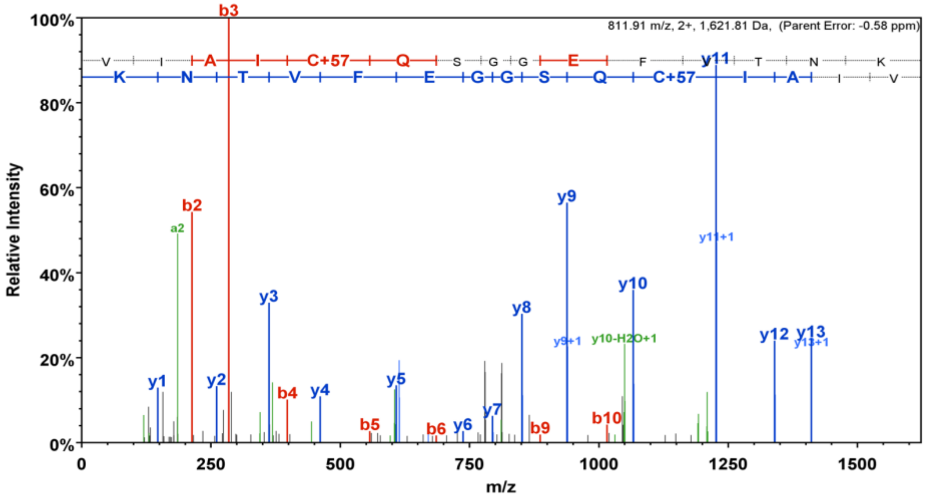


#### No. 14. AtChr3@18791524@18795901: Transposable_element_gene

**AtChr3@18791524@18795901 (100%), 164,046.9 Da**
**| potential novel**
**2 exclusive unique peptides, 2 exclusive unique spectra, 4 total spectra, 35/1459 amino acids (2% coverage)**

TLTGPKMADP YPFPDNVHVS SSVTLKLNDS NYLLWKTQFE SLLSCHKLIG
FVNGGITPPP RTLNVVTGDT SVDVANPQYE SWFCTDQLIR SWLFGTLSEE
VLGYVHNLQT SRDIWISLAE NFNKSSVARE FTLRRTLQLL SKKDKTLSAY
CREFIAVCDA LSSIGKPVDE SMKIFGFLNG LGR**EYDPITT** **VIQSSLSK**IS
PPTFRDVISE VKGFDVK**LQS** **YEESVTANPH** **MAFNTQR**SEY TDNYTSGNRG
KGRGGYGQNR GRSGYSTRGR GFSQHQTNSN NTGERPVCQI CGRTGHTALK
CYNRFDHNYQ SVDTAQAFSS LRVSDSSGKE WVPDSAATAH VTSSTNNLQA
ASPYNGSDTV LVGDGAYLPI THVGSTTISS DSGTLPLNEV LVCPDIQKSL
LSVSKLCDDY PCGVYFDANK VCIIDINTQK VVSKGPRSNG LYVLENQEFV
AFYSNRQCAA SEEIWHHRLG HSNSRILQQL KSSKEISFNK SRMSPVCEPC
QMGKSSKLQF FSSNSRELDL LGRIHCDLWG PSPVVSKQGF KYYVVFVDDY
SRYSWFYPLK AKSDFFAVFV AFQNLVENQF NTKIKVFQSD GGGEFTSNLM
KKHLTDCGIQ HRISCPYTPQ QNGIAERKHR HFVELGLSMM FHSHTPLQFW
VEAFFTASFL SNMLPSPSLG NVSPLEALLK QKPNYAMLRV FGTACYPCLR
PLGEHKFEPR SLQCVFLGYN SQYKGYRCLY PPTGRVYISR HVIFDEETFP
FKQKYQFLVP QYESSLLSAW QSSIPQADQS LIPQAEEGKI ESLAKPPSIQ
KNTIQDTTTQ PAILTEGVLN EEEEEDSFEE TETESLNEET HTQNDEAEVT
VEEEVQQEPE NTHPMTTRSK AGIHKSNTRY ALLTSKFSVE EPKSIDEALN
HPGWNNAVND EMRTIHMLHT WSLVQPTEDM NILGCRWVFK TKLKPDGSVD
KLKARLVAKG FHQEEGLDYL ETFSPVVRTA TIRLVLDVAT AKGWNIKQLD
VSNAFLHGEL KEPVYMLQPP GFVDQEKPSY VCRLTKALYG LKQAPRAWFD
TISNYLLDFG FSCSKSDPSL FTYHKNGKTL VLLLYVDDIL LTGSDHNLLQ
ELLMSLNKRF SMKDLGAPSY FLGVEIESSP EGLFLHQTAY AKDILHQAAM
SNCNSMPTPL PQHIENLNSD LFPEPTYFRS LAGKLQYLTI TRPDIQFAVN
FICQRMHSPT TADFGLLKRI LRYVKGTIHL GLHIKKNQNL SLVAYSDSDW
AGCKETRRST TGFCTLLGCN LISWSAKRQE TVSKSSTEAE YRALTAVAQE
LTWLSFLLRD IGVTQTHPTL VKCDNLSAVY LSANPALHNR SKHFDTDYHY
IREQVALGLV ETKHISATLQ LADIFTKPLP RRAFIDLRIK LGVAEPPTTS
LRGNVSEMSE AKEMGQTHNK SSPTIKPGQQ IQKMTKIKSC LSSVHVPAGN
IERERKTLL

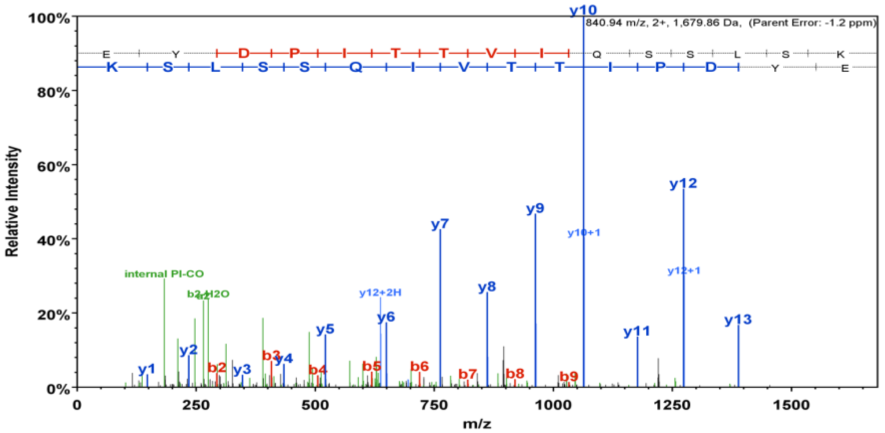

840.94 m/z, 2+, 1,679.86 Da, (Parent Error: -1.2 ppm)
E Y D P I T T V I Q S S L S K
K S L S S Q I V T T I P D Y E
Relative Intensity
100%
80%
60%
40%
20%
0%
m/z
0
250
500
750
1000
1250
1500
y10
y12
y9
y10+1
y7
y12+1
internal PI-CO
b2-H2O
y12+2H
y8
y6
y5
y11
y13
y1
y2
b2
y3
b3
y4
b4
b5
b6
b7
b8
b9


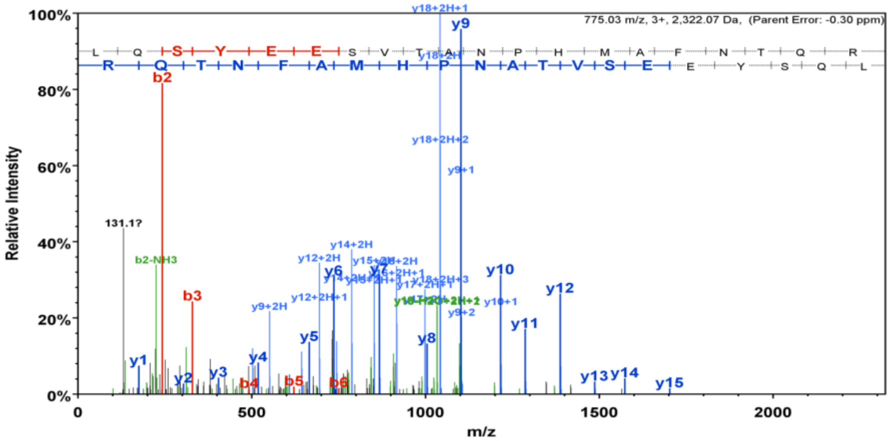

775.03 m/z, 3+, 2,322.07 Da, (Parent Error: -0.30 ppm)
L Q S Y E E S V T A N P H M A F N T Q R
R Q T N F A M H P N A T V S E E Y S Q L
Relative Intensity
100%
80%
60%
40%
20%
0%
m/z
0
500
1000
1500
2000
y18+2H+1
y9
b2
y18+2H+2
y9+1
131.1?
b2-NH3
y14+2H
y12+2H
y10
b3
y12
y9+2H
y9+2
y10+1
y11
y5
y4
y8
y1
y2
y3
b4
b5
b6
y13
y14
y15

No. 15. AtChr4@7689781@7690192@0: Novel_transcribed_region

**AtChr4@7689781@7690192@0 (100%), 16,144.3 Da**
**| potential novel**
**1 exclusive unique peptides, 1 exclusive unique spectra, 9 total spectra, 41/137 amino acids (30% coverage)**

I R E M L R L I F F  I L V T A I C S T K  T E A C E K **N A I V**  **I I N D L G P G R** I  L Q Y H C R Y T K K
D L G V Q H L N F H  A I K T I H L Q D E  G K N I T K W H C L  F K Q G L N M R Y R  **N Y V E A Y S Q N T**
**E A P Q C G Q V R** V  W T A R M S G I W F  K **I S Y N N P S G R**  Y I R G W K Y

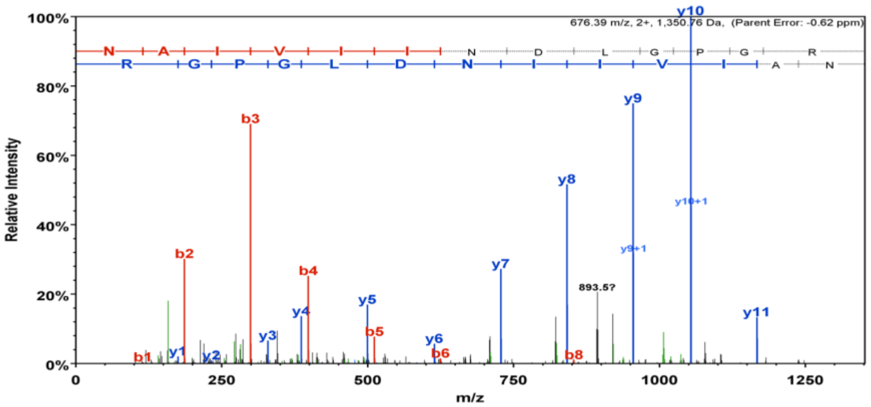


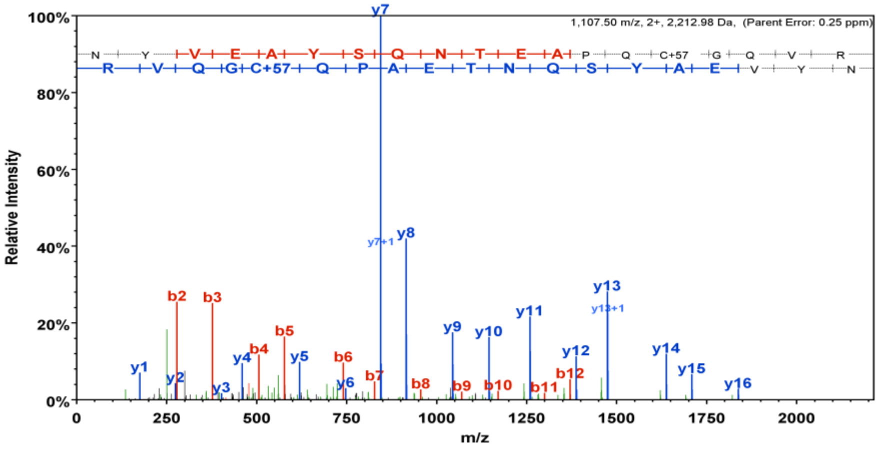


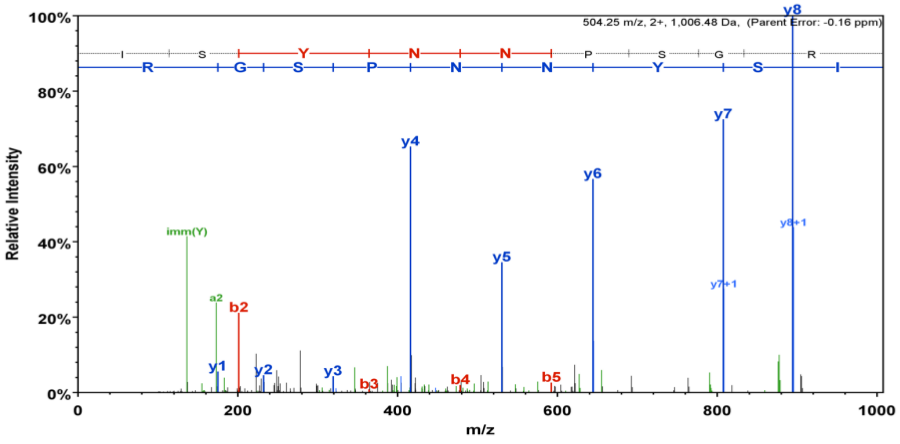

### No. 16. AtChr5@10277623@10277968: Novel_transcribed_region

**AtChr5@10277623@10277968 (100%), 13,016.9 Da**
**| potential novel**
**2 exclusive unique peptides, 2 exclusive unique spectra, 4 total spectra, 22/115 amino acids (19% coverage)**

NKTISYSITL SFLIATNLFM LNMKTIPLMF IALIILSISF LAPIKAQGIV
GCDVELDKCV VMR**NNQQWAL LYNTCCK**KFK KPGPQPCMCL FFKYPTPK**ET**
**ALSLLR**YCKA PIPKC

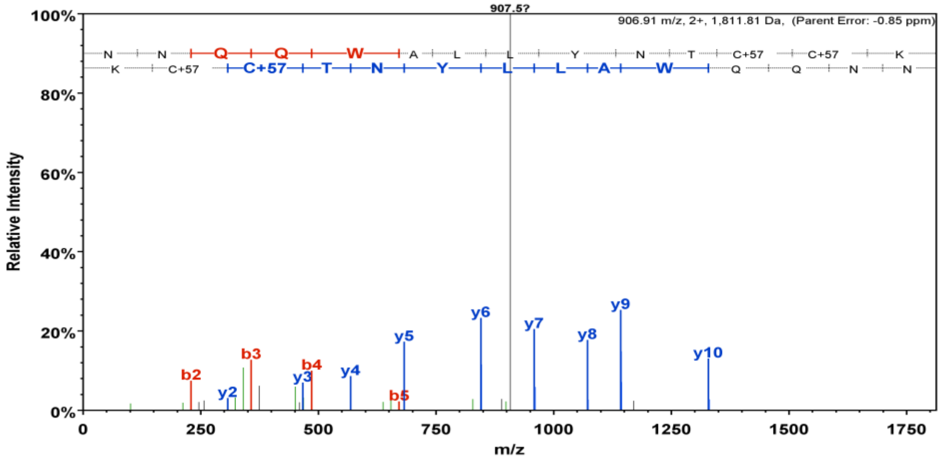


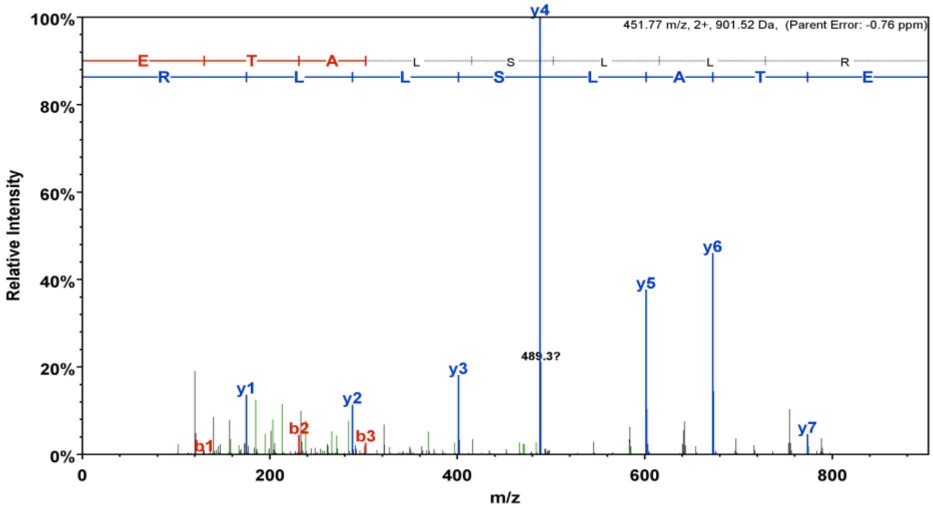

#### No.17. AtChr1@10428741@10429146: Antisense_long_noncoding_rna

**AtChr1@10428741@10429146 (100%), 15,336.7 Da**
**| potential novel**
**3 exclusive unique peptides, 3 exclusive unique spectra, 3 total spectra, 29/135 amino acids (21% coverage)**

ANSRKILTLF PFSLSTQKKR QLSKNTDSTV SMPESSSSFS SLFLLLIIFF
LLLLSPSLSS PESEVHVLDR ELLEIQTNPK PGKTNRKLKC CEMRSR**SQCS**
**AFPR**CRWCR**S** **EALDDLCFSK** AEALR**LPSQV** **FLCEL**

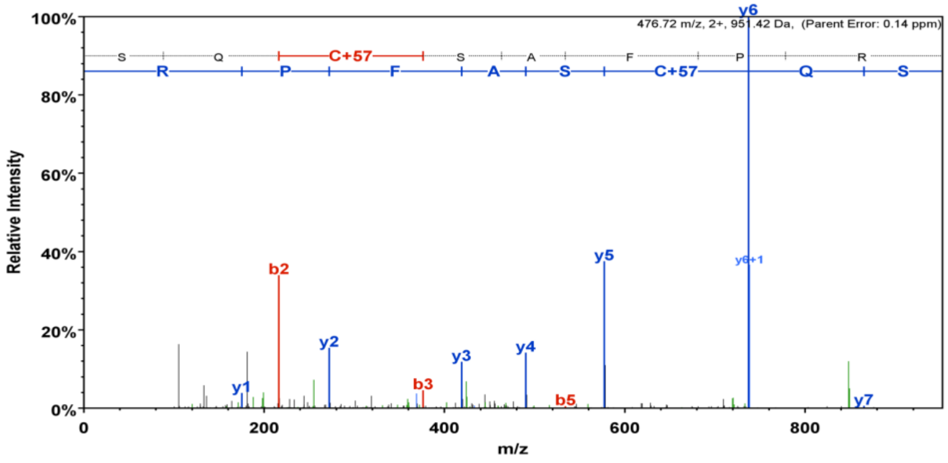


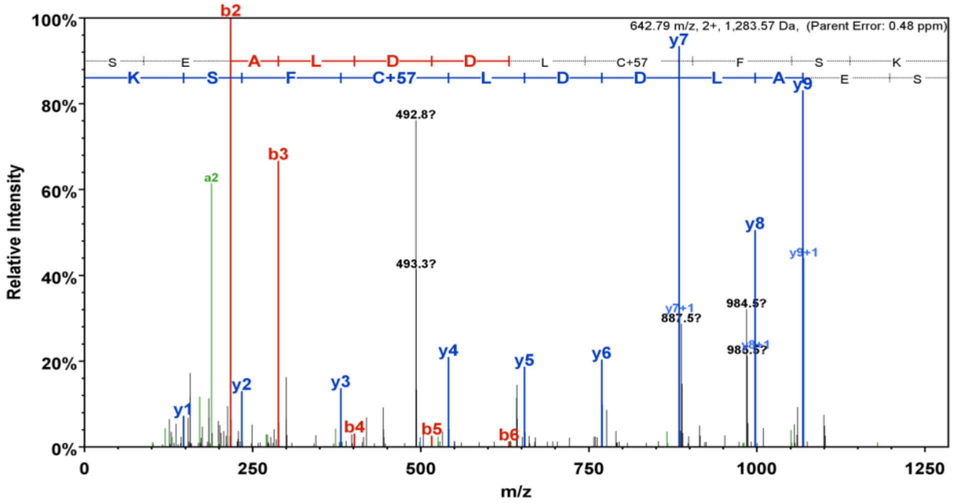

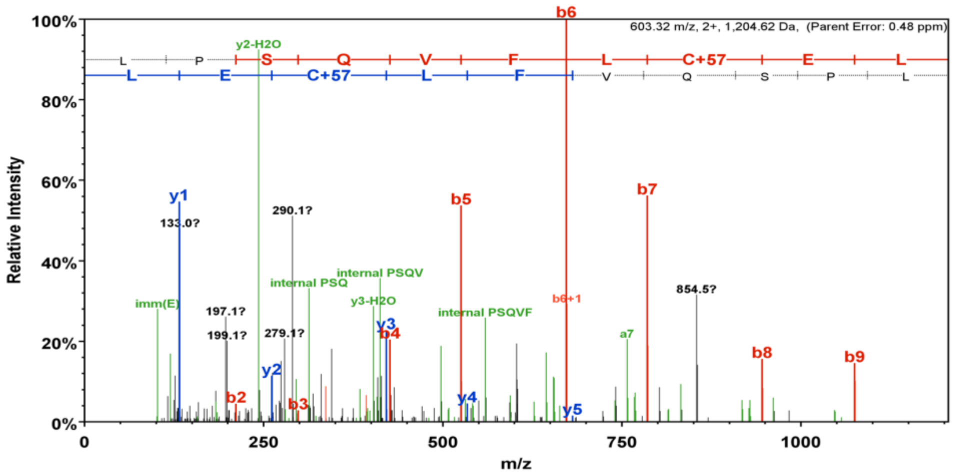
603.32 m/z, 2+, 1,204.62 Da, (Parent Error: 0.48 ppm)
Relative Intensity
m/z
L P S Q V F L C+57 E L
L E C+57 L F V Q S P L
b6
y2-H2O
y1
133.0?
290.1?
b5
b7
854.5?
internal PSQV
internal PSQ
y3-H2O
imm(E)
197.1?
199.1?
279.1?
internal PSQVF
b6+1
a7
b8
b9
y3
b4
y2
b2
b3
y4
y5

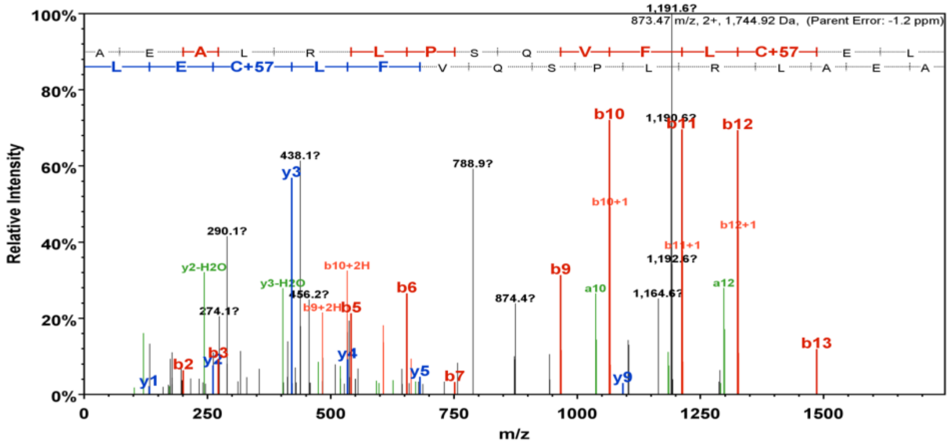
1,191.6?
873.47 m/z, 2+, 1,744.92 Da, (Parent Error: -1.2 ppm)
Relative Intensity
m/z
A E A L R L P S Q V F L C+57 E L
L E C+57 L F V Q S P L R L A E A
b10
1,190.6?
b11
b12
438.1?
y3
788.9?
b10+1
b12+1
290.1?
b11+1
1,192.6?
y2-H2O
b10+2H
y3-H2O
b9
a10
a12
456.2?
b6
274.1?
b9+2H
b5
874.4?
1,164.6?
b13
b2
y2
b3
y4
y5
b7
y9
y1

**No. 18. AtChr4@9188288@9188534: Long_noncoding_rna**

**AtChr4@9188288@9188534 (100%), 9,087.1 Da**
**| potential novel**
**2 exclusive unique peptides, 2 exclusive unique spectra, 5 total spectra, 27/82 amino acids (33% coverage)**

F D I V G F E S E K   I P C C S L V G K **V**   **G G L I P C G P P S**   **K** V C M D R S K Y V   F W D P Y H P T E A
A N I I I A R R **L L**   **S G D T S D I Y P I**   **N L R** Q L A N L K I   Y A

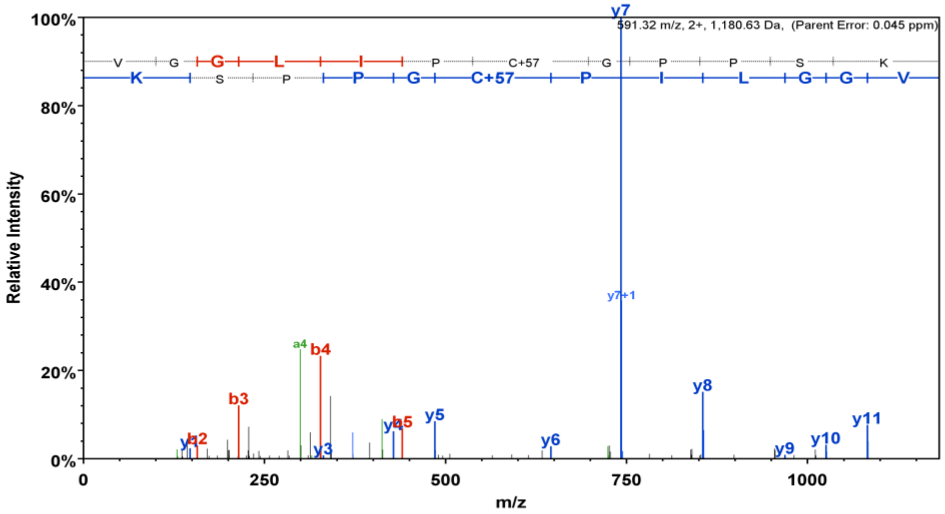

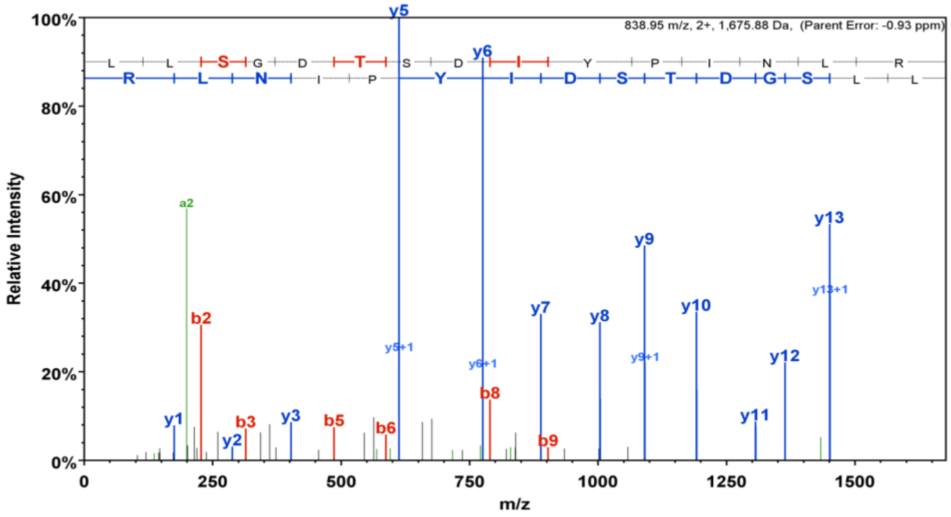

#### No. 19. AtChr3@3253541@3253958: Long_noncoding_rna

**AtChr3@3253541@3253958 (100%), 16,526.7 Da**
**| potential novel**
**2 exclusive unique peptides, 2 exclusive unique spectra, 3 total spectra, 27/139 amino acids (19% coverage)**

```
F K I M N L V F S F   L F F I F V L C C V   S K E D K C H K N T   V A F Q N N L F Q S   H S I L K V H C K S
R S D D L G E H F V   K F Q D T A Y N F S   F H D N L L F T T I   F K C N L W K G A R   M E Y H R N I T A Y
E G D L L Y R C G A   L Y S W D I R D D A   I Y L S E N D N P E   K L M Y S W I K G
```

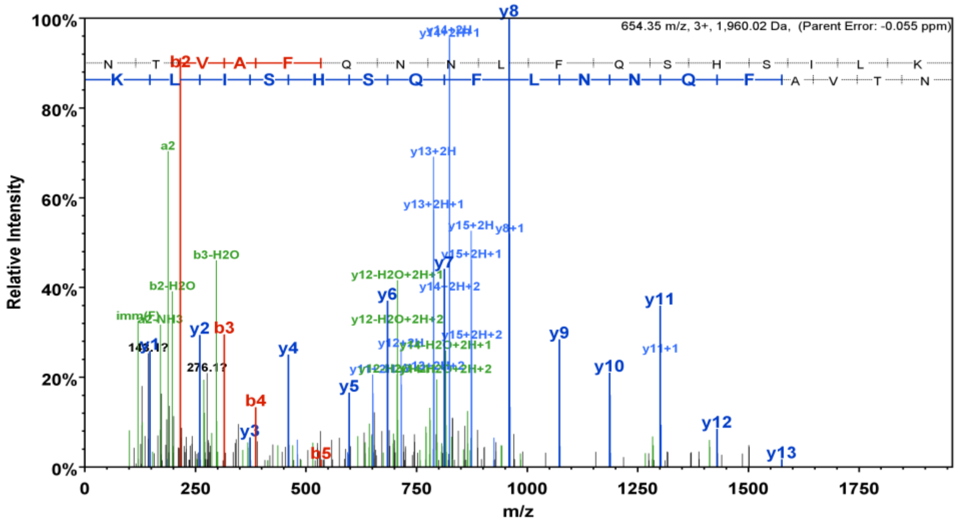


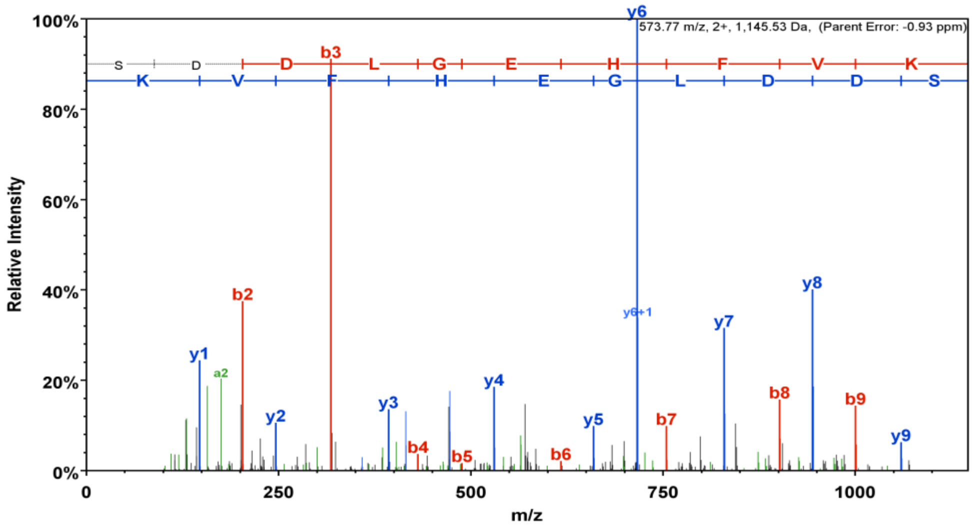

**No. 20. AtChr5@26518786@26519188: Arabidopsis thaliana drought-induced 8**

**AtChr5@26518786@26519188 (100%), 14,782.2 Da**
**| potential novel**
**3 exclusive unique peptides, 3 exclusive unique spectra, 8 total spectra, 29/134 amino acids (22% coverage)**

LSSKCLLKTR RTWRLTRTVQ EVRPLTSTET RSSSSMTSTE IRWEEEDTEL
VVVEELQVAK DTEQVAKGTD QVAK**GTEPVA** **KDTEPGPGLK** **ALELAEELGT**
**TAK**SNSTRKV VVAWEECFTA PDLDPALARY VSSV

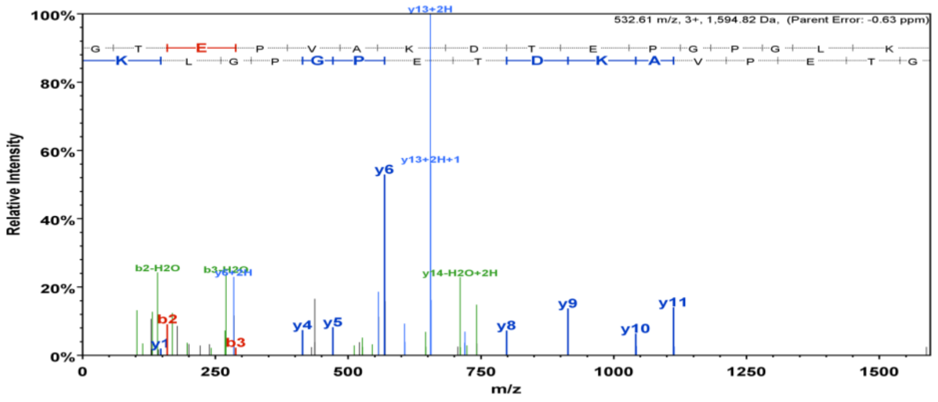


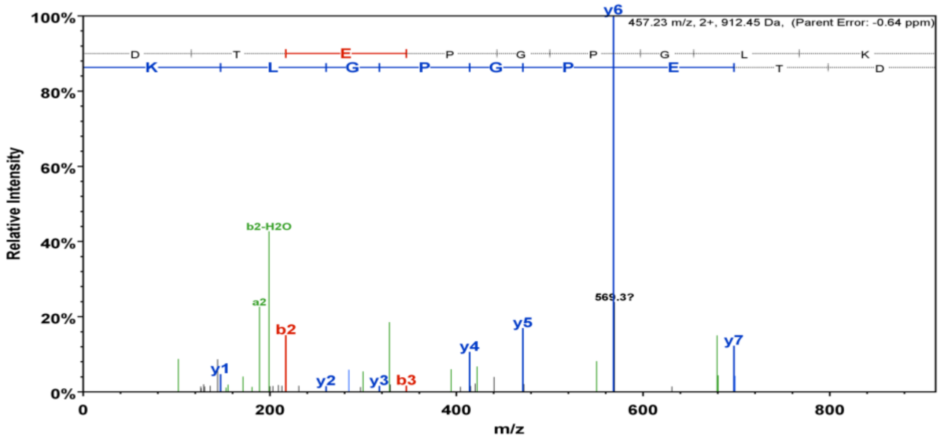


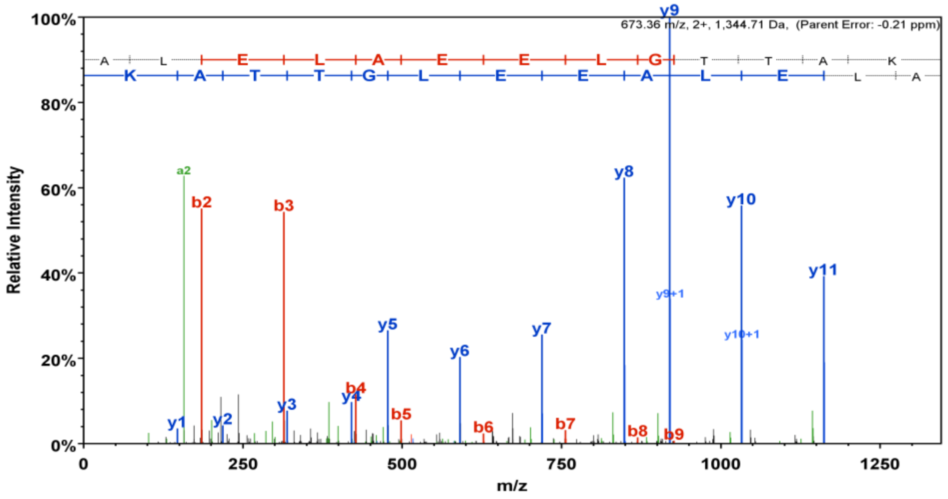

#### No. 21. AtChr2@17714928@17715315: Late embryogenesis abundant 25

**AtChr2@17714928@17715315 (100%), 14,747.1 Da**
**| potential novel**
**2 exclusive unique peptides, 2 exclusive unique spectra, 4 total spectra, 27/129 amino acids (21% coverage)**

L I V V W W C F V L  L K K L C F G F V L  R E V R K K **S N Q G**  **L S Q E Q G E T L E**  **K** S T G K K E G V R
K M S T A M E Q K **E**  **Y P E K G G E M L G**  **R** Q S T E E G V R K  M S S D M D Q K A R  S K K Q G E T L E K
N T E E E E A V R D  M L K K K G L E R E  G C L G Q S A R L

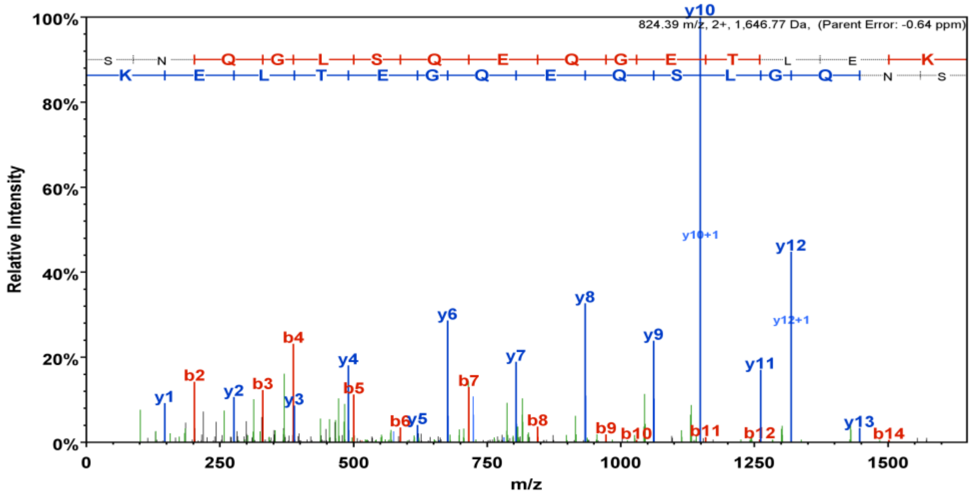


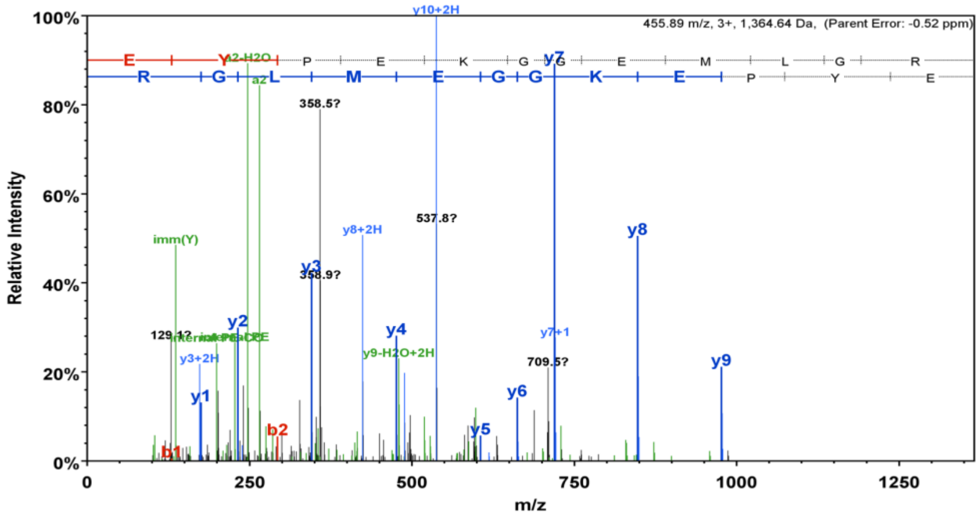

#### No. 22. AtChr2@2056255@2056639: Glycine-rich protein family

**AtChr2@2056255@2056639 (100%), 13,590.1 Da**
**| potential novel**
**2 exclusive unique peptides, 2 exclusive unique spectra, 2 total spectra, 25/128 amino acids (20% coverage)**

R A A V A A A K E D   I R A A E V V V A K   E D T K **A V E V V A   A K E D T K A V E V   V D K E A V D T K** A
A V V A K E D T K A   V E E V V M L V E A   A E E V E V E V V G   V V E A E E E V V V   E V A V E E V E A V
G C R I S L P I L A   S F S L N A K C L R   I L N Y N K V T

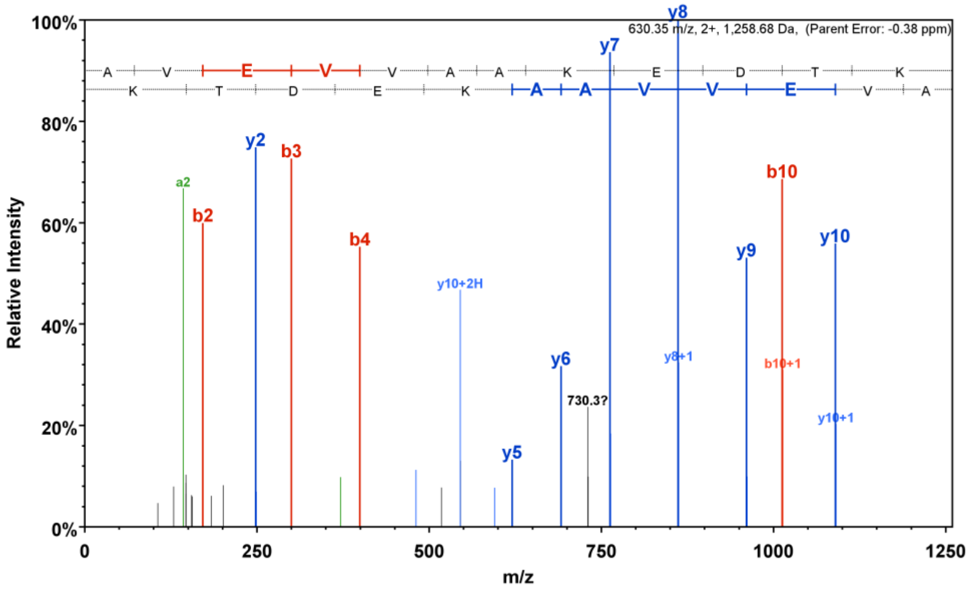


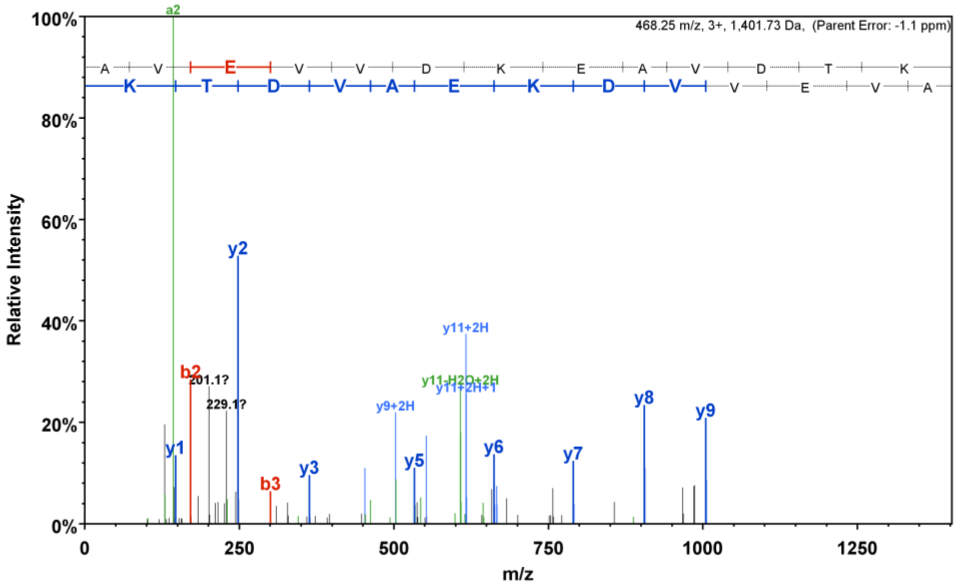

**No. 23. AtChr4@6693384@6693678: Plant self-incompatibility protein S1 family**

**AtChr4@6693384@6693678 (100%), 10,554.5 Da**
**| potential novel**
**3 exclusive unique peptides, 5 exclusive unique spectra, 7 total spectra, 24/98 amino acids (24% coverage)**

```
LLREIDKFTL IERYIHIHRQ RREMAGPAQA AKQSSEVLGQ RKSLGICPLR
AAAVGAVIIG GIGYVVLYSK KKPEASAGDV AKVMSGVGGT PENTRPRN
```

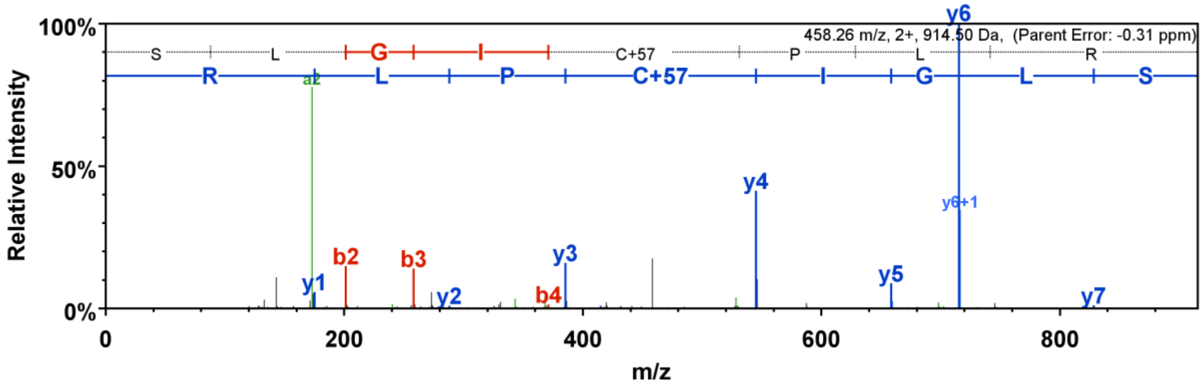


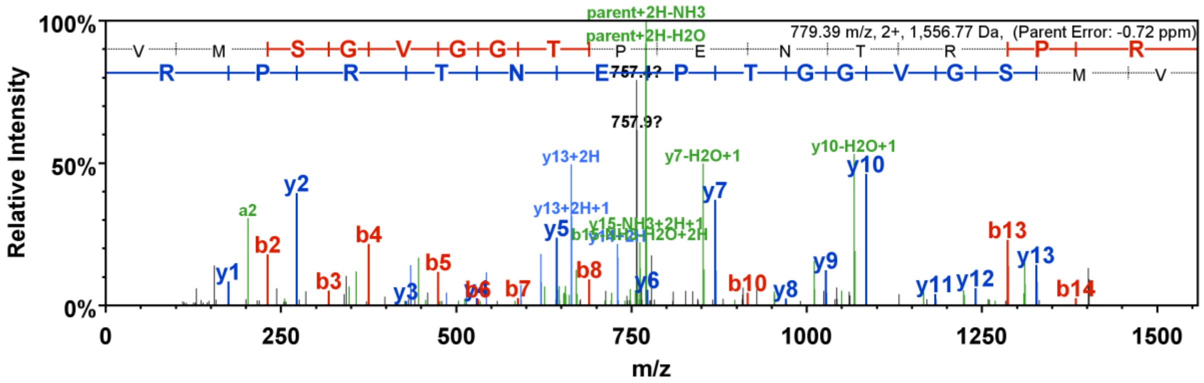


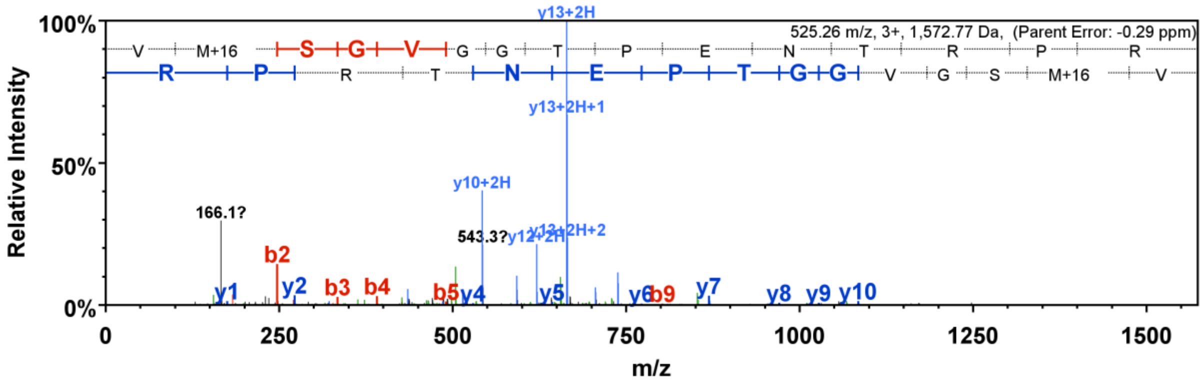

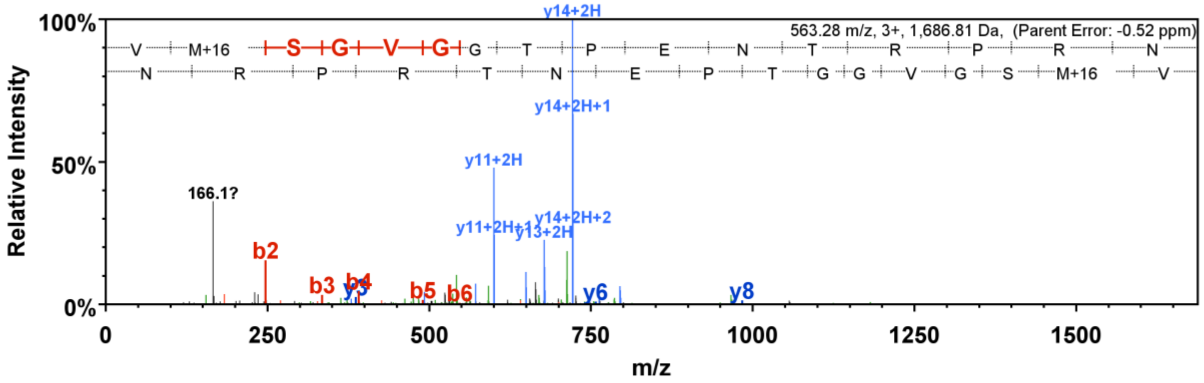


**No. 24. AtChr5@7534553@7534787: Arabidopsis histone deacetylase 2**

Peptide (SPNIEQYPFFDRFD) is the novel identification, it is not in AT5G22650.1 from TAIR10 and TAIR11.

**AT5G22650.1 (100%), 32,347.6 Da**
**| Symbols: HD2B, HDT02, HDT2, ATHD2B, HDA4, HD2, ATHD2 | histone deacetylase 2B | chr5:7534120-7536054 FORWARD L**
**3 exclusive unique peptides, 5 exclusive unique spectra, 52 total spectra, 98/306 amino acids (32% coverage)**

```
MEFWGVAVTP KNATKVTPEE DSLVHISQAS LDCTVKSGES VVLSVTVGGA
KLVIGTLSQD KFPQISFDLV FDKEFELSHS GTKANVHFIG YKSPNIEQDD
FTSSDDEDVP EAVPAPAPTA VTANGNAGAA VVKADTKPKA KPAEVKPAEE
KPESDEEDES DDEDESEEDD DSEKGMDVDE DDSDDDEEED SEDEEEEETP
KKPEPINKKR PNESVSKTPV SGKKAKPAAA PASTPQKTEE KKKGGHTATP
HPAKKGGKSP VNANQSPKSG GQSSGGNNNK KPFNSGKQFG GSNNKGSNKG
KGKGRA
```

**AtChr5@7534553@7534787 (100%), 8,603.8 Da**
**| potential novel**
**1 exclusive unique peptides, 1 exclusive unique spectra, 42 total spectra, 70/78 amino acids (90% coverage)**

```
ASLDCTVKSG ESVVLSVTVG GAKLVIGTLS QDKFPQISFD LVFDKEFELS
HSGTKANVHF IGYKSPNIEQ YPFFDRFD
```

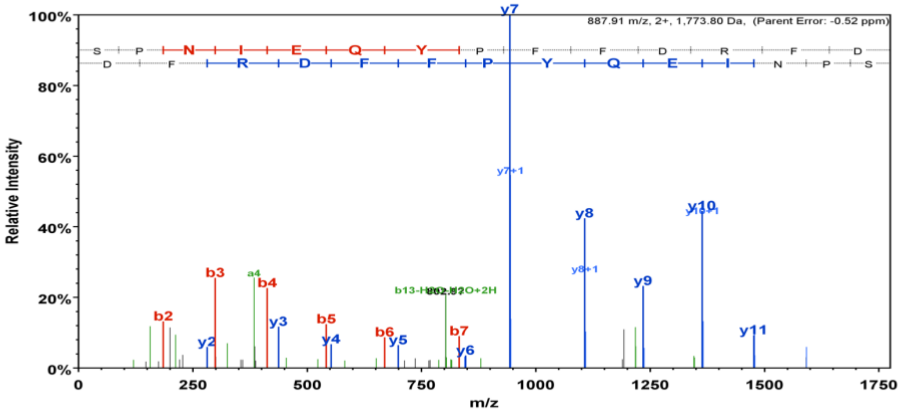

**No. 25. AtChr1@20390129@20390324: Acyl carrier protein 3**

Peptide (QAKPETVDKVCAVVR) is the novel identification. It is not present in AT1G54630.1 from TAIR10 and TAIR11.

**AT1G54630.1 (100%), 14,650.2 Da**
**| Symbols: ACP3 | acyl carrier protein 3 | chr1:20401642-20402919 REVERSE LENGTH=136**
**1 exclusive unique peptides, 1 exclusive unique spectra, 36 total spectra, 37/136 amino acids (27% coverage)**

M A S I A T S A S T S L Q A R P R Q L V I G A K Q V K S F S Y G S R S N L S F N L R Q L P T R L T V
Y C A A K P E T V D K V C A V V R **K Q L** **S L K E A D E I T A** **A T K** F A A L G A D S L D T V E I V M G
L E E E F G I E M A E E K **A Q S I A T V** **E Q A A A L I E E L** **L L E K** A K

**AtChr1@20390129@20390324 (100%), 7,064.8 Da**
**| potential novel**
**1 exclusive unique peptides, 1 exclusive unique spectra, 21 total spectra, 31/65 amino acids (48% coverage)**

L G S L I R L G D W L I Y I S I F L L L C K **Q A K P E T V D** **K V C A V V R K Q L** **S L K E A D E I T A**
**A T K** F A A L G A D S L D T V

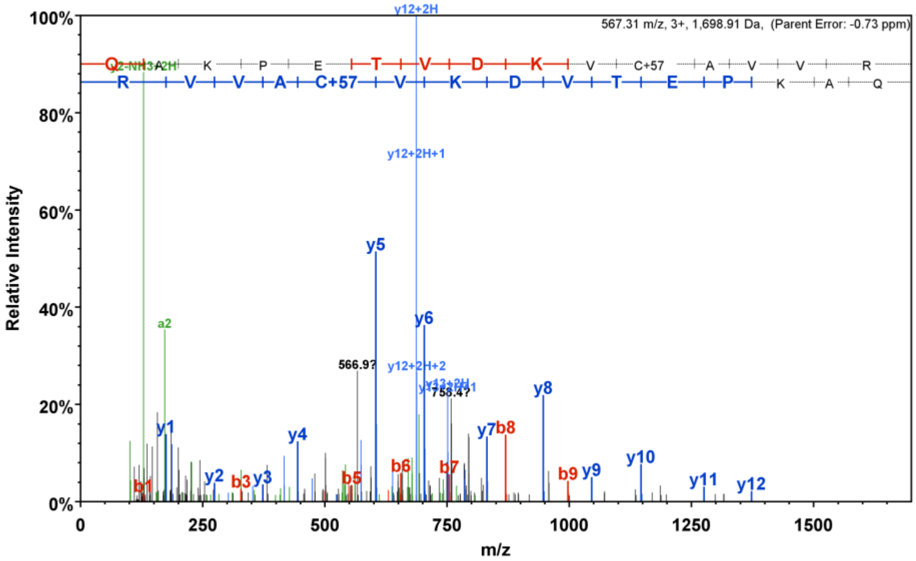

#### No. 26. AtChr1@18512092@18512548: DNA/RNA polymerases superfamily protein

**AtChr1@18512092@18512548 (85%), 16,579.7 Da**
**| potential novel**
**1 exclusive unique peptides, 1 exclusive unique spectra, 6 total spectra, 9/152 amino acids (6% coverage)**

RNPEEMKMAL SPPKLILSPS NSLGDSNSLP SMSDILTSSK ARKLDLK**IQT**
**LGPFFR**VTGK NADTGGEVGR AEGVVRPWFG RGLVLHLDTI RLTKETVAMD
KSVLGVGLYV GAVAIRHGYD CGCRTAQLLA IYDSDLYHSK VSLSLHNSFY
RN

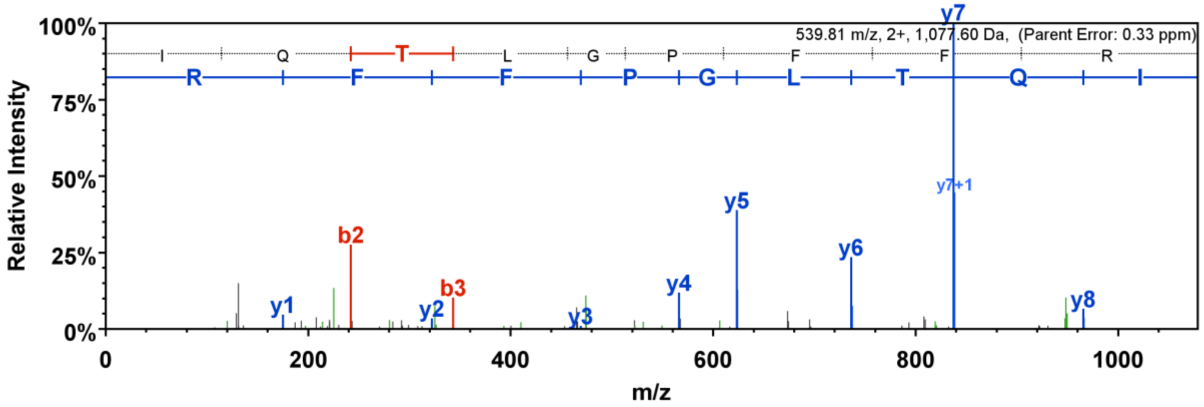


#### No. 27. AtChr2@15278911@15279358: Transcriptional activator (DUF662)

Peptide (STEDSSSCFGLR) is the novel identification. It is not present in AT2G36410.1 from TAIR10 or TAIR11.

**AT2G36410.1 (100%), 22,347.1 Da**
**| Symbols: | Family of unknown function (DUF662) | chr2:15279041-15280270 FORWARD LENGTH=195**
**2 exclusive unique peptides, 2 exclusive unique spectra, 10 total spectra, 55/195 amino acids (28% coverage)**

MTQSQTNDGA GAGAVTTVES VPPQPQSQPQ PQPQQQSNEM VLHTGSLSFS
SHMSREDEEM TRSALSAFRA KEDEIEKRRM EVRERIQAQL GRVEQETKR**L**
**STIREELESM** **ADPMRK**EVSV VRKKIDSVNK **ELKPLGSTVQ** **K**KER**EYKEAL**
**DTFNEK**NREK VQLITK**LMEM** **EQLVGESEKL** **R**MIKLEELSK SIETV

**AtChr2@15278911@15279358 (100%), 16,765.7 Da**
**| potential novel**
**1 exclusive unique peptides, 1 exclusive unique spectra, 4 total spectra, 25/149 amino acids (17% coverage)**

NTPKSAKIGL IEEPHRK**STE** **DSSSCFGLR**K KNSKNLKEKK KSKMTQSQTN
DGAGAGAVTT VESVPPQPQS QPQPQPQQQS NEMVLHTGSL SFSSHMSRED
EEMTRSALSA FRAKEDEIEK RRMEVRER**IQ** **AQLGRVEQET** **K**RLSTIREV

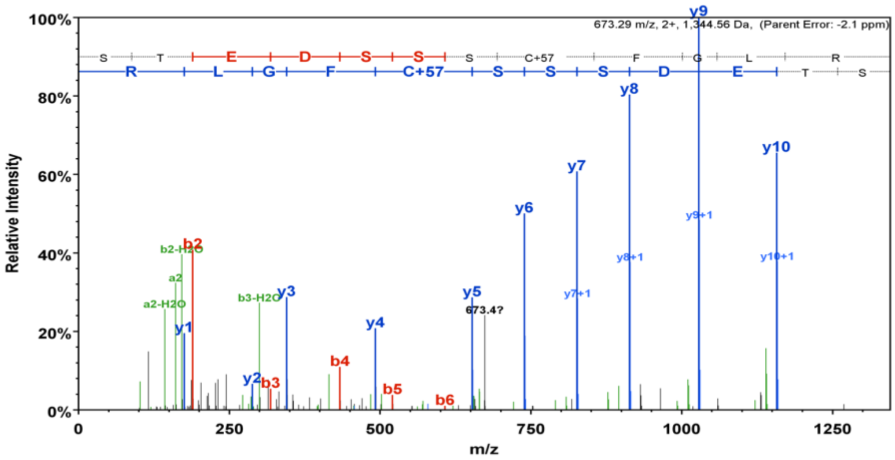

673.29 m/z, 2+, 1,344.56 Da, (Parent Error: -2.1 ppm)
Relative Intensity
m/z


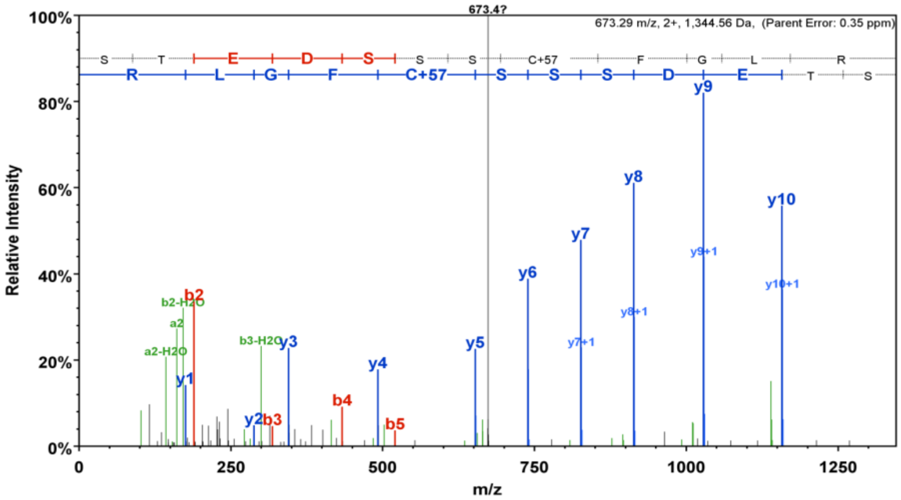

673.29 m/z, 2+, 1,344.56 Da, (Parent Error: 0.35 ppm)
Relative Intensity
m/z

No. 28. AtChr5@21376666@21377527: DNA/RNA polymerases superfamily protein

Peptide (IDPLIIQGLDGK) is the novel identification. It is not present in AT5G52710.1 from TAIR10 or TAIR11.

**AT5G52710.1 (100%), 48,127.5 Da**
**| Symbols: | Copper transport protein family | chr5:21375753-21377841 FORWARD LENGTH=423**
**5 exclusive unique peptides, 5 exclusive unique spectra, 13 total spectra, 131/423 amino acids (31% coverage)**

```
MSNPHYNLKT QEKTAEFKLD VSDENIRRNT MKIVWMFPVT FVDIKEKGIL
KVRGKFDHIE MREILQQHID ESVDLINPMK KSEKKQIPGL ATVFNYFKTP
ATQEIVVFEF RVLDERIIPK AMEVIWEFPV TSVEIENDFY LKVKGGEINK
EVMKTQLKDV DKHVKIKWHG QDVEEEPKKQ SEAGGSKLQP EQLSKKKNIY
EDAYGEHLKK RQGINQAKQP AFYDDWLLEE AHGPFNQQQG FKQENLLPIY
GGAHAQGLYK QNANKVFQPQ PKGGFAALNL GEKKKQAEVK KQNQDAGWFG
NMFGNQQPQV KQEFRGKREN QHAQRIDPLI IQEFGSSSTR NRSSFSRSSS
AMDLSQTTSS SSVAYSSYST TPNSSRNSSH SRSRSSENLS QFQSNSTGNS
YTSSQSNTAS GSKQGQSSKK PGK
```

**AtChr5@21376666@21377527 (100%), 32,697.2 Da**
**| potential novel**
**1 exclusive unique peptides, 1 exclusive unique spectra, 15 total spectra, 56/287 amino acids (20% coverage)**

```
INPCLLLLLL LLVVTSVEIE NDFYLKVKGG EINKEVMKTQ LKDVDKHVKI
KWHGQDVEEE PKKQSEAGGS KLQPEQLSKK KNIYEDAYGE HLKKRQGINQ
AKQPAFYDDW LLEEAHGPFN QQQGFKQENL LPIYGGAHAQ GLYKQNANKV
FQPQPKGGFA ALNLGEKKKQ AEVKKQNQDA GWFGNMFGNQ QPQVKQEFRG
KRENQHAQRI DPLIIQGLDG KFHAYKEKNN NPYARSSNGP TSNNGTSQNL
GINKLVSVYQ IYFYGITFKI KFLLLFVVDL TILAFFC
```

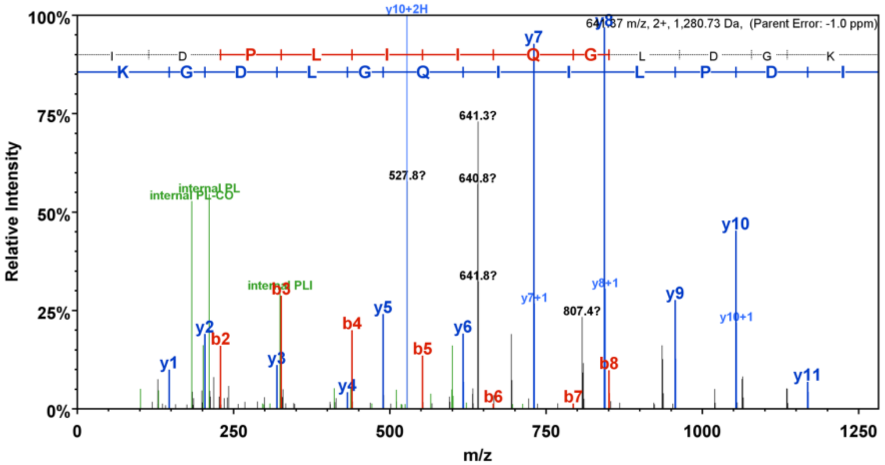
