## Supplementary File 2 for "Arabidopsis Proteome and the Mass Spectral Assay Library"

**Table 3. The list of novel proteins identified using proteogenomics approach. These proteins/sequences were either not present in TAIR10 or incomplete, but present in the latest TAIR11 proteome database**. The detailed information about the novel identification, its annotation from TAIR webpage and their matched peptides (highlighted in yellow) were presented following with this table.

Furthermore:

- If the spectral counts are smaller than 4, the matched spectra were also presented.
- If the novel identification shares with TAIR10 protein entry a group of peptides, then the matched TAIR10 protein entry were included, and the unique peptide(s) associated with the novel identification were specified.
- If different set of peptides (from different tissues) were associated with the novel identification, all sequence matches were listed.

| No | 6ORF ID | Genomics Locus sequence (DNA) | TAIR annotation | Notes (in Tair11) | Total Spectral Count | | | | | | | | | |
|---|---|---|---|---|---|---|---|---|---|---|---|---|---|---|
| | | | | | Cotyle-done | Rosette | Cauline | Stem | Pollen grain | Flower | Root | Silique | Root cells | Seeds |
| 1 | Atchr1@19256097@19256652 | AT1G51850.2 | Leucine-rich repeat protein kinase family protein | New isoform | | | | | | | 2 | | | |
| 2 | AtChr5@18590218@18590728 | AT5G45830.4 | Delay of gemination 1 | New isoform | | | | | | | | 3 | | 31 |
| 3 | AtChr3@1743539@1745543 | AT3G05850.1 | MuDR family transposase | New entry | 7 | 1 | 2 | 1 | 14 | 13 | 15 | 6 | 29 | 1 |
| 4 | AtChr3@1745505@1745964 | AT3G05850.1 | MuDR family transposase | New entry | | | | | 1 | 2 | | 1 | 2 | |
| 5 | AtChr5@20952547@20953303 | AT5G51585.1 | Transmembrane protein | New entry | | | | | | | | | 2 | |
| 6 | AtChr3@6174451@6174928 | AT3G18040.4 | MAP kinase 9 | New entry | | | | | 1 | 1 | 1 | | | |
| 7 | AtChr4@9187917@9188217 | AT4G16233.1 | GDSL-like lipase/acylhydrolase superfamily protein | New entry | | | | | | | 4 | | 4 | |
| 8 | AtChr3@3252605@3253004 | AT3G10455.1 | Plant self-incompatibility protein S1 family protein | New entry | | | | | 3 | 6 | | | | |
| 9 | AtChr5@4376549@4378589 | AT5G13590.1 | Unknown protein | AA correction | 2 | | 1 | | 2 | | 1 | | 1 | |
| 10 | AtChr4@15093731@15095657 | AT4G30990.3 | ARM repeat superfamily protein | Sequence extension | | | | | | | | | 2 | |
| 11 | AtChr4@13558438@13559653 | AT4G27010.2 | Ribosome 60S biogenesis amino-terminal protein | Sequence extension | | | | | | | | | 2 | |
| 12 | AtChr4@7455224@7455494 | AT4G12610.1 | Transcription initiation factor IIF subunit alpha RAP74 | Sequence extension | | | | 1 | 2 | 5 | 4 | 3 | 6 | 4 |
| 13 | AtChr4@1344112@1344550 | AT4G03050.1 | 2-oxoglutarate-dependent dioxygenase | Sequence extension | | | | | | | | 40 | | 19 |
| 14 | AtChr5@15331919@15332135 | AT5G38360.1 | Alpha/beta-Hydrolases superfamily protein | Sequence extension | 3 | 1 | 3 | | 1 | 1 | | 1 | | |

**No.1. Atchr1@19256097@19256652: Leucine-rich repeat protein kinase family protein**

Peptide (TNSLQVCLIK) is not in TAIR10 but present in TAIR 11.

**AT1G51805.1 (100%), 98,078.5 Da**
**| Symbols: | Leucine-rich repeat protein kinase family protein | chr1:19221187-19225590 REVERSE LENGTH=884**
**18 exclusive unique peptides, 20 exclusive unique spectra, 78 total spectra, 293/884 amino acids (33% coverage)**

MESHRVFVAT FMLILHLVQA QDQPGFINVD CGLLPRDSPY NALGTGLVYT
SDVGLVSSGK TGKIAKEFEE NNSTPNLTLR YFPDGARNCY NLNVSRDTNY
MIKATFVYGN YDGHKDEPNF DLYLGPNLWA TVSRSETVEE IIHVTKSDSL
QVCLAKTGDF IPFINILELR PLKKNVYVTE SGSLKLLFRK YFSDSGQTIR
YPDDIYDRVW HASFLENNWA QVSTTLGVNV TDNYDLSQDV MATGATPLND
SETLNITWNV EPPTTKVYSY MHFAELETLR ANDTREFNVM LNGNDLFGPY
SPIPLKTETE TNLKPEECED GACILQLVKT SKSTLPPLLN AIEAFTVIDF
LQVETDEDDA AAIKNVQNAY GLINRSSWQG DPCVPKQYSW DGLKCSYSDS
TPPIINFLDL SASGLTGIIA PAIQNLTHLE ILALSNNNLT GEVPEFLADL
KSIMVIDLRG NNLSGPVPAS LLQKKGLMLH LDDNPHILCT TGSCMHKGEG
EKKSIIVPVV ASIVSLAVII GALILFLVFR KKKASKVEGT LPSYMQASDG
RSPRSSEPAI VTKNKRFTYS QVVIMTNNFQ RILGKGGFGI VYHGFVNGVE
QVAVKILSHS SSQGYKQFKA EVELLLRVHH KNLVGLVGYC DEGENMALIY
EYMANGDLKE HMSGTRNRFI LNWETRLKIV IDSAQGLEYL HNGCKPLMVH
RDVKTTNILL NEHFEAKLAD FGLSRSFPIG GETHVSTVVA GTPGYLDPEY
YKTNRLTEKS DVYSFGIVLL EMITNRPVID QSREKPYISE WVGIMLTKGD
IISIMDPSLN GDYDSGSVWK AVELAMSCLN PSSTRRPTMS QVLIALNECL
VSENSRGGAS RDMDSKSSLE VSLTFDTDVS PMAR

**AtChr1@19256097@19256652 (100%), 20,718.7 Da**
**| potential novel**
**1 exclusive unique peptides, 1 exclusive unique spectra, 19 total spectra, 44/185 amino acids (24% coverage)**

VIISLIVLLG FISVDCGLAP RESPYNEAKT GLTYTSDDGL VNVGKPGRIA
KEFEPLADKP TLTLRYFPEG VRNCYNLNVT SDTNYLIKAT FVYGNYDGLN
VGPNFDLYFG PNLWTTVSSN DTIKEIIHVT KTNSLQVCLI KTGISIPFIN
VLELRPMKKN MYVTQGESLN YLFRVYISNS STRIR

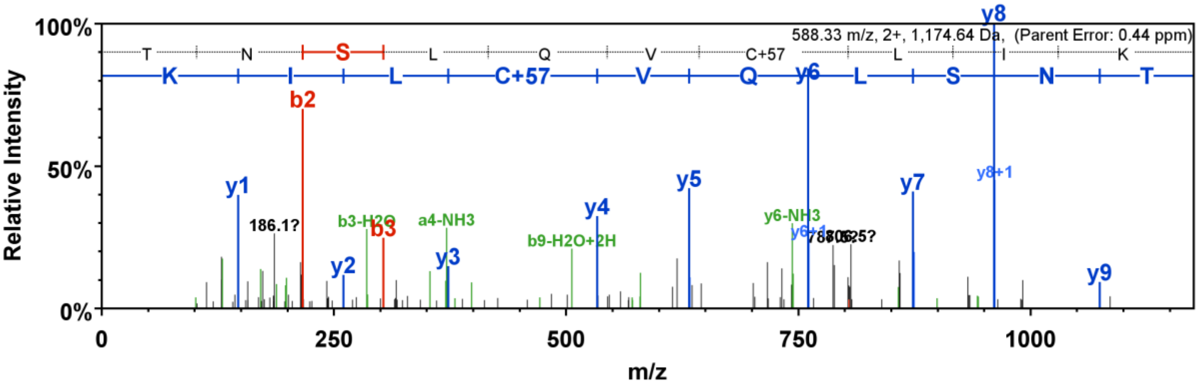


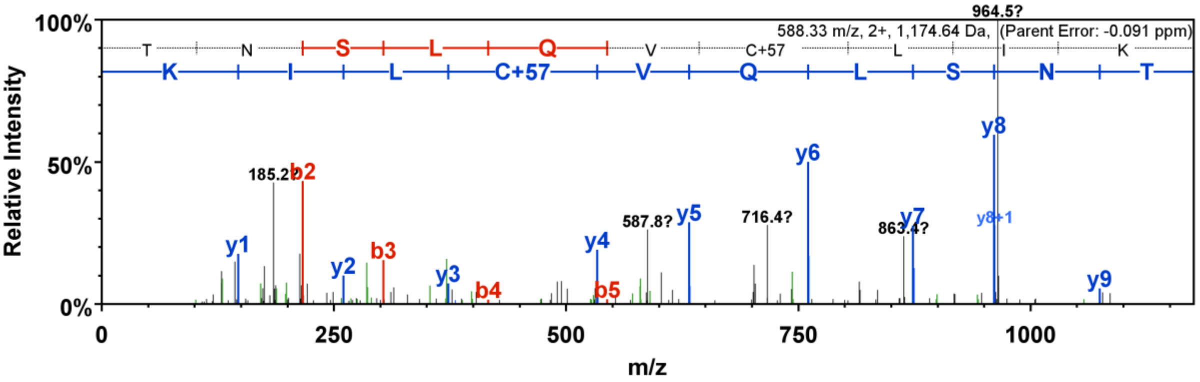

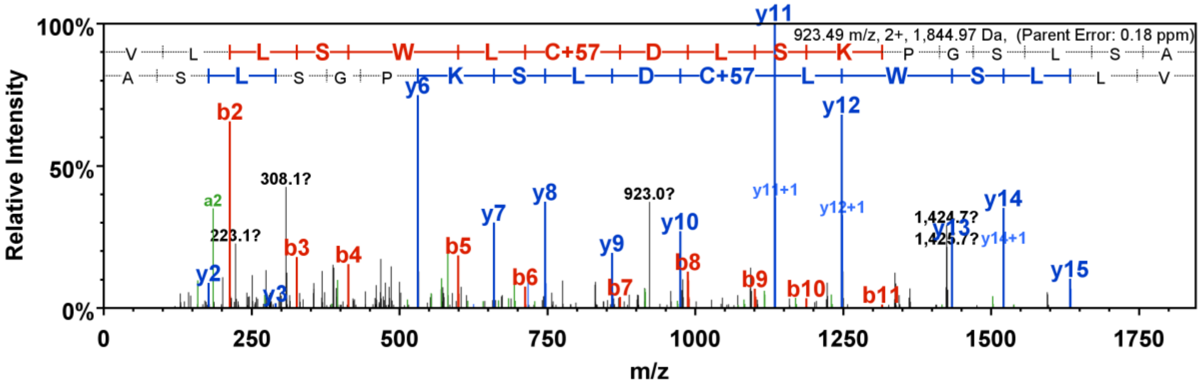
923.49 m/z, 2+, 1,844.97 Da, (Parent Error: 0.18 ppm)
V L L S W L C+57 D L S K P G S L S A
A S L S G P K S L D C+57 L W S L L V
100%
50%
0%
Relative Intensity
m/z
0 250 500 750 1000 1250 1500 1750
b2 b3 b4 b5 b6 b7 b8 b9 b10 b11
y2 y3 y6 y7 y8 y9 y10 y11 y12 y13 y14 y15
y11+1 y12+1 y14+1
a2
308.1? 223.1? 923.0? 1,424.7? 1,425.7?

**No. 2. AtChr5@18590218@18590728: Delay of gemination 1**

Peptide (DCMVDTEGNAGGEEGK) is not in TAIR10 but present in TAIR 11

**AT5G45830.1 (100%), 32,470.4 Da**
**| Symbols: DOG1, GSQ5, ATDOG1 | delay of germination 1 | chr5:18589669-18591400 REVERSE LENGTH=291**
**1 exclusive unique peptides, 2 exclusive unique spectra, 58 total spectra, 112/291 amino acids (38% coverage)**

MGSSSKNIEQ AQDSYLEWMS LQSQRIPELK QLLAQRRSHG DEDNDNKLRK
LTGKIIGDFK NYAAKRADLA HRCSSNYYAP TWNSPLENAL IWMGGCRPSS
FFRLVYALCG SQTEIRVTQF LRNIDGYESS GGGGGASLSD LSAEQLAKIN
VLHVKIIDEE EKMTKKVSSL QEDAADIPIA TVAYEMENVG EPNVVVDQAL
DKQEEAMARL LVEADNLRVD TLAKILGILS PVQGADFLLA GKKLHLSMHE
WGTMRDRRRR DCMVDTEVIF DACTTVNSGP RPTETTNNER N

**AtChr5@18590218@18590728 (100%), 18,446.2 Da**
**| potential novel**
**1 exclusive unique peptides, 4 exclusive unique spectra, 54 total spectra, 54/170 amino acids (32% coverage)**

FRTIDLYQNS NGFGSVGSIW FGLTGGGGGA SLSDLSAEQL AKINVLHVKI
IDEEEKMTKK VSSLQEDAAD IPIATVAYEM ENVGEPNVVV DQALDKQEEA
MARLLVEADN LRVDTLAKIL GILSPVQGAD FLLAGKKLHL SMHEWGTMRD
RRRRDCMVDT EGNAGGEEGK

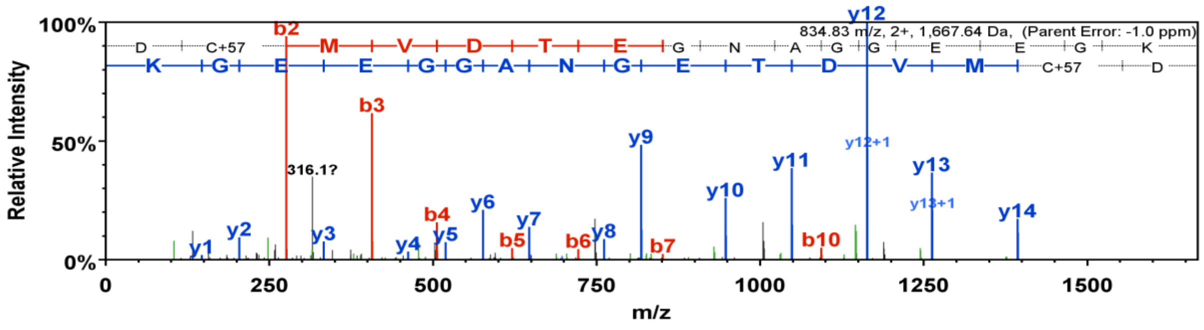


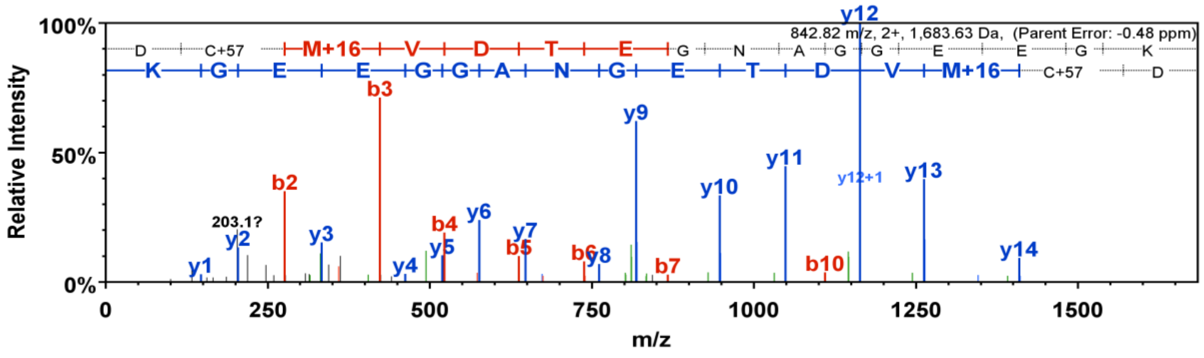


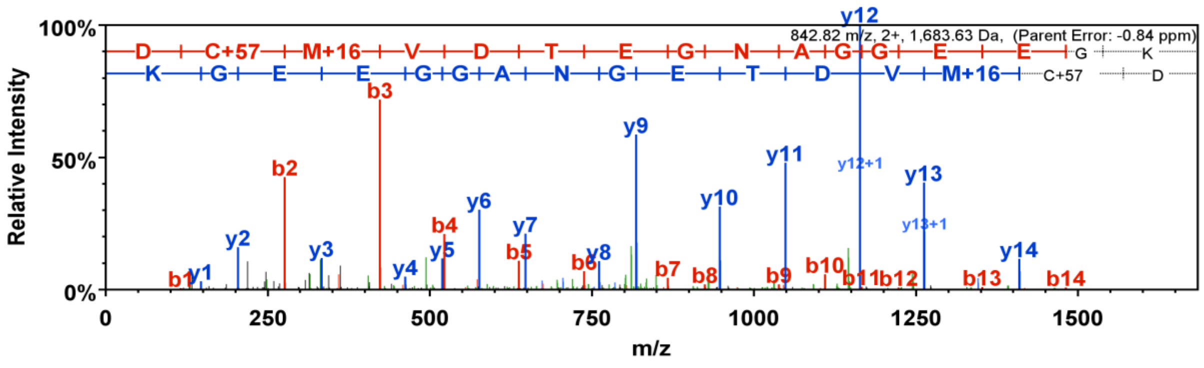

### No. 3, AtChr3@1743539@1745543: MuDR family transposase

**AtChr3@1743539@1745543 (100%), 75,566.1 Da**
**| potential novel**
**5 exclusive unique peptides, 5 exclusive unique spectra, 7 total spectra, 67/668 amino acids (10% coverage)**

```
EFFSSRSSRT TLSEAIPPVP MDDMIDDTMG PEELPISISV SAPPAVTMEE
VMNRAEDAQI IINPSELMSS IVEVPNPKGD ILTKARTQQW QNTITGVGQR
FKNVGEFREA LRKYAIANQF GFRYKKNDSH RVTVKCKAEG CPWRIHASRL
STTQLICIKK MNPTHTCEGA GGINGLQTSR SWVASIIKEK LKVFPNYKPK
DIVSDIKEEY GIQLNYFQAW RGKEIAREQL QGSYKDGYKQ LPLFCEKIME
TNPGSLATFT TKEDSSFHRV FVSFHASVHG FLEACRPLVF LDSMQLKSKY
QGTLLAATSV DGDDEVFPLA FAVVDAETDD NWEWFLLQLR SLLSTPCYIT
FVADRQKNLQ ESIPKVFEKS FHAYCLRYLT DELIKDLKGP FSHEIKRLIV
DDFYSAAYAP RADSFERHVE NIKGLSPEAY DWIVQKSQPD HWANAYFRGA
RYNHMTSHSG EPFFSWASDA NDLPITQMVD VIRGKIMGLI HVRRISANEA
NGNLTPSMEV KLEKESLRAQ TVHVAPSADN NLFQVRGETY ELVNMAECDC
SCKGWQLTGL PCHHAVAVIN YYGRNPYDYC SKYFTVAYYR STYAQSINPV
PLLEGEMCRE SSGGSAVTVT PPPTRRPPGR PPKKKTPAEE VMKRQLQCSR
CKGLGHNKST CKDYLLEC
```

Highlighted: **TQQW QNTITGVGQR**, **LIV DDFYSAAYAP R**, **GLSPEAY DWIVQK**, **E SSGGSAVTVT PPPTR**, **ST CKDYLLEC**

### No. 4, AtChr3@1745505@1745964: MuDR family transposase

**AtChr3@1745505@1745964 (81%), 17,555.2 Da**
**| potential novel**
**1 exclusive unique peptides, 1 exclusive unique spectra, 1 total spectra, 28/153 amino acids (18% coverage)**

```
WFCLFQDMSG KRVIAICMSG GEFQTEKGGS LSYKGGDAHA IDVDEQMKFI
DFISEIGEMF NCDVRTVSLK YFLPDNKKTL ISISNDKDLK RMIKFHENSN
TADVYLLPEE ATPDISNMPA SRLAKPLHKS LYHLIFFKFL RVFFVQIKQN
NVV
```

Highlighted: **FHENSN TADVYLLPEE ATPDISNMPA SR**

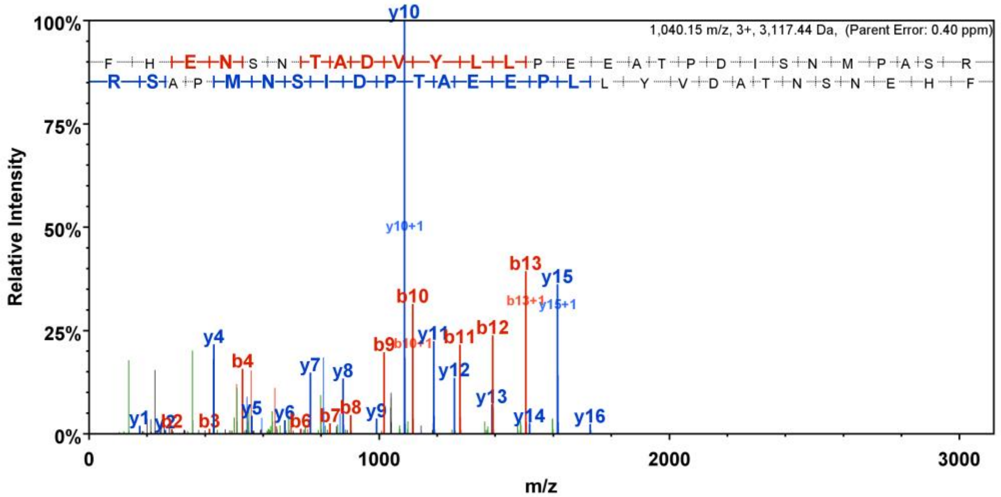

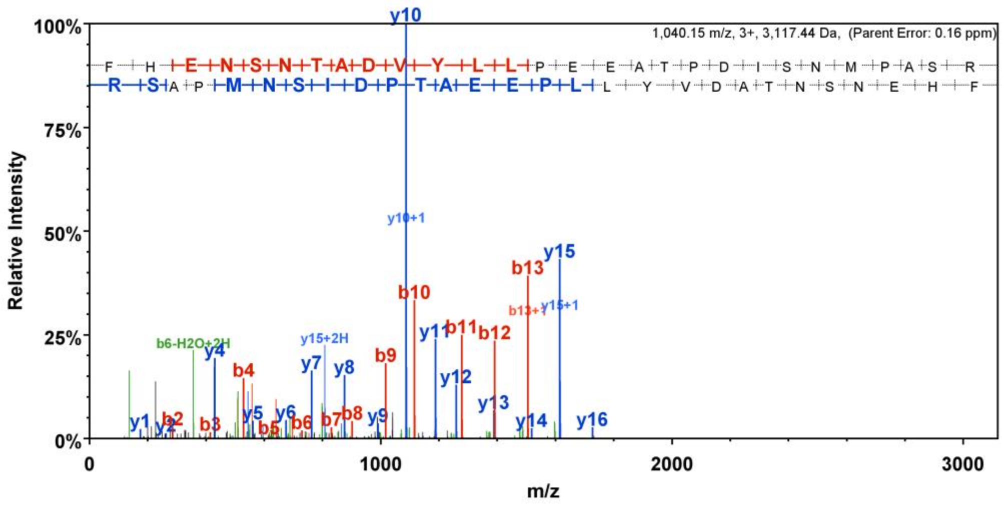
1,040.15 m/z, 3+, 3,117.44 Da, (Parent Error: 0.16 ppm)
F H E N S N T A D V Y L L P E E A T P D I S N M P A S R
R S A P M N S I D P T A E E P L L Y V D A T N S N E H F
Relative Intensity
100%
75%
50%
25%
0%
y10
y10+1
y15
b13
b13+1
y15+1
b10
y11
b11
b12
b6-H2O+2H
y15+2H
y4
b4
y7
y8
b9
y12
y13
y1
y2
b2
b3
y5
b5
y6
b6
b7
b8
y9
y14
y16
0
1000
2000
3000
m/z

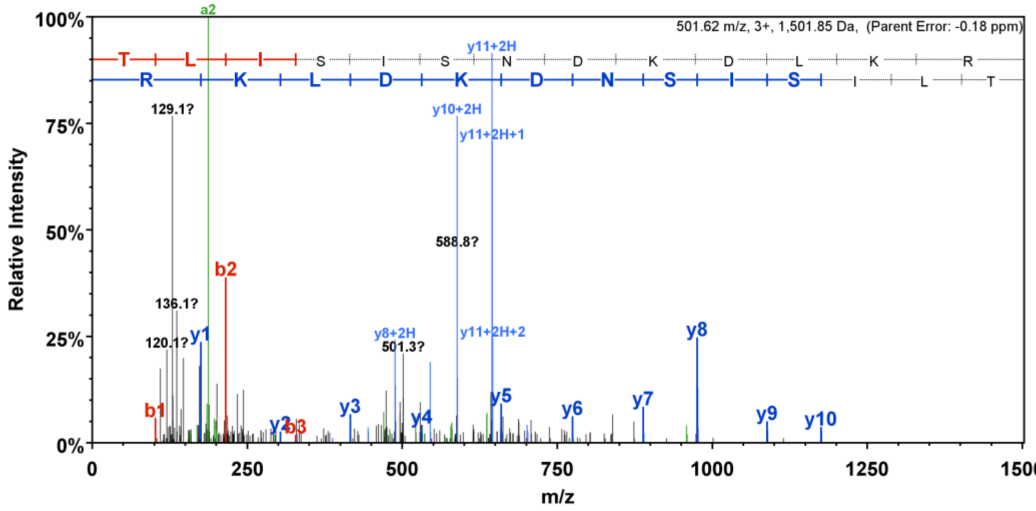
501.62 m/z, 3+, 1,501.85 Da, (Parent Error: -0.18 ppm)
T L I S I S N D K D L K R
R K L D K D N S I S I L T
Relative Intensity
100%
75%
50%
25%
0%
a2
y11+2H
129.1?
y10+2H
y11+2H+1
588.8?
b2
136.1?
120.1?
y1
y8+2H
501.3?
y11+2H+2
y8
b1
y2
b3
y3
y4
y5
y6
y7
y9
y10
0
250
500
750
1000
1250
1500
m/z

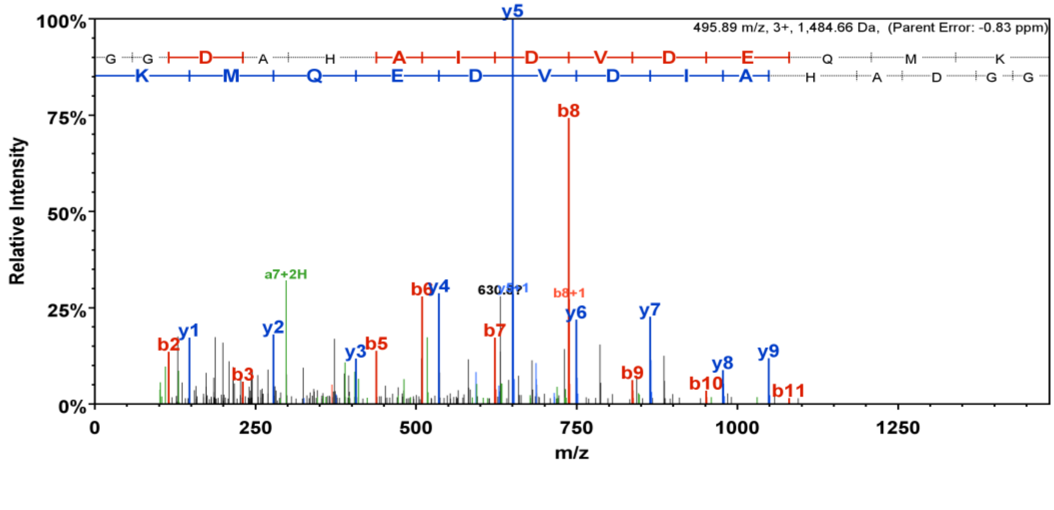
495.89 m/z, 3+, 1,484.66 Da, (Parent Error: -0.83 ppm)
G G D A H A I D V D E Q M K
K M Q E D V D I A H A D G G
Relative Intensity
100%
75%
50%
25%
0%
y5
b8
a7+2H
b6
y4
b8+1
y1
y2
b2
b3
y3
b5
b7
y6
y7
b9
y8
b10
y9
b11
0
250
500
750
1000
1250
m/z

### No.5. AtChr5@20952547@20953303: Transmembrane protein

**AtChr5@20952547@20953303 (100%), 27,669.6 Da**
**| potential novel**
**2 exclusive unique peptides, 2 exclusive unique spectra, 2 total spectra, 33/252 amino acids (13% coverage)**

```
LKKMMIDSLM ELVNKTTSNS LFMFLFCNFI IILILMGNSK PGSQDTPNPG
LQKSVMFSDS VLSSKPGFEN SMMISDSVLQ SKPGFENTVR ISESTISSKP
DFEKSLLTSM PVLLSKPGFE KLGLKSKPVL LSKPGFEKSV LISKSKNLTS
NPGLQEPGLI SKSNLTSNTG LQEPGLISKS SLSSKPAGSE KALTSKPGLE
PGSDIAKNDE KGSLEENESE MECVLRRRVE EFIRKVNTQW KSENTKSNYR
LY
```

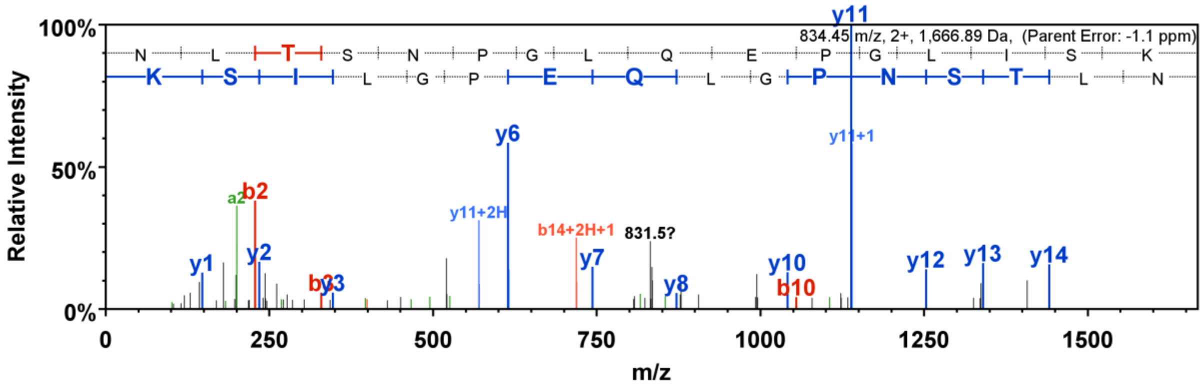


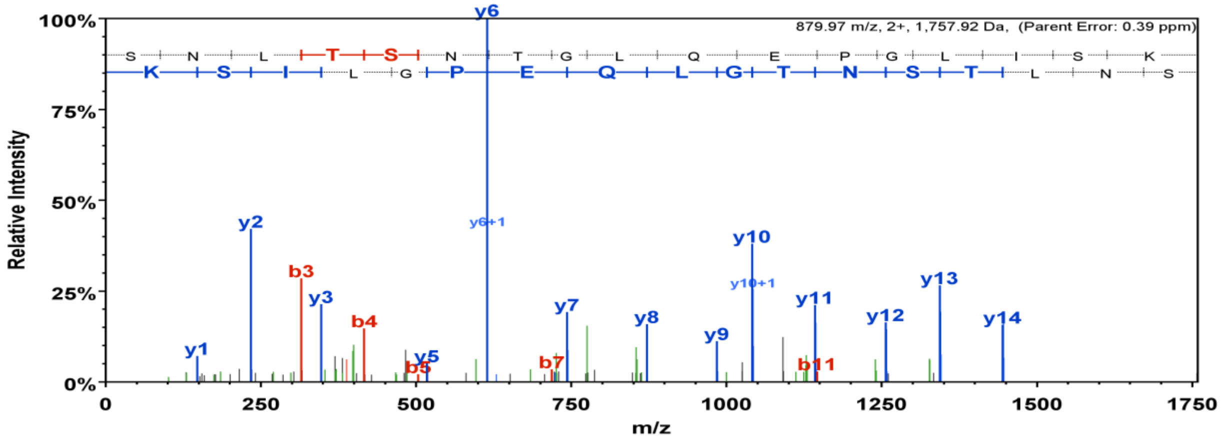

### No. 6. AtChr3@6174451@6174928: MAP kinase 9

**AtChr3@6174451@6174928 (100%), 18,895.3 Da**
**| potential novel**
**1 exclusive unique peptides, 1 exclusive unique spectra, 1 total spectra, 22/159 amino acids (14% coverage)**

```
KWWWRCLMGA  SHSTNVNNHP  HSRNASNHPL  TNSNSTSSRH  SASSSDRLSV
SNLRSQLTTI  YRNQEEEEEE  EEEEEEEEEG  GKEKRAEEEA  KSFSLVRDFD
LSGLNCIRVS  RRNYILMDPH  KKVRFCSLCK  RFGYCLLIWI  CRFIYLFDFS
RFRIDFWTS
```

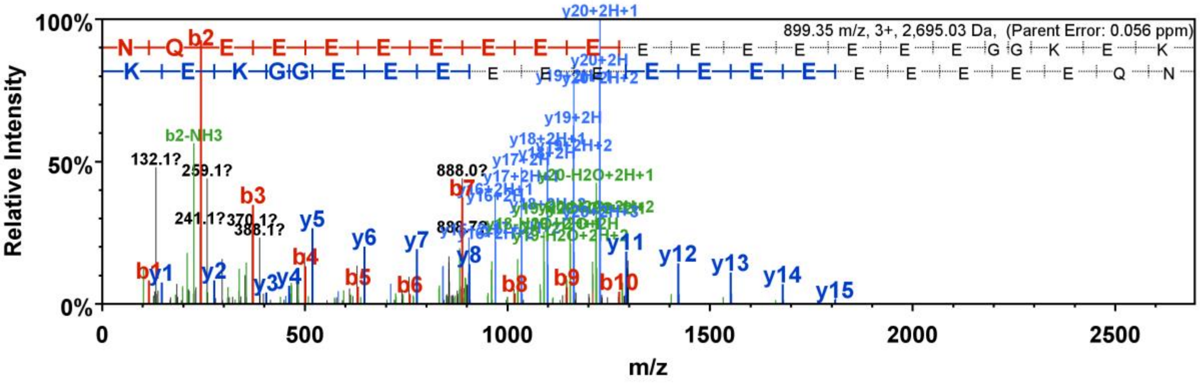


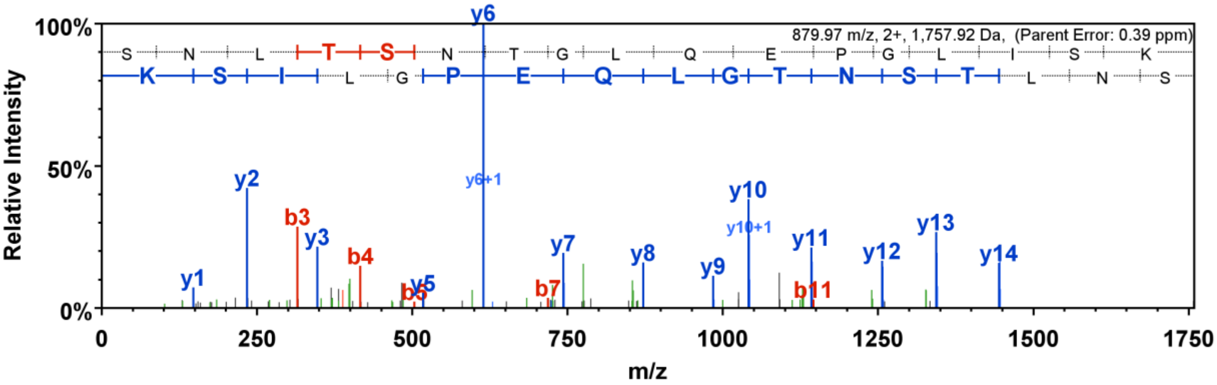

### No. 7. AtChr4@9187917@9188217: GDSL-like lipase/acylhydrolase superfamily protein

**AtChr4@9187917@9188217 (100%), 11,802.8 Da**
**| potential novel**
**2 exclusive unique peptides, 2 exclusive unique spectra, 4 total spectra, 22/100 amino acids (22% coverage)**

VLFLKFYTRV CVEFELHFHK FLYNRLISDP IGCIPFERES DPMAGYECSV
EPNEVAQMYN LKLKILVEEL NNNLQGSRFV YGDVFRIVYD IIQNYSSYGT

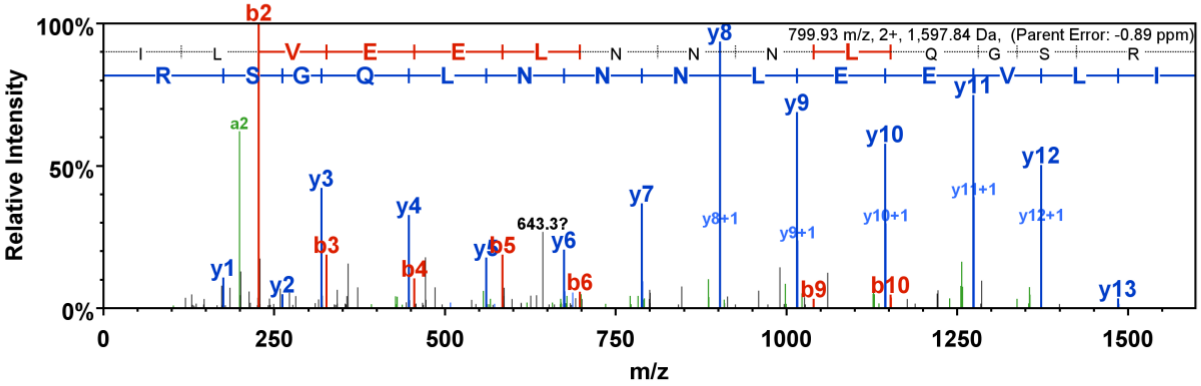


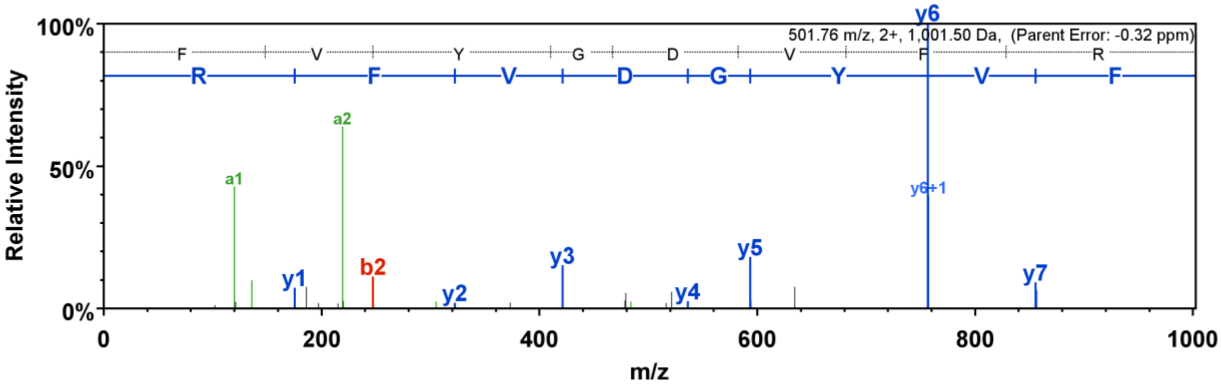

### No. 8. AtChr3@3252605@3253004: Plant self-incompatibility protein S1 family protein

**AtChr3@3252605@3253004 (100%), 15,184.1 Da**
**| potential novel**
**2 exclusive unique peptides, 2 exclusive unique spectra, 3 total spectra, 23/133 amino acids (17% coverage)**

```
KKMNRLIAFL  LIIALSFGLN  KACEDCTIVF  RNNLSPGIIL  KVNCESNNKN
RVTGTVKFQS  DTVRINFREA  AFERTTWHCL  VQQGGYSQHF  RAYRGSAPIP
RCGELRVYIA  KRDGIYLSAN  AGPEKLDQRW  MKN
```

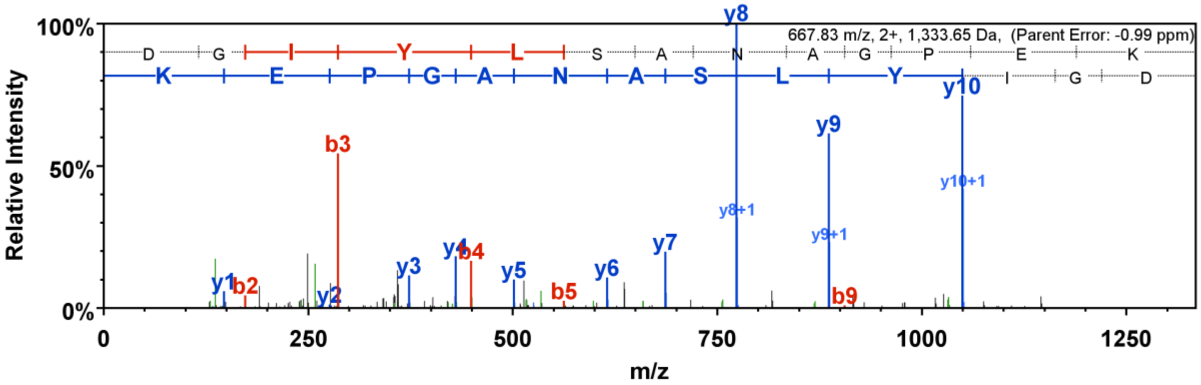


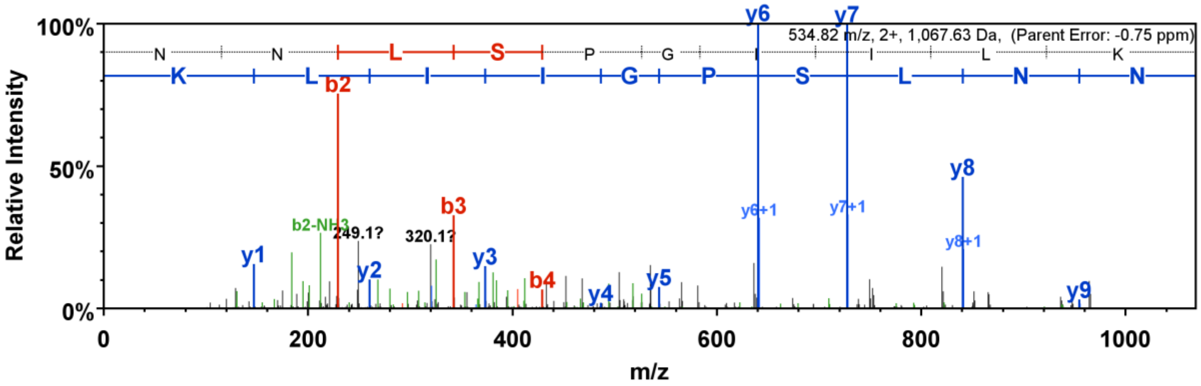

### No. 9. AtChr5@4376549@4378589: Unknown protein

**AT5G13590.1 (100%), 129,532.6 Da**
**| Symbols: | unknown protein; Has 150 Blast hits to 121 proteins in 42 species: Archae - 0; Bacteria - 8; Metazoa - 80; Fungi**
**2 exclusive unique peptides, 2 exclusive unique spectra, 17 total spectra, 115/1168 amino acids (10% coverage)**

MSGSQEPRIR PSTWSCSDIP IKKRKYLVQP QMEEAVSTQI PQPNEQGDTR
SAHADETQKM TGR**EPTSSLP** **SVPVGISGK**G K**SIGNIVFDQ** **TR**VKFEKPSS
PIHSSPLAGF DIPSSSNVLG SSIHFPMGKL PVGAEHAGLV VPSNQTRMKV
EKTVLKTHDI VRKTGDKETL RGECQTEASS GAKTVSLQLS CNTKNNSPYW
KDEEPTELNL SLSKGVCPAH NTDSTSTKSG NSGLNRENWD LNTTMDVWED
ALDRTRGAFL NSNRSLRDIE RSSCRDTTAI TKSVSERQKE SVGFSSPKVT
LMQFDNHVNP TCSLSLGLSS YPPIEK**SPSL** **PATTSEAR**AG NVCSVNLRTV
KSEIVEESVR **QATESTQVSP** **IGLSIK**GLKH EGIGRFSQGN SPSFGILK**TV**
**VPISIK**AEPN TFSQSEVFNR KDGMLNHPHT PIMQSNEIPD LPTSSTPYQK
DKYLPCSNGI SNAPMPLSGM TIIPGVQSDP DCTSKENSGQ SSSLANGKLR
EVLKHGGVYT TYSGHGDHNL NASGVNVTSL TEEKILDDCK PCISKELPCN
SRGTDELSRN DEEKITLPGK ELEEQLYSYG FESDRGYDLS RVIKEQVGKR
NLCDDGK**VQG** **PAAVFTESNE** **VAHPECGGSE** **TEQR**NINVPC HVHFHNSNHV
EEK**GSQPALL** **GYTGETEGR**I VQDGEGTSGV STVSGGIENP EIVDNSSPVS
LKAEMSTIDN DSPMECSDGS QSRIINLTQV KSPVKALDAS GSFVPPRMER
DRFHDFPLEP REYTFRGSDE SCKFSRERYH GRIMRSPRLN FIPDRRRLPD
NTESNLHDQD TKKFEFDNHG NTRRGGAFMS NFQRGRRPAN DGVTPYAHSF
PRRSPSFSYN RGPTNKEDTS AFHGFRDGEK FTRGLQCNNT EPLFMNHQRP
YRGRSGFARG RTKFVNNPKR DFPGFRSRSP VRSRERSDGS SSSFRNRSQE
EFSGHTDFSH RRSPSGYKVE RMSSPDHSGY SREMVVRRHN SPPFSHRPSN
AGRGRGYARG RGYVRGRGYG RDGNSFRKPS DHVVHRNHGN MNNLDPRERV
DYSDDFFEGQ IHSER**FGVDV** **NAER**RRFGYR HDGTSSSFRP SFNNDGCAPT
NVENDPDAVR FQQDPRIKIE EQGSLMEIDG ENKNSTENAS GRTKNMEEEE
TSKNSKIWQP DELGGDGF

**AtChr5@4376549@4378589 (100%), 73,948.7 Da**
**| potential novel**
**2 exclusive unique peptides, 2 exclusive unique spectra, 6 total spectra, 55/680 amino acids (8% coverage)**

DSHIDVFFLC LIQPRIRPST WSCSDIPIKK RKYLVQPQME EAVSTQIPQP
NEQGDTRSAH ADETQKMTGR EPTSSLPSVP VGISGKGK**SI** **GNIVFDQTR**V
KFEKPSSPIH SSPLAGFDIP SSSNVLGSSI HFPMGKLPVG AEHAGLVVPS
NQTRMKVEKT VLKTHDIVRK TGDKETLRGE CQTEASSGAK TVSLQLSCNT
KNNSPYWK**NE** **EPTELNLSLS** **K**GVCPAHNTD STSTKSGNSG LNRENWDLNT
TMDVWEDALD R**TSGAFLNSN** **R**SLRDIERSS CRDTTAITKS VSERQKESVG
FSSPKVTLMQ FDNHVNPTCS LSLGLSSYPP IEKSPSLPAT TSEARAGNVC
SVNLRTVKSE IIEESVRQAT ESTQVSPIGL SIKGLKHEGI GR**FSQGNSPS**
**FGILKTVVPI** **SIK**AEPNTFS QSEVFNRKDG MLNHPHTPIM QSNEIPDLPT
SSTPYQKDKY LPCSNGISNA PMPLSGMTII PGVQSDPDCT SKENSGQSSS
LANGKLREVL KHGGVYTTYS GHGDHNLNAS GVNVTSLTEE KILDDCKPCI
SKELPCNSRG TDELSRNDEE KITLPGKELE EQLYSYGFES DRGYDLSRVI
KEQVGKRNLC DDGKVQGPAA VFTESNEVAH PECGGSETEQ RNINVPCHVH
FHNSNHVEEK GSQPALLGYT GNKHIKACST

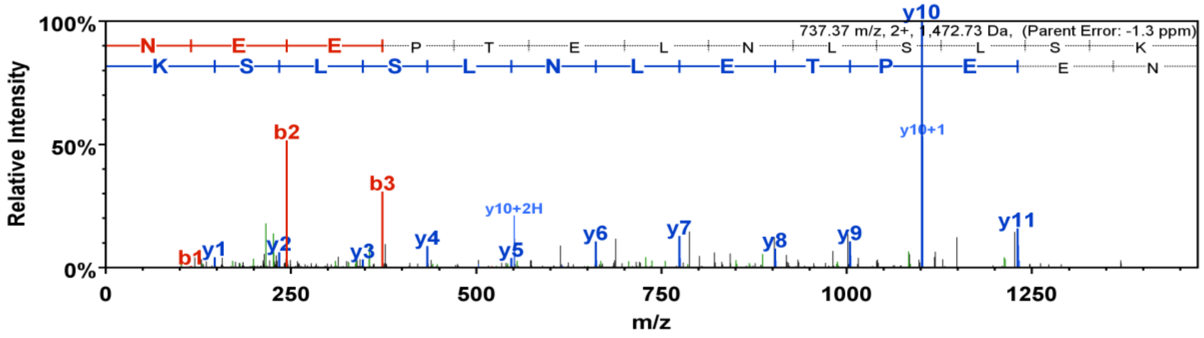


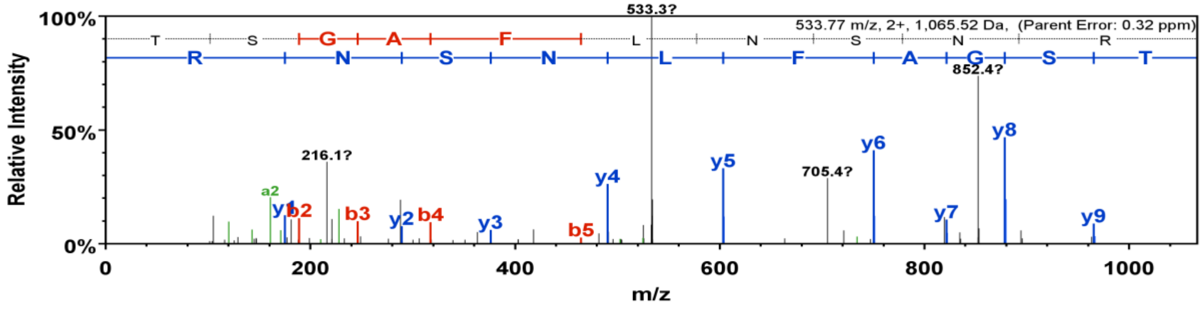

**No. 10. AtChr4@15093731@15095657: ARM repeat superfamily protein**

Two peptides (LDNTDEELLSKLDPFVK, and YEDIVSSSLR) are not present in TAIR10 but in TAIR11

**AT4G30990.2 (100%), 296,991.4 Da**
**| Symbols: | ARM repeat superfamily protein | chr4:15084456-15097860 FORWARD LENGTH=2620**
**11 exclusive unique peptides, 11 exclusive unique spectra, 25 total spectra, 167/2620 amino acids (6% coverage)**

| | | | | |
|---|---|---|---|---|
| MATSADARAV | KSLNTSEGRK | RFVFKSASQR | **TNDIDNISNY** | **R**NLDKVKAEP |
| SEGSTFFRDC | LIEWRELNTA | EDFILFYEEM | LPSVQSLSLI | IMQKER**IFSN** |
| **LVSR**LQMKAR | LSLEPILRLI | AALSRDLLND | FIPFLPQIVN | SFVTLLNNGA |
| HNDPEIIQQV | FTSWASIIVS | LQKYLVCDIE | GILRDTLELR | YYPKDNISEF |
| MSESMSFLLR | KAQDEQLEKG | MKMILSEVAH | PSKK**AGGVGV** | **LYNVMR**GTYG |
| RLHSKAGRVL | SFLLKDSTLS | FLDNFPQGPC | TVVEVVSLVL | QRICEDLEAE |
| KLSAMWEYLY | KKINKSISNK | KSVHLSR**LLS** | **VLMAVVK**IKE | GRKVHDIPSL |
| IGIVSRIVST | FFTSSETAVE | GDNLSAVLDE | VLELILCTIN | TVNEMETVAS |
| LWAPIFALK**S** | **SSLLTFLR**EF | LKKDQSVVKA | FTKNILCAIN | NMIWESSEEV |
| IPLLLTLCEE | HKTQQTSHDV | VNSISQTFES | RYERIHEFLE | AKIKKVQQNI |
| ENAGLAQINE | AELAAIWGVV | KCYPYFKVDS | SLLICFKKTL | RQHLAVSDVD |
| TCSGPELMWQ | SLLGTTLRSC | YKMTGINHSD | LEEALSFAKD | YKSCEQVLSP |
| VADVLEFMHR | PALAHGRSKP | YPELQANK**AG** | **DAFEIFSENL** | **R**HPNKNIRLM |
| TLRILCHFET | LSSDPSFEEH | PPKKKMKTEK | NVLQLLLLFE | ETAPTVDTSR |
| MLAGYISTIQ | DNLSAGRIHS | AYVKLVLNGM | LGILHISYRP | LCVQASECLA |
| VLVRKYTGAV | WSDFVCYLGQ | CQLKFETLHD | HSENANQSMS | ERHACNLNLN |
| GRFNLFLFPP | SAITPTATVS | DVVSQLLQTL | QKASSVAQSR | **ASEILPLLLK** |
| FLGYNSENPG | SVGSYNGRVC | KGEDWK**TVLV** | **QWLTLLK**LMK | NPRSFCFSQF |
| LNDILQNRFL | DDNDAEIQTN | VLECLLLAND | FLLPHRQHLL | NLIKPKELRE |
| ELTTWNLSEN | IGEPHRSYIF | SLVIRILMPK | VRTLKNSASR | KHTSIRHRKA |
| VLCFISQLDV | NELALFFALL | IKPLNIISEE | TMDSFWSSGK | SSLDYFQNSN |
| FLKYFTVDTI | STLSRNQKFG | FLHVIQHILE | VFDELRVRPF | LDFMMGCVVR |
| **LLVNYAPNVD** | **EEMNIDSLAL** | **R**NVTAAPSTS | DDKENASINH | DQAGTAFKQF |
| KELRSLCLKI | IAHVLDKYED | CDLGSEFWDL | FFSAVSPLIK | SFKQEGSSSE |
| KPSSLFSCFL | SMSKSRNLVN | LLCREESLVP | DIFSILTVTT | ASEAIKSSAL |
| KFIENLLCLD | NVLGEDENMI | RGFVDPYIEA | LINSLHSLFI | GDILKRKSVK |
| YHGEREIKIL | KLLSKRMQDR | SHVMKYLDVL | LSFLNKSVKD | PDIRREALLA |
| IQDIIAYLGM | ESTSKIINTV | SPLLVDAELD | VR**LCICDLLE** | **SLAK**IDFSLD |
| DVAKRVRDMN | AISAMEVDDL | DYEKIVNAYV | EINADFFIKS | SEQHTMIILS |
| QSSILCREAP | AHSEFGKEVK | NADVSWTGDR | VLCILRNFIL | KHIGDAINRG |
| GIIIKEWILL | IREMVTKLPD | AANLSAFRPL | CSEDENVDFF | KAIVHIQAHR |
| RARAISRFSS | VVKDSSLPEG | VVRKLLVSVF | FNMLLEGQDG | KDNNVRNACT |
| EALASISAHM | SWTSYYALLN | RCFREMNKHT | KKGKILLRLI | CLILDKFHFA |
| KDGYPHEAEE | IRTCLQKIVF | PRMQKLMNSD | SDNVNVNSSV | AALKVLK**LLP** |
| **EDVLDSNLSS** | **IVHK**IASFLK | NRLESTRDEA | RLALVACLKE | LGLEYLQVVV |
| NILRAILKRG | SEVHVLGYTL | NSILSKCLSN | PTCGKLDHCL | VDLLAVVETD |
| ILGEVAEQKE | VEKFASKMKE | TRKRKSFETL | K**LIAENVTFR** | SHGLK**LLSPV** |
| **TAQLQR**HLTP | KIKTNLEKML | KQIAAGIEGN | TSVDQGDLFL | FIYGLVDDGI |
| NNRSGLGDQV | SLPPSKKKKK | SRDLKETSGL | CFGPKSCPHL | ITVFALDLFY |
| NRMKKLRLDN | TDEELLSKCF | TSLVKFPLPS | LTSEADELKT | ALLTIAQSAV |
| SSSSPLVQSC | LKLLTTLLKN | INITLSSEQL | KMLIQFPIFI | DLESDSSFVT |
| LSLLKAIMNR | KLVVPEIYDI | AIQVSKLMVN | SQLESIRKKC | KHILLQFMVH |
| YTLSEKRLEQ | HVNFLLENLR | YEFPTGR**EAV** | **LDMLHALILK** | FSEPNLGKQS |
| VLDQQSQKLF | IQLTVCLSNE | TDRKVLPLVG | AVIEVLIGRM | SKDQVDSSLL |
| YCLCWYKQQN | LSAAAAQVLG | FFISAMKKTF | RKHIYNTVED | ARTILESAIS |
| ASSLQLQDTV | EEASLPFWKE | AYYSLVMIEK | MLEQFPDLRF | GKDLERSPAR |
| KITALALSAS | ITTLDLDIWK | MVFKFLLHPH | AWLRNKSCRL | LNLYFEALAG |
| RKRPECRTLV | ADSLLEKPSS | LFMVAVSLCF | QLKEQPTTGN | IDVDLLTANI |

**AtChr4@15093731@15095657 (100%), 72,626.0 Da**
**| potential novel**
**2 exclusive unique peptides, 2 exclusive unique spectra, 21 total spectra, 89/642 amino acids (14% coverage)**

GDPLNFKYFS MQLLYSDRLI QMLVLQGVVR KLLVSVFFNM LLEGQDGKDN
NVRNACTEAL ASISAHMSWT SYYALLNRCF REMNKHTKKG KILLRLICLI
LDKFHFAKDG YPHEAEEIRT CLQKIVFPRM QK**LMNSDSDN** **VNVNSSVAAL**
**K**VLK**LLPEDV** **LDSNLSSIVH** **K**IASFLKNRL ESTRDEARLA LVACLKELGL
EYLQVVVNIL RAILKRGSEV HVLGYTLNSI LSKCLSNPTC GKLDHCLVDL
LAVVETDILG EVAEQKEVEK FASKMKETRK RKSFETLKLI AENVTFRSHG
LK**LLSPVTAQ** **LQR**HLTPKIK TNLEKMLKQI AAGIEGNTSV DQGDLFLFIY
GLVDDGINNR SGLGDQVSLP PSKKKKKSRD LKETSGLCFG PKSCPHLITV
FALDLFYNRM KKLR**LDNTDE** **ELLSKLDPFV** **K**LLAGCLSSK **YEDIVSSSLR**
CFTSLVKFPL PSLTSEADEL KTALLTIAQS AVSSSSPLVQ SCLKLLTTLL
KNINITLSSE QLKMLIQFPI FIDLESDSSF VTLSLLKAIM NRK**LVVPEIY**
**DIAIQVSK**LM VNSQLESIRK KCKHILLQFM VHYTLSEKRL EQHVNFLLEN
LRYVLCTSYE FNFFSFSYSK LICSSGMSFQ LEEKQFLTCF MP

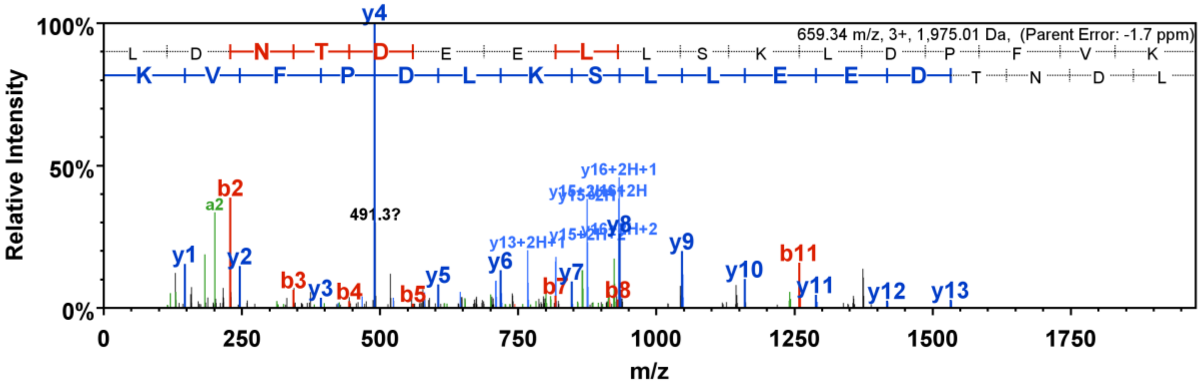


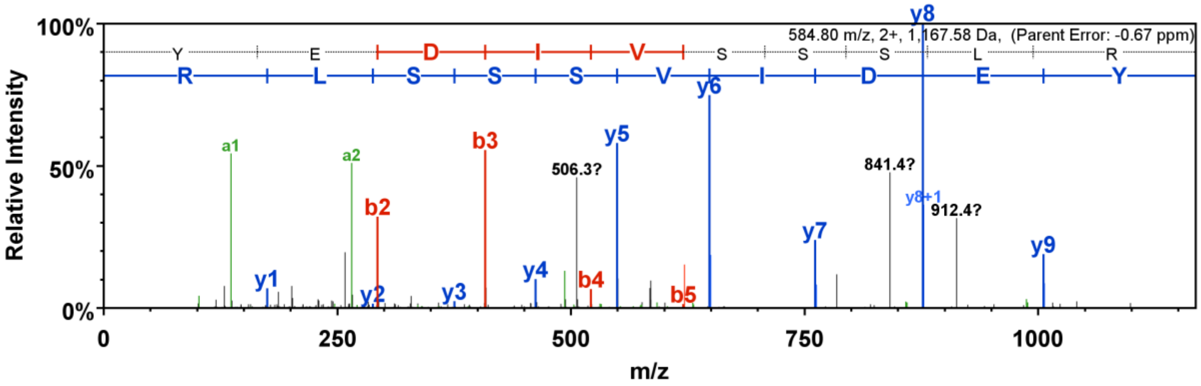

### No.11. AtChr4@13558438@13559653: Ribosome 60S biogenesis amino-terminal protein

Peptide (QQSAALSLLASIVR) is not present in TAIR10 but in TAIR11

**AT4G27010.2 (100%), 269,816.0 Da**
**| Symbols: | INVOLVED IN: biological_process unknown; LOCATED IN: cellular_component unknown; EXPRESSED IN: 20 plant structures; EXPRESSED DURING: 10**
**15 exclusive unique peptides, 17 exclusive unique spectra, 54 total spectra, 282/2402 amino acids (12% coverage)**

```
MASEIAKKFD FKGFAKLAEY NTQGTEKVKK HSTRKAFVGF AISFLEVGKP GLLSSVLNKK
EMYSKVLPGL GKDDDDTVAS VLSTLKDKIL VEESLISPGL RSVLFGIVTL KHLASISARE
DAGIVNELAH DVLVKVCTDP SNGLMPDAKR KLRGNSDRLL MLMKGLRAAE IGYHRDLLLA
IVRGRPSLAS DFLDEFPYNV EDFSSPSWFS SISLAANLVS SVRTSCSFDF LNPDQRATPP
SGGSDVQTIM KCICPRPFSR SLITKGMLHS DFLVKHGTLR FLLETLRLLD SFLTAWNLCS
SHRCSVEQIQ ISLERNVMGE VSSFFPDSQV LLIVLKSLDG SSGTQKLSLK REAELDSGLV
GRKKRIKRSE KDVLEEEAVD IVIGGVGSDK DIFLAEDNMD AHMTDQEDAE KEYLGIVSDI
WISELCSNPI DSVEEAEMCF HIKLLDALKI YVRAVPNELE GSFDIFMKFL SNSFGMPVEL
QRALLSLLSE YISWTPKSQS DRGPTRIPPL MHKHLRVFIN LLLFSPHNGV KDLAYNLAVA
AMNSTGAFEN NPSEIGAWFL FLPCFEKIKL PLELQEAVQS MSSVVVSFLC DAVSTVGNNL
FKHWDIVRSS LSHLKGVSIG FSPLIICLLQ KCVRLLNSES KTSLPEKSAI SLYVCSTLKY
LLQTQVDSKL LSCLIQSVLS EVVDESKDSL CEWRPLRMLL CFSQSLSNEK PIILHSRRTT
GLPADSSFAE TLDEIKRLVR SISPDEIAGI VKAFSSALIC ATPESILQNF ASVMDVSWAF
YGTPFSFLQS ITFLEENFLG NLSKLSPDLF ASGSEFTGSG NLCEGTVDSE IDFSGHSSVT
EEIRSKMDNR DMESSAFSIF LKQAPFPVLL NAIMSMDISC LPEFPRISEL LLLKVSQPKS
GSIDSNIQLI LFWLFQIRSS YKVQPAPVLH QLSEICLRLM KNLFSQISEP ELVSGPSSNK
LPASFAKWKH QVAETVLCHP VVMALLESPL DCGTLPPVQN VEIFSETSLT MGRLVFSEID
QHILNLLVST CEHFLFDEKP PNLWKEDLRK NKSIIAFKDL VERLLLEFRV KFELCVGSQS
YVSLLQPAQL IHALLRFISP FKLFNIAHSM LSKIDEEGLT SPNSSIILSL GLGIAGGAFE
MLVLYSHQPT AKRGVYDLLW ELEEKNYASN IIEKVYSMAC KFSTSLDLDS ADICLLKVCG
GIFRGKHNQN YAVDPLVLKI SLIVGRTPED LIIHCINRAS ITRAKILFYL VESSPLHLLV
FGHFFFSMLS KKQDDSALTD DQFIMLLPAV LSYLTSVIAK LEKPCNRCLD ITSVYSNILI
NGFLQWPRFL ARCIFEEKHE EILLSTTEDM ETMFNASLIG KAVRMFQYHF SLTESPTKED
DLFKVFNSMF PLSSTGKEML DYEIKEVDVQ SVDQILNVAI RVVAKVTVSR ICLFPEDSSM
CHLKRAAGTC VKESSSKIGC NRAILSKPLL DALVNSWQCV VKKSDGSFKG NYEGKQDRCW
SLCKSLENFI LRSILQFLES MCEELVQLDS LPFLDRLMKS VLLYRFEDSK TLKILREIFS
LLSRGKYSYA PYIQRLIYHS RFTPTISSLS ISSSNTGELF RPVSSILNHL IILSPDSVRV
KRCCLEAPKY AKQLEIVKIL RVLLSNCGKD SGMKELLSDL HFLLLCSYGA TLREIDLEIY
KLMHDIKLIE AEQTLNVSET DYLWGKAALK IREGLSQDAS DVCQVDLVED VRQGLIKENL
CVDPKICALT VLFFPYQRTT EKSENFYLYD DPINEVPVFS FNFQLIVLGY IEPVEFASLG
LLAVAFVSMS SADLGMRKLG YETLQIFLDA LEMGKDIVEG NILAFPICTE DFNWFCKGLM
NCRKNKHVTG LRLLLMYVQN GVEEPWQRIP TVSAIFAAET SMILLDPSHE HYVPINKLLK
SSSTLKLRGI PLFHDFFWSS AVNFRSQRFW ELRLVYLGLK SDDDVQIYIK NSILETVISF
SSSPLADDET KRLILQVVRK SVKFHKIARH LVENCGLFSW CSSFISNFTT KPIGDKDLHL
VVVLEIITDV LASRNITEWL QRFGLEGLME ISSRLYKLLG GGLVSVQENG TSVDLILQIL
SATLKISQKR NMYQPHFTIT IEGIFQLFEG VANFGSPQVE ASAESGLITI LMSTPPVDIL
CMDVDKLRRF LLWGTSTALK SDFKKGSKPS ESHEDTKILI EGPQEETMVA KFLRWLSASV
ILGKSYSKAS DSDPTFLSKT KPETLLTSLE YFKKRNLEDS MQNSEHIIGE VIVHLQQFLS
TNYMFLLPSV VFALSLMLLH NDLGTGESDG DYKLIKSLCS KISSPPEAIP GWRWSYYQAW
RDLSSEQATD LDKINELHAC QHLLLIFSAM LGETPQESQQ VLLRKSFDMS HVFEWERSLV
ET
```

**AtChr4@13558438@13559653 (100%), 45,115.6 Da**
**| potential novel**
**1 exclusive unique peptides, 2 exclusive unique spectra, 15 total spectra, 93/405 amino acids (23% coverage)**

```
LNTLFDISGM VVDGGNRDVE VGKIPVMAFR PSHEAKLREL LHNICLHEIK
LCSDAAKEFV KLLKGETGGD LLRLYFQSSP NFAELLEAWK LRHEKQGLSY
IFSLIQTILS HPEGKDRSTD IGRAIDQFGR LLVEEKLDDI YKELNSKEGK
QQSAALSLLA SIVRRGPGMA SEIAKKFDFK GFAKLAEYNT QGTEKVKKHS
TRKAFVGFAI SFLEVGKPGL LSSVLNKKEM YSKVLPGLGK DDDDTVASVL
STLKDKILVE ESLISPGLRS VLFGIVTLKH LASISAREDA GIVNELAHDV
LVKVCTDPSN GLMPDAKRKL RGNSDRLLML MKGLRAAEIG YHRDLLLAIV
RGRPSLASDF LDEFPYNVED FSSPSWLVFF RVRKCIYLYF TASTVQFLLF
CHSSN
```

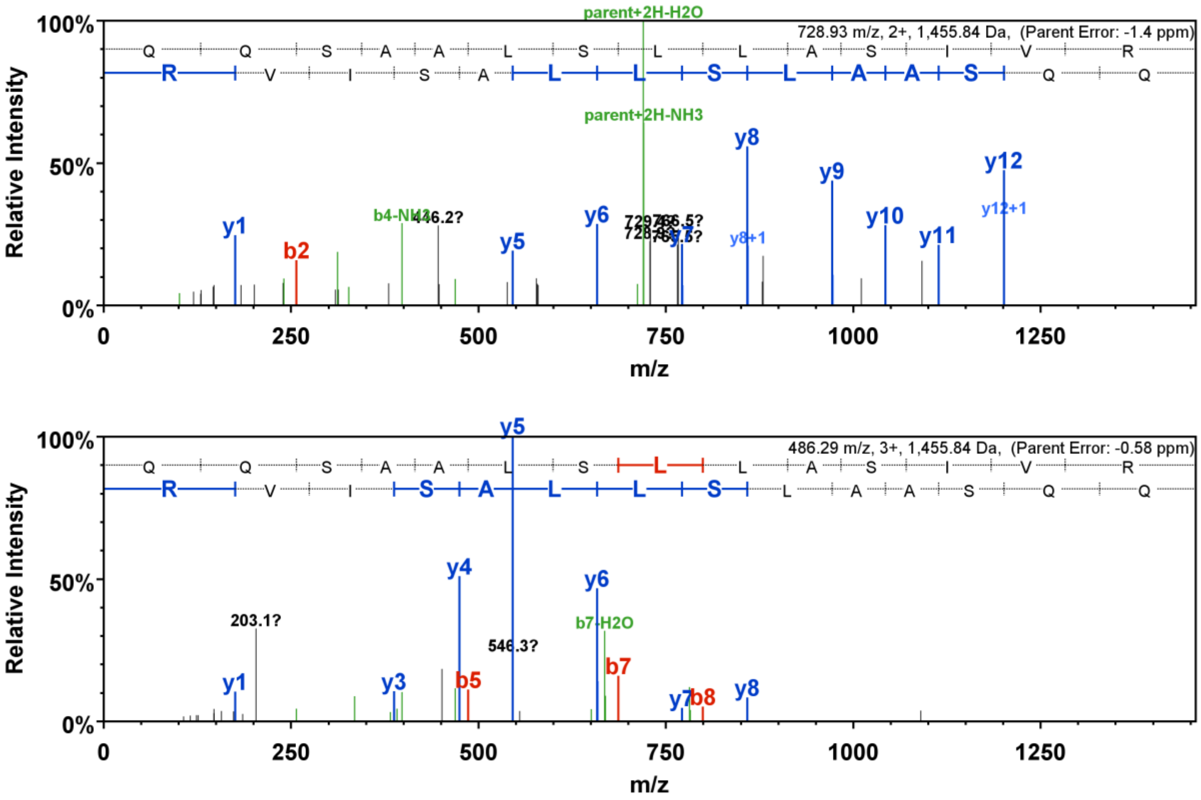

728.93 m/z, 2+, 1,455.84 Da, (Parent Error: -1.4 ppm)
parent+2H-H2O
parent+2H-NH3
Relative Intensity
m/z
y1
b2
b4-NH3
446.2?
y5
y6
y8
y8+1
y9
y10
y11
y12
y12+1
486.29 m/z, 3+, 1,455.84 Da, (Parent Error: -0.58 ppm)
203.1?
546.3?
y3
y4
b5
b7-H2O
b7
y7
b8

### 12 AtChr4@7455224@7455494: Transcription initiation factor IIF subunit alpha RAP74

**AtChr4@7455224@7455494 (100%), 10,497.8 Da**
**| potential novel**
**3 exclusive unique peptides, 3 exclusive unique spectra, 4 total spectra, 23/90 amino acids (26% coverage)**

E Y N V R **A A A P T** **D K N Y F I G R F V** **T G L P N F K K** G S E N K W S L R K D I P Q G R Q F T D A Q
R V I L C L F W I I P I I L M V F T M S F V W I L I F V G E V K E Q T L D L G R

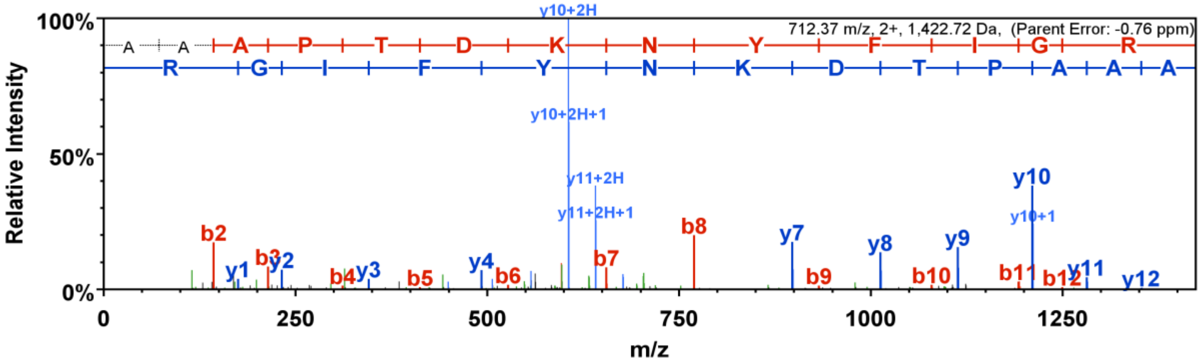


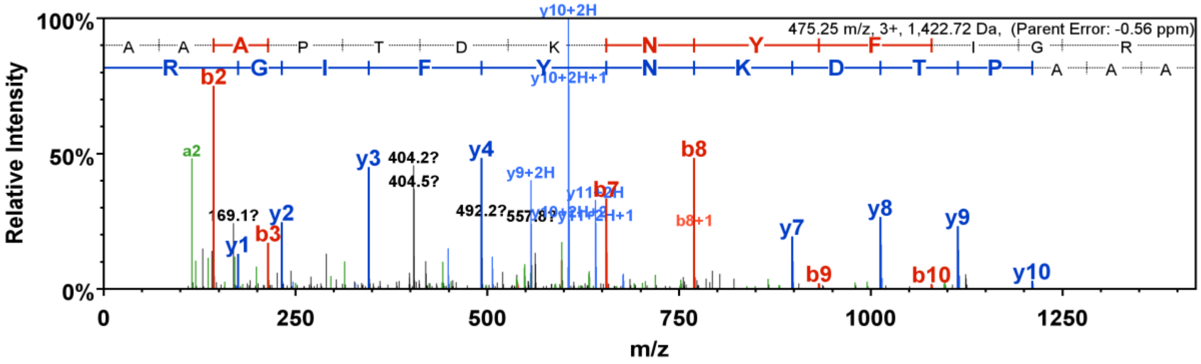


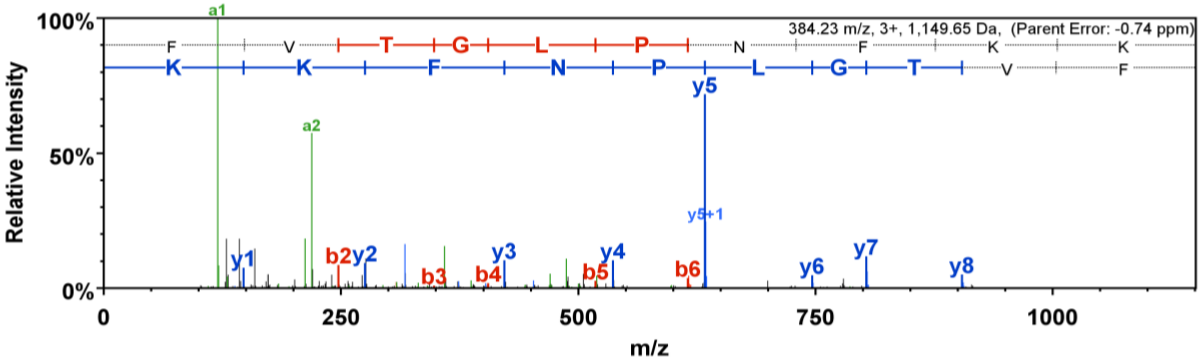


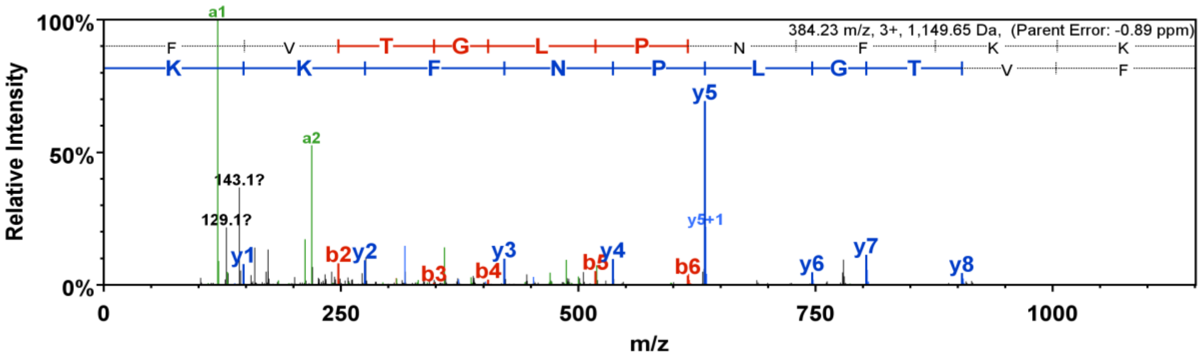

### No. 13. AtChr4@1344112@1344550: 2-oxoglutarate-dependent dioxygenase

**AtChr4@1344112@1344550 (100%), 16,507.6 Da**
**| potential novel**
**3 exclusive unique peptides, 5 exclusive unique spectra, 40 total spectra, 53/146 amino acids (36% coverage)**

```
ITFLHLCLKL TTAKHFDGTK ERRMGSCSPQ LPLICLSDQT LKPGSSKWVK
VRSDVRKALE DYGCFEAKID QVSMELQGSV LKAMQELFAL PTEAKQRNVC
PKPFTGYLSH NGLSESFGIK DANILEKAHE FTQQLWPEGN KSIRYM
```

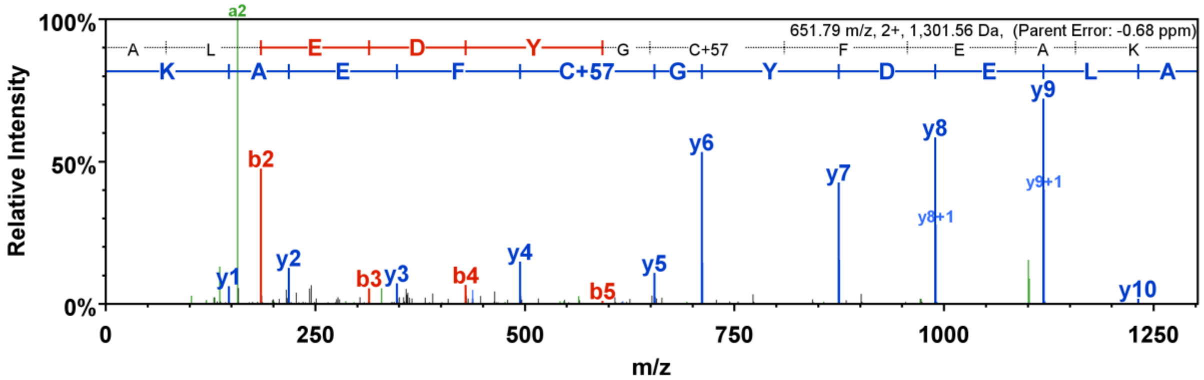


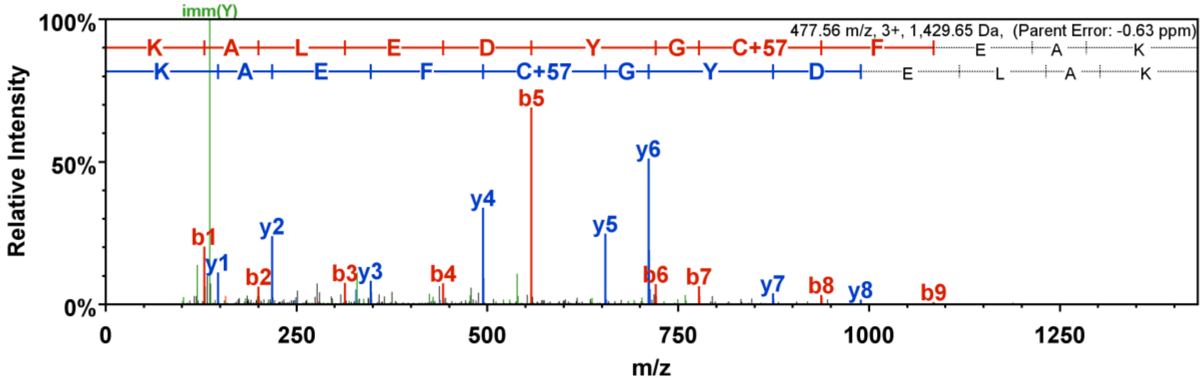


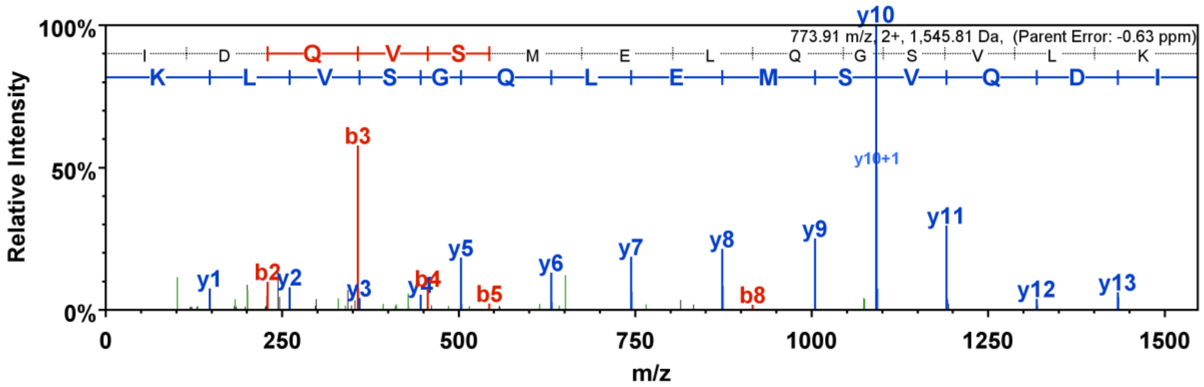

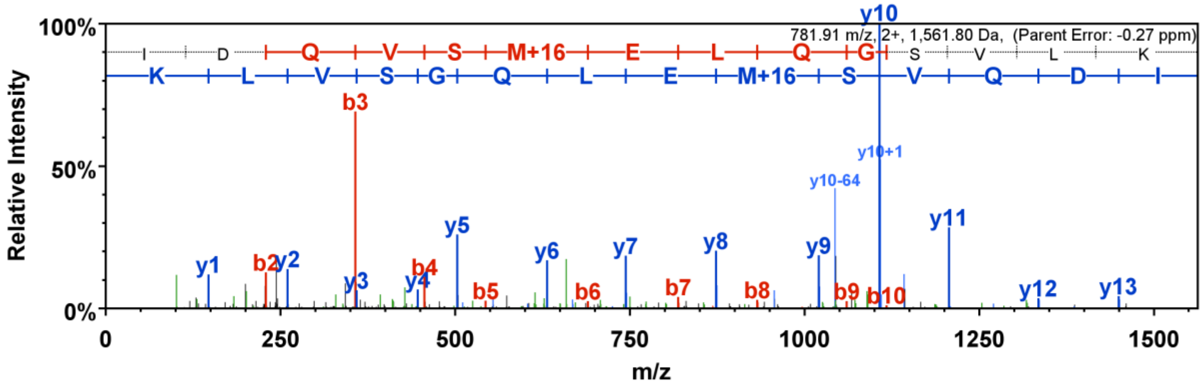


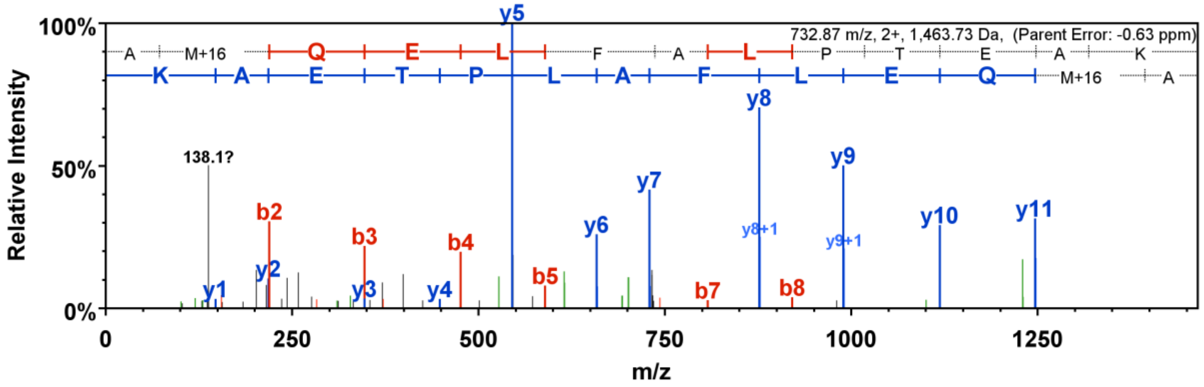


### No. 14. AtChr5@15331919@15332135: Alpha/beta-Hydrolases superfamily protein

**AtChr5@15331919@15332135 (100%), 8,143.7 Da**
**| potential novel**
**2 exclusive unique peptides, 2 exclusive unique spectra, 3 total spectra, 27/72 amino acids (38% coverage)**

KFFNLGTCIK PKWVMDFQDD EGLKETGR**FY** **EVEPVCIK**GV AHDMMLDCSW
EKGAK**VLLSW** **LCDLSKPGSL** **SA**

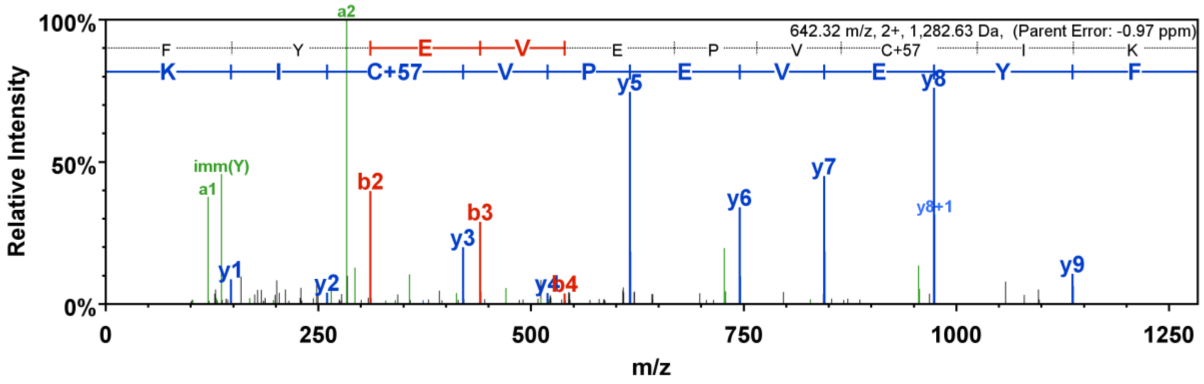
